# Examination of the enrichment of neuronal extracellular vesicles from cell conditioned media and human plasma using an anti-NCAM immunocapture bead approach

**DOI:** 10.1101/2025.05.13.653678

**Authors:** Mary E W Collier, Natalie Allcock, Nicolas Sylvius, Jordan Cassidy, Flaviano Giorgini

## Abstract

**Background:** The isolation of neuron-derived extracellular vesicles (nEVs) from biofluids offers the potential to discover novel biomarkers to aid in diagnosis and treatment of psychiatric and neurodegenerative diseases. A few studies have used anti-NCAM antibody-bead-based immunocapture to enrich nEVs from plasma, some with little method validation. We therefore examined in detail this method for nEV enrichment.

**Methods:** EVs were isolated from SH-SY5Y cell-conditioned media by precipitation, or from plasma using size exclusion chromatography. EVs were characterised using nanoparticle tracking analysis (NTA), transmission electron microscopy (TEM) and immunoblot analysis. SH-SY5Y-EVs were incubated with anti-NCAM immunocapture beads and examined by flow cytometry, immunoblot analysis and scanning electron microscopy (SEM). Immunocaptured plasma-derived EVs were examined using a sensitive NCAM ELISA, SEM and qPCR for miRNAs.

**Results:** Characterisation of SH-SY5Y-derived and plasma-derived EVs revealed the expected size distributions of EVs using NTA, the presence of EV markers using immunoblot analysis, and a cup-shaped morphology using TEM. Anti-NCAM beads, but not anti-L1CAM or IgG beads, captured NCAM-positive SH-SY5Y-EVs as shown by flow cytometry and immunoblot analysis. Both SH-SY5Y and plasma-derived EVs were visualised on the surface of anti-NCAM immunocapture beads using SEM. A sensitive NCAM ELISA detected NCAM antigen in plasma-derived EVs immunocaptured on anti-NCAM beads. qPCR analysis of plasma-derived EVs detected many miRNAs in total plasma-EVs with high expression of hsa-miR-16-5p, hsa-miR-451a and hsa-miR-126-3p. However, only between two and seven miRNAs were detected in EVs captured on anti-NCAM-beads from three blood donors. Finally, tissue distribution analysis of miRNAs from plasma-derived EVs on anti-NCAM beads revealed that these miRNAs are enriched in tissues or organs such as blood vessels, lung, bone, thyroid and heart, but were not enriched for brain-derived miRNAs. Discussion: This study indicates that anti-NCAM beads can efficiently enrich NCAM-positive EVs from cell culture conditioned media. However, nEV levels in small volumes of plasma are possibly too low to enable efficient anti-NCAM immunocapture for subsequent miRNA analysis. Other neuron-specific markers with high expression levels on nEVs are therefore required for processing patient samples where plasma volumes are likely to be low, and to allow efficient isolation of nEVs in clinical studies for subsequent cargo analysis.

## Introduction

Neurodegenerative diseases and severe psychiatric disorders have a major effect on the quality of life for patients but can be difficult to diagnose for effective early intervention and to provide suitable therapies. The discovery of biomarkers that indicate the state of the brain and central nervous system (CNS) in pathological conditions is therefore crucial to detect disorders as early as possible, to accurately diagnose patients, to monitor disease progression and to improve quality of life for patients with targeted therapies. Recently there has been increasing interest in the use of extracellular vesicles (EVs) as biomarkers of disease, particularly since EVs can be isolated by non-invasive methods from biofluids such as plasma and serum [1].

EVs are released from all cell types including cells in the brain such as neurons [2], microglia [3,4], astrocytes [5,6] and oligodendrocytes [7,8]. EVs released from brain cells, including neuron-derived EVs (nEVs), are thought to be able to cross the blood brain barrier and enter the peripheral blood circulation, particularly under inflammatory conditions when the blood brain barrier may be more permeable [9]. nEVs are potential biomarkers of neurological diseases because they contain nucleic acids and proteins derived from the neurons from which they were released. The miRNA and protein content of nEVs therefore reflects that of the parent cell and changes in the miRNA or protein content of nEVs from patients relative to healthy individuals, have the potential of giving insights into pathological processes in the brain and CNS which are not readily observed. In the last few years, antibody-bead immunocapture techniques have been developed to enrich nEVs from biofluids and take advantage of this “window into the brain” [10,11]. However, an important aspect of isolating nEVs from biofluids is the choice of neuronal marker used to target and enrich this specific population of EVs from biofluids, as well as thorough validation of the method used.

The adhesion molecules L1CAM (CD171) and NCAM1 (CD56) are expressed by neurons [2, 12] and have been previously used as targets to immunocapture and enrich nEVs from various biofluids including plasma, serum, cerebrospinal fluid (CSF), saliva and urine [10,11,13,14,15]. However, L1CAM is not strictly neuron-specific because it is expressed in other cell types [16,17,18,19,20] and a recent study has suggested that L1CAM is not associated with EVs but is predominantly in a soluble form in plasma and CSF [21]. NCAM has been detected on nEVs [22,23,24,25] and has been used in a small number of studies as a neuronal marker to enrich nEVs from plasma [14,15,26,27] and more recently, to enrich nEVs from brain tissue [23].

The aim of this study was to examine in detail the process of anti-NCAM immunocapture for nEV enrichment from cell culture conditioned medium and plasma and therefore validate the use of the neuronal adhesion molecule NCAM as a suitable target to enrich nEVs. Furthermore, unlike previous studies, we used a range of techniques, including scanning electron microscopy to directly visualise intact EVs on the surface of anti-NCAM immunocapture beads. Initially, EVs derived from the neuronal cell line SH-SY5Y were isolated from conditioned media and used to optimise the immunocapture technique using anti-NCAM immunocapture beads. The protocol was then applied to human plasma samples in an attempt to enrich for nEVs from plasma and profile the miRNA content of the anti-NCAM immunocaptured plasma-derived EVs in an attempt to confirm whether this approach can be used to efficiently isolate nEVs from plasma for subsequent miRNA expression analysis. We show that NCAM-positive EVs were immunocaptured from conditioned cell culture media and human plasma using anti-NCAM immunocapture beads. However, the small volumes of plasma (0.5 ml) used in this study, and typically used for clinical investigations, were too low to reliably measure miRNA levels in the immunocaptured nEVs, indicating that other more neuron-specific markers that are highly expressed on nEVs are required for efficient nEV isolation from plasma.

## Materials and Methods

### Cell culture and isolation of cell-derived EVs from conditioned media

The neuroblastoma cell line SH-SY5Y (ECACC 94030304) was cultured in DMEM containing 10 % (v/v) foetal calf serum (FCS) at 37°C under 5 % CO_2_. Prior to EV isolation, 7 x10^5^ cells were seeded into T75 flasks in DMEM containing 5 % (v/v) FCS and incubated for 72 h at 37°C. The cells were then incubated in medium containing 2 % (v/v) FCS for a further 24 h. Cells were washed three times with 10 ml of PBS, pH 7.4 and placed in 10 ml of serum free media (SFM) for 1 h at 37°C. The medium was then replaced with 10 ml of fresh SFM containing recombinant nerve growth factor (NGF) (50 ng/ml) (Thermo Fisher Scientific, Loughborough, UK) and incubated for 24 h at 37°C. The medium was then transferred to 50 ml tubes and supplemented with 1X cOmplete protease inhibitor cocktail (Sigma-Aldrich, Gillingham, UK) and EDTA to a final concentration of 0.5 mM. The medium was then centrifuged at 300 *g* for 10 min followed by 2000 *g* for 20 min at 4°C to remove cells and cell debris. ExoQuick-TC (System Biosciences, Palo Alto, CA, USA) was added to the media at a 1:5 ratio, gently mixed and incubated overnight at 4°C. The EVs were pelleted by centrifugation at 1500 *g* for 30 min at 4°C, followed by an additional 1500 *g* for 5 min. EVs were resuspended in 100 µl of 0.1 µm-filtered PBS containing 0.1 % (w/v) bovine serum albumin fraction V (BSA) (Sigma) for use in immunocapture assays. Alternatively, EVs were lysed in radioimmunoprecipitation assay (RIPA) buffer containing 1 % (v/v) Halt protease inhibitor cocktail (Thermo Fisher Scientific) for immunoblot analysis.

### Differentiation of SH-SY5Y cells

Cells were seeded into 6-well plates at a density of 0.75 x 10^5^ in DMEM supplemented with 5 % FCS and incubated at 37°C for 48 h. Cells were treated with 10 µM of all-trans retinoic acid (RA) (Sigma) in DMEM containing 5 % FCS and incubated for 48 h. Cells were treated with 10 µM RA in DMEM containing 2 % FCS and incubated for a further 48 h. Cells were then placed in SFM supplemented with 10 µM RA and 12.5 ng/ml brain-derived neurotrophic factor (BDNF) (Cambridge Biosciences, Cambridge, UK). The RA/BDNF-supplemented SFM was then replaced every 2-3 days up to day 11. On day 12, cells were washed twice with PBS and placed in SFM for 24 h. Cells were lysed in 150 µl of RIPA buffer containing 1 % (v/v) Halt protease inhibitor cocktail for immunoblot analysis. The conditioned media was pooled together (5 ml) and EVs isolated using ExoQuick-TC as described above.

### Blood collection and plasma preparation

Blood collection was approved by the University of Leicester Ethical Practices Committee (application 23825-fg36-Is) and all subjects provided written informed consent. Venous blood from healthy volunteers (non-fasting) was taken between 10 and 11 am with collection into 4.9 ml S-Monovette EDTA tubes (1.6 mg/ml) (Sarstedt, Nümbrecht, Germany) using Safety-Multifly 21G needles (Sarstedt) and centrifuged at 2500 *g* for 15 min at room temperature. The plasma was carefully removed avoiding the buffy coat and centrifuged again at 2500 *g* for 15 min at room temperature. Plasma (1 ml) was then aliquoted into screw-cap 1.5 ml tubes and stored at -80°C.

### Isolation of plasma-derived EVs

Plasma samples (1 ml) were defrosted at room temperature and supplemented with cOmplete protease inhibitor cocktail. Plasma was centrifuged at 1500 *g* for 10 min followed by 10,000 *g* for 10 min at room temperature. Exo-spin mini size exclusion chromatography columns (Cell Guidance Systems Ltd, Cambridge, UK) were equilibrated to room temperature, washed twice with 250 µl of 0.22 µm-filtered PBS and 100 µl of plasma added to the top of each column and allowed to enter the column. PBS (180 µl) was then added to the top of the column resulting in the elution of EVs into low protein-binding 1.5ml tubes. The Exo-spin columns were then washed four times with 200 µl of PBS and the process repeated until 0.5 ml of plasma had been processed. The EV fractions for each sample were pooled together and concentrated using Amicon Ultra 10 kDa molecular weight cut-off centrifugal filters (Sigma) by centrifugation at 3000 *g* for 45 min at 4°C. EVs were recovered by inverting the centrifugal filters and centrifuging at 1000 *g* for 2 min. EVs were supplemented with cOmplete protease inhibitor cocktail and BSA to a final concentration of 0.1 % for use in immunocapture assays.

### Antibody-bead immunocapture technique to immunocapture SH-SY5Y-derived EVs or plasma-derived EVs

Exosome-Streptavidin Isolation/Detection reagent Dynabeads (Thermo Fisher Scientific) (100 µl) were conjugated to 1 µg of biotinylated antibodies against L1CAM (clones 5G3 or UJ127; Thermo Fisher Scientific), NCAM (clone CMSSB; Thermo Fisher Scientific), CD81 (clone REA513; Miltenyi Biotec Ltd, Surrey, UK), calnexin (clone CANX/1543; Generon Ltd, Slough, UK), or mouse IgG (Thermo Fisher Scientific) in 100 µl PBS containing 0.1 % (w/v) BSA for 1 h at room temperature. Beads were then washed three times with 1 ml of PBS/BSA by placing the tubes on a PureProteome magnetic stand (Merck Millipore, Watford, UK) for 2 min followed by removal of the buffer. EVs in 100 µl PBS containing 0.1 % (w/v) BSA and cOmplete protease inhibitor cocktail were then added to the antibody-labelled beads and incubated at 4°C overnight with rotation. The beads were then washed twice with 500 µl of PBS containing 0.1 % (w/v) BSA, resuspended in 200 µl PBS and transferred to fresh 1.5 ml tubes. The PBS was removed and the EVs were lysed from the beads in 15 µl of RIPA buffer containing 1 % (v/v) Halt protease inhibitor cocktail (Thermo Fisher Scientific) and incubated on ice for 15 min. Loading buffer (15 µl) containing 10 % (v/v) beta mercaptoethanol was added to each sample and heated to 95°C for 5 min. Alternatively, EVs were lysed in suitable assay diluent containing 0.1 % (v/v) Triton X-100 for Meso Scale Discovery (MSD) assays. The beads were removed using the magnetic stand and the supernatant retained and stored at -80°C.

### Depletion of natural killer cell-derived EVs (NK-EVs) from plasma-derived EV samples

Before incubation of plasma-derived EVs with anti-NCAM or IgG immunocapture beads, EVs in PBS/0.1 % BSA (100 µl) containing cOmplete protease inhibitor were incubated with 20 µl of Dynabeads conjugated to 0.2 µg of biotinylated mouse anti-NKG2A antibody (131411; R&D Systems, Abingdon, UK) for 1 h at 4°C with rotation. This step was included to deplete NK-EVs and also acted as a clearing step to reduce non-specific binding. The non-bound EVs were then removed and the EVs were added to anti-NCAM or IgG-immunocapture beads as described above.

### SDS-PAGE and immunoblot analysis of EVs

Equal volumes of EV lysates were separated by 10 % (v/v) SDS-PAGE, transferred onto nitrocellulose membranes and blocked with Tris-buffered saline Tween-20 (TBST) (50 mM Tris-HCl, pH 8, 150 mM NaCl, 1 % (v/v) Tween-20) containing 5 % (w/v) low fat milk for 1 h. Membranes were incubated overnight at 4°C with the following primary antibodies diluted in TBST-milk; rabbit monoclonal anti-NCAM1 (Cell Signaling Technology, Leiden, The Netherlands; clone E7X9M; 1:1000 dilution), rabbit anti-L1CAM (Merck Millipore, 1:1000), mouse anti-CD81 (Thermo Fisher Scientific, 1:1500), rabbit anti-CD63 (R&D Systems, 1:1000), rabbit anti-Tsg101 (Sigma, 1:1000), rabbit anti-calnexin (Cell Signalling Technology; 1:1000), rat anti-alpha tubulin (Santa Cruz Biotechnology, Heidelberg, Germany, clone YOL1/34, 1:2500) or rabbit anti-ApoAI (R&D Systems; 1:2000). Membranes were washed three times for 5 min each with TBST and then incubated for 1 h at room temperature with an anti-rabbit-HRP conjugated secondary antibody (Vector Labs, Kirtlington, UK; 1 in 5000 dilution), an anti-mouse-HRP antibody (Vector Labs; 1 in 5000 dilution), or an anti-rat-HRP antibody (Vector Labs, 1 in 10,000 dilution). Membranes were washed four times with TBST and developed with Westar Supernova ECL detection reagent (Cyanagen, Bologna, Italy) and imaged using a G:BOX Chemi XX9 gel imaging system (Syngene, Cambridge, UK).

### Flow cytometry of EV-immunocapture beads

Exosome human CD81 Flow Detection Reagent Dynabeads (Thermo Fisher Scientific) (20 µl) or Exosome-Streptavidin Isolation/Detection reagent Dynabeads (20 µl) conjugated to 0.2 µg biotinylated antibodies were incubated with 100 µl of SH-SY5Y-derived EVs in PBS containing 0.1 % (w/v) BSA and 1x cOmplete protease inhibitor cocktail at 4°C overnight with rotation. Beads without antibody conjugation, beads conjugated to biotinylated anti-calnexin antibody, or beads conjugated to biotinylated mouse IgG were incubated with EVs as negative controls. The beads were washed twice with 500 µl of PBS/BSA and then resuspended in 220 µl of PBS/BSA. 100 µl of the beads/EV samples were transferred to 1.5 ml tubes and incubated with either 20 µl of mouse anti-CD81-PE antibody (clone JS-81; BD Biosciences, San Jose, CA, USA) or 20 µl of mouse IgG1k-PE matched isotype control (clone MOPC-21; BD Biosciences) for 1 h at room temperature. Alternatively, 100 µl of the beads/EVs were labelled with mouse anti-NCAM-PE (clone TULY56; 0.065 ug; Thermo Fisher Scientific) or mouse IgG-PE matched isotype control (clone P3.6.2.8.1; 0.065 ug, Thermo Fisher Scientific) for 1 h at room temperature. The beads were then washed twice with 500 µl of PBS/BSA, resuspended in 600 µl PBS/BSA and examined using a BD FACSCanto II flow cytometer (BD Biosciences). Beads were gated using forward and side scatter. A gate was set at 2 % of the control bead (no antibody conjugation) sample labelled with IgG-PE and data was analysed using Flow Jo software v10 (BD Biosciences).

### Nanoparticle Tracking Analysis

The size distributions and concentrations of SH-SY5Y-derived EVs and plasma-derived EVs were examined by Nanoparticle Tracking Analysis (NTA) using a Nanosight LM10 instrument (Malvern Panalytical Ltd, Malvern, UK). EVs from 20 ml of cleared SH-SY5Y conditioned media were resuspended in 200 µl of 0.1 µm filtered PBS and were diluted 1 in 5 in 1 ml of 0.1 µm-filtered PBS and loaded into the LM10 chamber. EVs isolated from plasma and concentrated using the Amicon centrifugal filters were diluted 1 in 84 in 0.1 µm-filtered PBS. Five 60 s videos with a camera level of 13 and a minimum of 150 completed tracks were obtained for each sample at room temperature using the NTA 2.2 software (Malvern Panalytical) and the temperature for each run was set. Videos were analysed using the NTA 2.2 software and the threshold was set to 10 with a gain of 1 and a minimal expected particle size of 100 nm.

### Transmission electron microscopy of EVs

EVs were resuspended in 0.1 µm-filtered PBS and fixed with 2 % (v/v) paraformaldehyde. SH-SY5Y-EVs were used neat whereas plasma-derived EVs were diluted 1 in 25 in PBS. EV samples (2.5 µl) were adsorbed onto freshly glow-discharged carbon film copper grids (Agar Scientific, Essex, UK) for 3 min, washed twice by transferring to 25 ml drops of PBS and stained with 1 % (w/v) uranyl acetate. EVs were visualised using a JEOL JEM1400 transmission electron microscope (JEOL UK Ltd, Welwyn Garden City, UK) with an accelerating voltage of 120 kV. Images were captured using an EMSIS Xarosa digital camera (EMSIS GmbH, Münster, Germany) with Radius software.

### Scanning electron microscopy analysis of EVs on captured on immunocapture beads

SH-SY5Y-derived EVs or plasma-derived EVs captured on immunocapture beads were washed twice with 0.5 ml of 0.22 µm-filtered PBS/0.1 % BSA and once with 0.5 ml of 0.1 µm-filtered PBS. Samples were then fixed with 2 % PFA. Bead suspension (20 µl) was adsorbed for 20 min onto poly-L-lysine treated glass slides in 24-well plates mounted on a 96 well plate magnetic bead mount. Samples were dehydrated through a series of 30, 50, 70, 90 and 2 x 100 % ethanol followed by 3:1, 1:1 & 1:3 ethanol:hexamethyldisilazane (HMDS). Samples were washed twice with 100 % HMDS over 15 min. Excess HMDS was removed, and samples were left to dry. Samples were mounted onto 13 mm aluminium stubs using carbon sticky tabs and coated with 7 nm of platinum using a Quorum Q150 TES coating unit (Quorum Technologies, Laughton, UK) Imaging Silver dag was applied to the edges of the glass to stabilise drift. EVs on beads were imaged using a Zeiss Gemini 360 FEG SEM (Carl Zeiss GmbH, Oberkochen, Germany) using SE2 detector at 1kV. Images were taken at 10,000- 50,000 x magnification. The size of the EVs were measured using ImageJ.

### Anti-NCAM-1 Mesoscale Discovery R-PLEX assays

R-PLEX single-plex assays for NCAM-1 (Meso Scale Discovery (MSD), Rockville, Maryland, USA) were used as sensitive electrochemiluminescence assays to quantify levels of NCAM in neuronally-enriched plasma-derived EV samples. EVs were lysed from the immunocapture beads in assay diluent supplemented with 0.1% (v/v) Triton X-100 and incubated on ice for 15 min. The beads were removed using the magnetic stand and supernatants were stored at -80°C until needed. Streptavidin-coated MSD GOLD 96 Small Spot SECTOR plates were coated with the provided biotinylated anti-NCAM capture antibody according to the manufacturer’s instructions. Plates were washed three times with 150 µl wash buffer and assay diluent (25 µl) and calibrator or diluted samples (25 µl) were added to the plates and incubated for 1 h at room temp on an orbital shaker at 750 rpm. Plates were washed three times with 150 µl wash buffer and incubated with the provided SULFO-TAG-conjugated detection antibody diluted to 1x in antibody buffer for 1 h at room temperature with shaking. Plates were washed three times with 150 µl wash buffer and developed by the addition of 150 µl of MSD GOLD Read Buffer B. The plates were read on a MESO QuickPlex SQ 120MM (MSD) and analysed using the Methodical Mind software (MSD).

### Quantitative PCR (qPCR) analysis of miRNAs in plasma-derived EVs

Total plasma-derived EVs eluted from the SEC columns (50 µl) or plasma-derived EVs immunocaptured on anti-NCAM or IgG immunocapture beads were lysed in 230 µl of lysis buffer C from the Maxwell RSC miRNA plasma and serum kit (Promega, Southampton, UK) containing 1 µl of the spike-in control mix (UniSp2, UniSp4 and UniSp5) (Qiagen, Manchester, UK) and incubated for 10 min on ice. The beads were then removed using a magnetic stand and the supernatant was retained and stored at -80°C. miRNA was isolated from the EV lysates using the Maxwell RSC miRNA from plasma and serum kit (Promega) on a Maxwell RSC instrument (Promega) according to the manufacturer’s instructions and eluted in 40 µl of nuclease free water. RNA (4 µl) was reverse transcribed using the miRCURY LNA RT kit (Qiagen) with the UniSp6 and cel-miR-39-3p spike-in controls included to monitor the efficiencies of the RT and PCR reactions. Samples were incubated at 42°C for 1 h, followed by inactivation of the RT enzyme at 95°C for 5 min. A master mix was prepared for each sample with 500 µl of miRCURY LNA miRNA SYBR Green master mix, 490 µl of water and 10 µl of cDNA. The master mix for each sample (10 µl) was pipetted into wells of the Human Serum/Plasma Small Focus (miScript) miRCURY LNA miRNA PCR panel (YAHS-206Z) (Qiagen). Real time quantification of the miRNAs was conducted on a LightCycler 480 thermocycler (Roche Diagnostics Limited, Burgess Hill, UK) with an initial activation at 95°C for 2 min followed by 45 cycles of 95°C for 10 s and 56°C for 1 min. Cp values were calculated using the LightCycler 480 1.5.0 SP3 software using absolute quantification three-point fit curves. Values for the noise band and the threshold for the fit-point curves were set for each miRNA on the plate and these settings were used for each miRNA for all samples to calculate Cp values.

## Results

### Characterisation of SH-SY5Y EVs

Initially, we examined whether non-differentiated or differentiated SH-SY5Y cells would be the best model for the isolation of neuronal EVs, since these cells can be differentiated to a more neuronal phenotype. White light microscopy images showed that treatment of cells with 10 µM RA for 11 days together with 12.5 ng/ml BDNF for the final six days resulted in a more neuronal morphology with the formation of neurites (Supplemental Data File 1 Figure 1A). Immunoblot analysis revealed increased expression of the neuronal marker synaptophysin in differentiated cells (Supplemental Data File 1 Figure 1B) and reduced levels of ID2 which is downregulated during differentiation (Supplemental Data File 1 Figure 1C). However, NTA showed reduced levels of EVs were released from differentiated cells compared to non-differentiated cells (Supplemental Data File 1 Figure 1D). Therefore, in this study we isolated EVs from non-differentiated SH-SY5Y cells.

**Figure 1.**
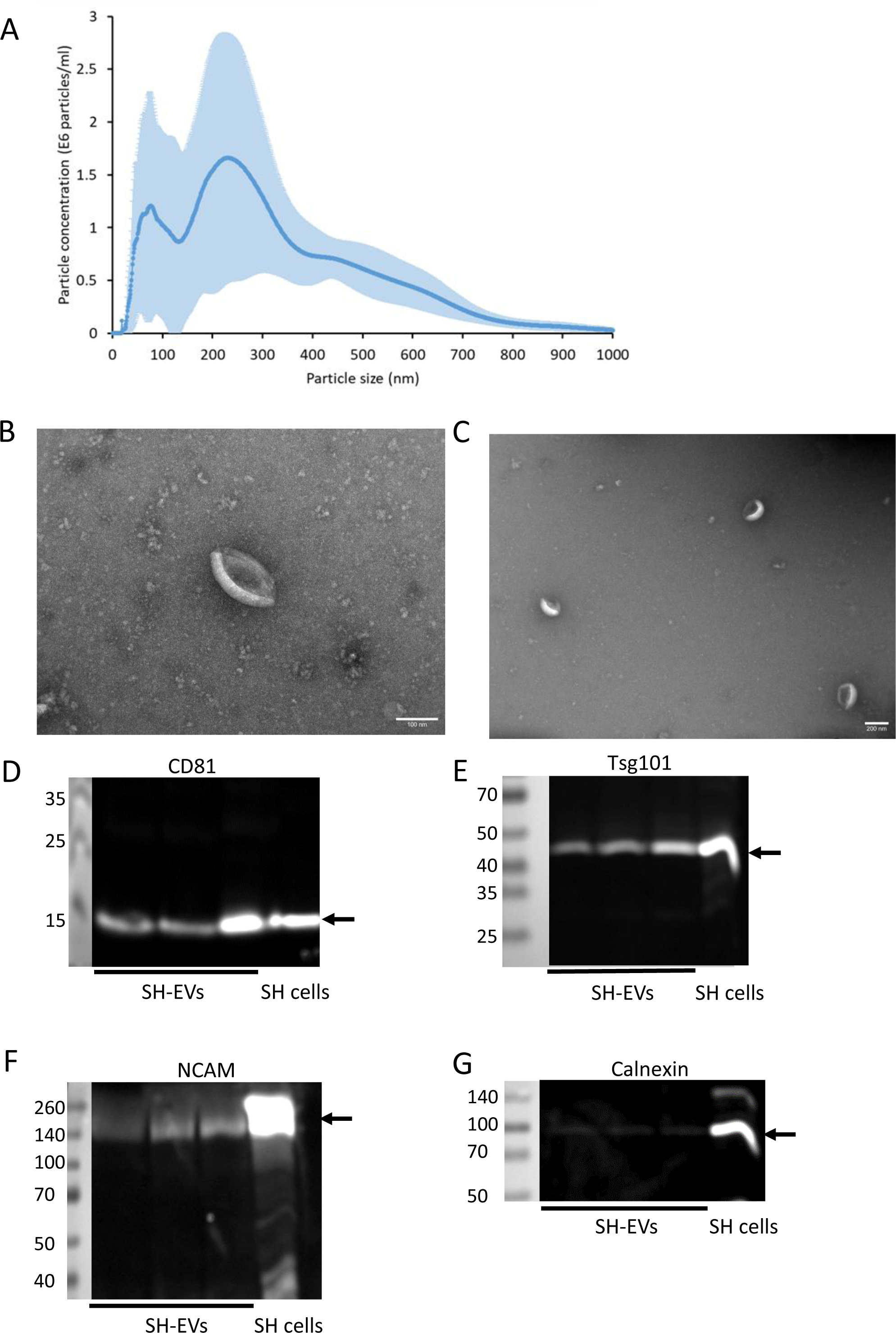
Characterisation of SH-SY5Y cell-derived EVs. SH-SY5Y cells were incubated in SFM supplemented with NGF (50 ng/ml) for 24 h and EVs were isolated from the cleared media using ExoQuick-TC. A) The size distribution of EVs was analysed by NTA (n=4 ± SD). B&C) Negative staining of EVs and TEM analysis revealed the presence of EVs with a cup-shaped morphology. Scale bar = 100 nm (B) and 200 nm (C). D-G) EVs were lysed in RIPA buffer and the presence of CD81 (D), Tsg101 (E), NCAM (F) and calnexin (G) was analysed by immunoblot analysis (data shows SH-EVs isolated from three T75 flasks of SH-SY5Y cells).

The size distributions of EVs isolated from non-differentiated SH-SY5Y cell conditioned media (SH-EVs) were examined using NTA and revealed that SH-EVs had a mode diameter of 225 ± 30 nm with two main populations of EVs; smaller EVs with a peak size of 80 nm and larger EVs with a peak size of 230 nm (Figure 1A). Negative staining and transmission electron microscopy (TEM) analysis of SH-EVs revealed the presence of vesicles with a cup-shaped morphology typical of EVs observed by this approach (Figure 1B&C). Finally, immunoblot analysis of SH-EVs confirmed the presence of the EV markers CD81 (Figure 1D) and Tsg101 (Figure 1E). Immunoblot analysis of neuronal adhesion molecules revealed that full length NCAM (140 kDa) was present in the SH-EVs (Figure 1F). Although full length L1CAM (200-220 kDa) was present in SH-SY5Y cell lysate, the majority of L1CAM in SH-EVs was in the form of a 130 kDa L1CAM fragment (Supplemental Data File 1 Figure 2). Very little of the endoplasmic reticulum marker calnexin was detected in the EV samples, indicating that the EV preparations were mostly free of contaminating cell debris (Figure 1G).

**Figure 2.**
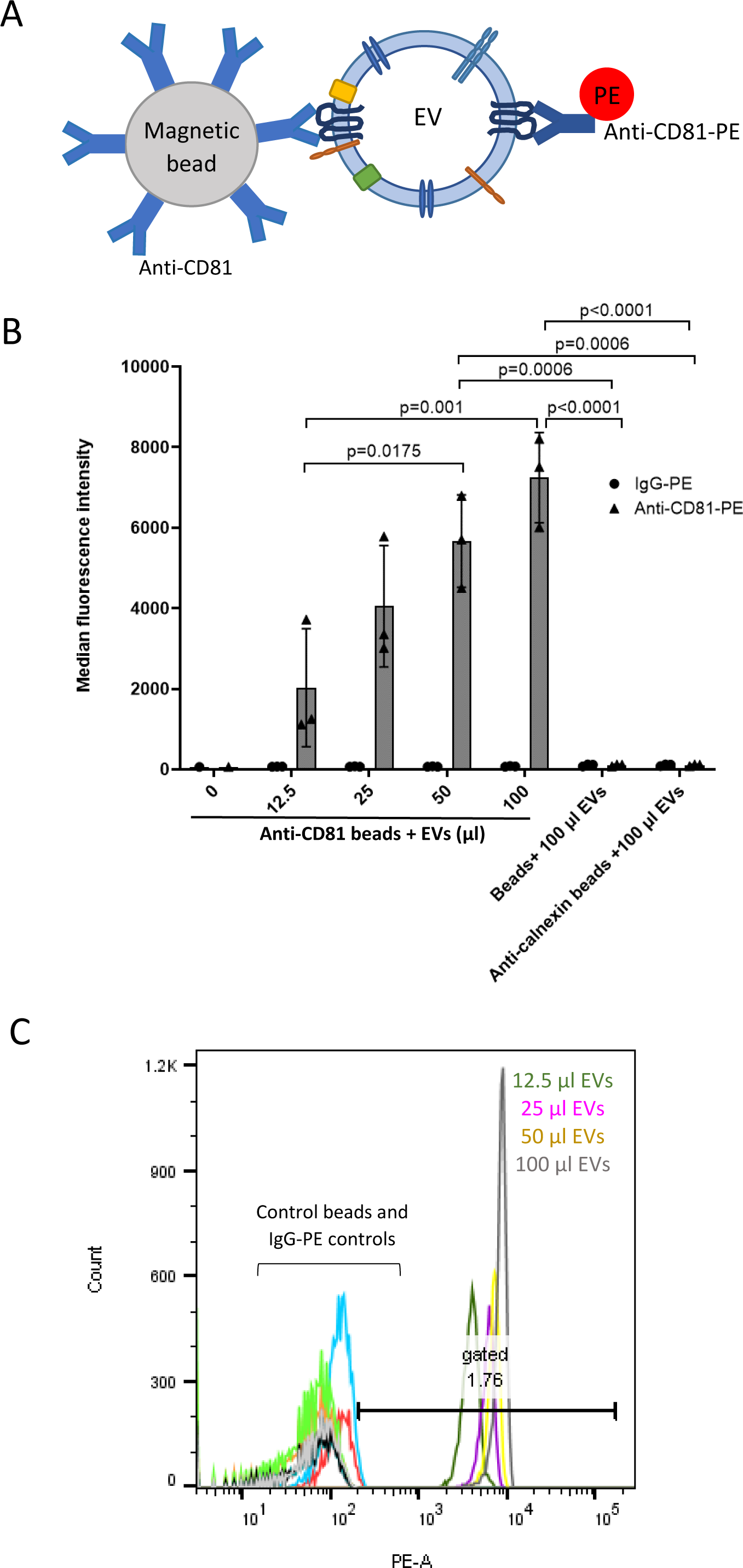
Flow cytometry of CD81-positive SH-SY5Y EVs immunocaptured using pre-conjugated anti-CD81 Dynabeads. A) Schematic representation of an EV captured by an anti-CD81-conjugated immunocapture bead and labelled with anti-CD81-PE. B) Increasing volumes (12.5-100 µl) of SH-SY5Y cell-derived EVs were incubated overnight with pre-conjugated anti-CD81 Dynabeads and then labelled with anti-CD81-PE or IgG-PE antibodies. Negative control samples included unlabelled beads or anti-calnexin conjugated magnetic beads incubated with 100 µl of SH-SY5Y EVs. Samples were examined by flow cytometry and median fluorescence intensities were plotted (n=3 ± SD, One Way ANOVA). C) Histogram plots showing anti-CD81-PE fluorescence intensities against bead count. Histograms for control beads (beads without conjugated antibody and anti-calnexin beads) and IgG-PE labelled samples are shown on the left side of the plot. Histograms for anti-CD81-PE labelled samples are shown on the right side of the plot: grey histogram = 100 µl of EVs, yellow histogram = 50 µl of EVs, purple histogram = 25 µl pf EVs and dark green histogram = 12.5 µl of EVs.

### The immunocapture of CD81-positive SH-SY5Y-derived EVs by anti-CD81-immunocapture beads optimised using flow cytometry

Initially the volume of SH-EVs required for detection by flow cytometry was optimised using anti-CD81 immunocapture beads. Exosome human CD81 flow detection reagent consists of Dynabeads pre-conjugated to an anti-CD81 antibody and was also initially used as a proof of concept to show that specific populations of EVs could be isolated using magnetic beads conjugated to antibodies specific for antigens present on the surface of EVs. Increasing volumes of SH-EVs (12.5-100 µl) were incubated with pre-conjugated anti-CD81 beads and then labelled with an anti-CD81-PE conjugated antibody or control IgG-PE antibody and examined by flow cytometry (Figure 2A). Median fluorescence intensity (MFI) values significantly and dose-dependently increased as the amount of SH-EVs incubated with the anti-CD81 beads doubled, indicating that the anti-CD81 beads captured CD81-positive EVs (Figure 2B&C). However, incubation of the anti-CD81 beads with 100 µl of EVs appeared to start saturating the binding sites on the beads, resulting in lower MFI values than expected. The binding of EVs to the beads was confirmed to be specific as no significant increase in fluorescence was observed in samples where the maximum volume of EVs (100 µl) was incubated with beads alone or with beads conjugated to an anti-calnexin antibody (Figure 2B). These data indicate that incubation of 100 µl of SH-EVs with antibody-conjugated immunocapture beads was optimal for detection by flow cytometry.

### NCAM-positive SH-SY5Y-derived EVs were detected on anti-NCAM immunocapture beads but not anti-L1CAM beads using flow cytometry

Initially, the expression of NCAM on the surface of SH-SY5Y cells was confirmed by flow cytometry (Supplemental Data File 1 Figure 3). To investigate whether SH-EVs can be isolated based on the presence of neuronal markers, Exosome Streptavidin isolation/ detection reagent magnetic beads were conjugated to a biotinylated antibody against NCAM (clone CMSBB) which recognises NCAM on the surface of non-fixed, non-permeabilised cells [28,29], or biotinylated antibodies against L1CAM (clones 5G3 and UJ127) which bind to the extracellular domain of L1CAM. The beads were then incubated with 100 µl of SH-EVs and labelled with either an anti-CD81-PE antibody or IgG-PE and examined by flow cytometry (Figure 3A). Beads were gated using forward and side scatter as shown in Figure 3B. Incubation of SH-EVs with anti-NCAM beads followed by labelling with anti-CD81-PE resulted in significant increases in MFI values compared to SH-EV samples incubated with IgG-conjugated beads (p=0.0062) or control beads without antibody (p=0.0062) (Figures 3C-E), indicating that NCAM-positive EVs expressing CD81 were immunocaptured on anti-NCAM beads. As expected, high levels of CD81-positive SH-EVs were captured by anti-CD81 beads (Supplemental Data File 1 Figure 4A), as shown by the high MFI values for these samples (MFI for anti-CD81 beads + anti-CD81-PE = 4865 ± 1238, MFI for anti-CD81 beads + IgG-PE = 111 ± 14, n=4). Neither the anti-L1CAM antibody 5G3, which is raised against the N-terminus of L1CAM, nor the anti-L1CAM antibody UJ127, antibody which targets the fourth fibronectin domain within the extracellular domain of L1CAM, immunocaptured CD81-positive SH-EVs (Figure 3C, Supplemental Data File 1 Figures 4B&C) since the MFI values for these samples were similar to those of the negative control samples (beads without antibody, IgG-conjugated beads and anti-calnexin conjugated beads) (Figures 4C&E, Supplemental Data File 1 Figure 4D).

**Figure 3.**
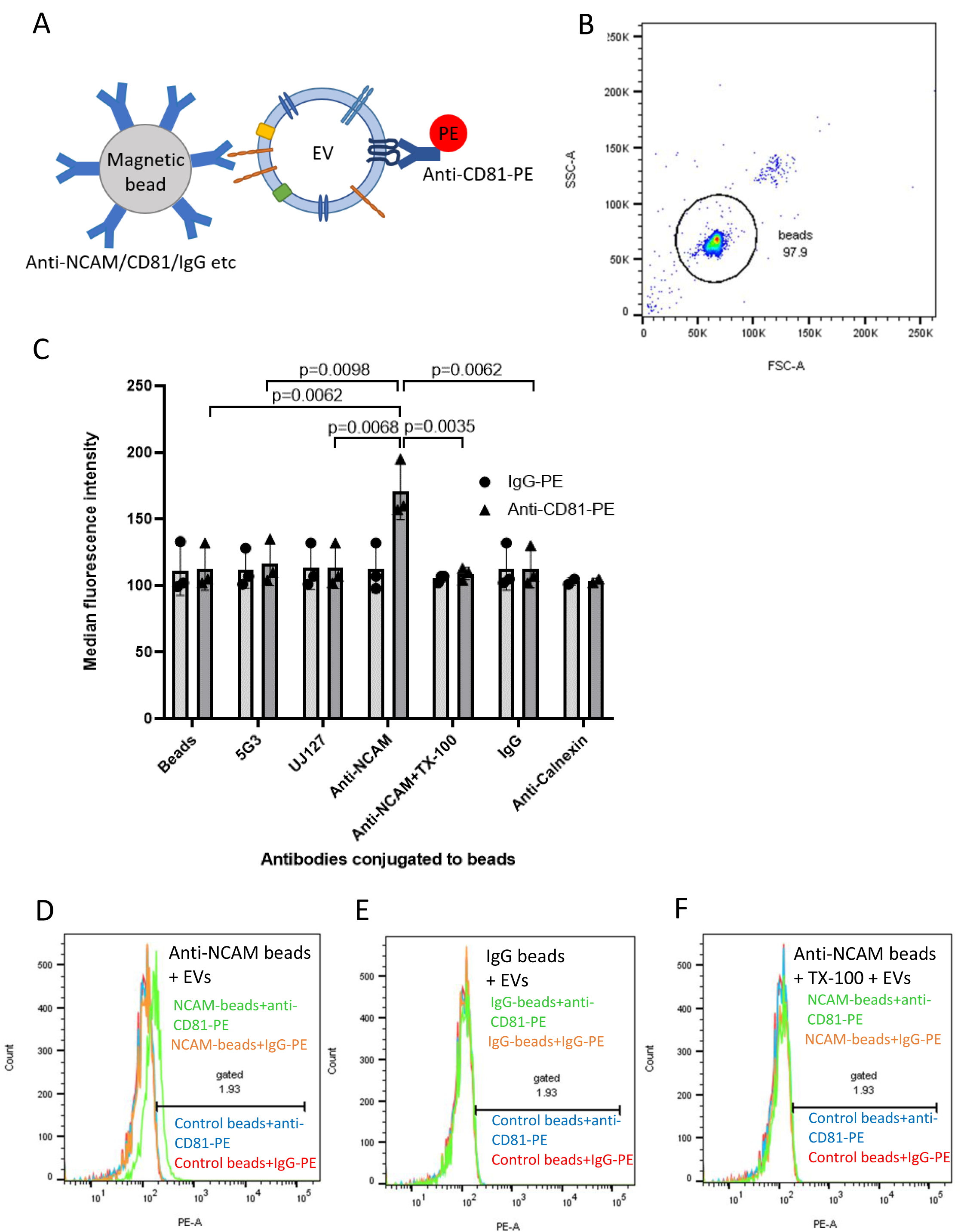
Flow cytometry of CD81-positive SH-SY5Y EVs immunocaptured using anti-NCAM or anti-L1CAM Dynabeads. A) Schematic representation of an EV captured by an anti-NCAM/CD81/IgG-conjugated immunocapture bead and labelled with anti-CD81-PE. B) Forward and side scatter plot of Dynabeads. C) SH-SY5Y cell-derived EVs (100 µl) were incubated overnight with antibody-conjugated beads and then labelled with anti-CD81-PE or IgG-PE antibodies and analysed by flow cytometry (n=3 ± SD, One Way ANOVA). Histograms are shown for anti-NCAM beads (D), IgG beads (E) or anti-NCAM beads washed with 0.25 % TX-100 (F) and then labelled with anti-CD81-PE (green histograms) or IgG-PE (orange histograms) compared to control beads (without conjugated antibody) labelled with anti-CD81-PE (blue histograms) or IgG-PE (red histograms).

**Figure 4.**
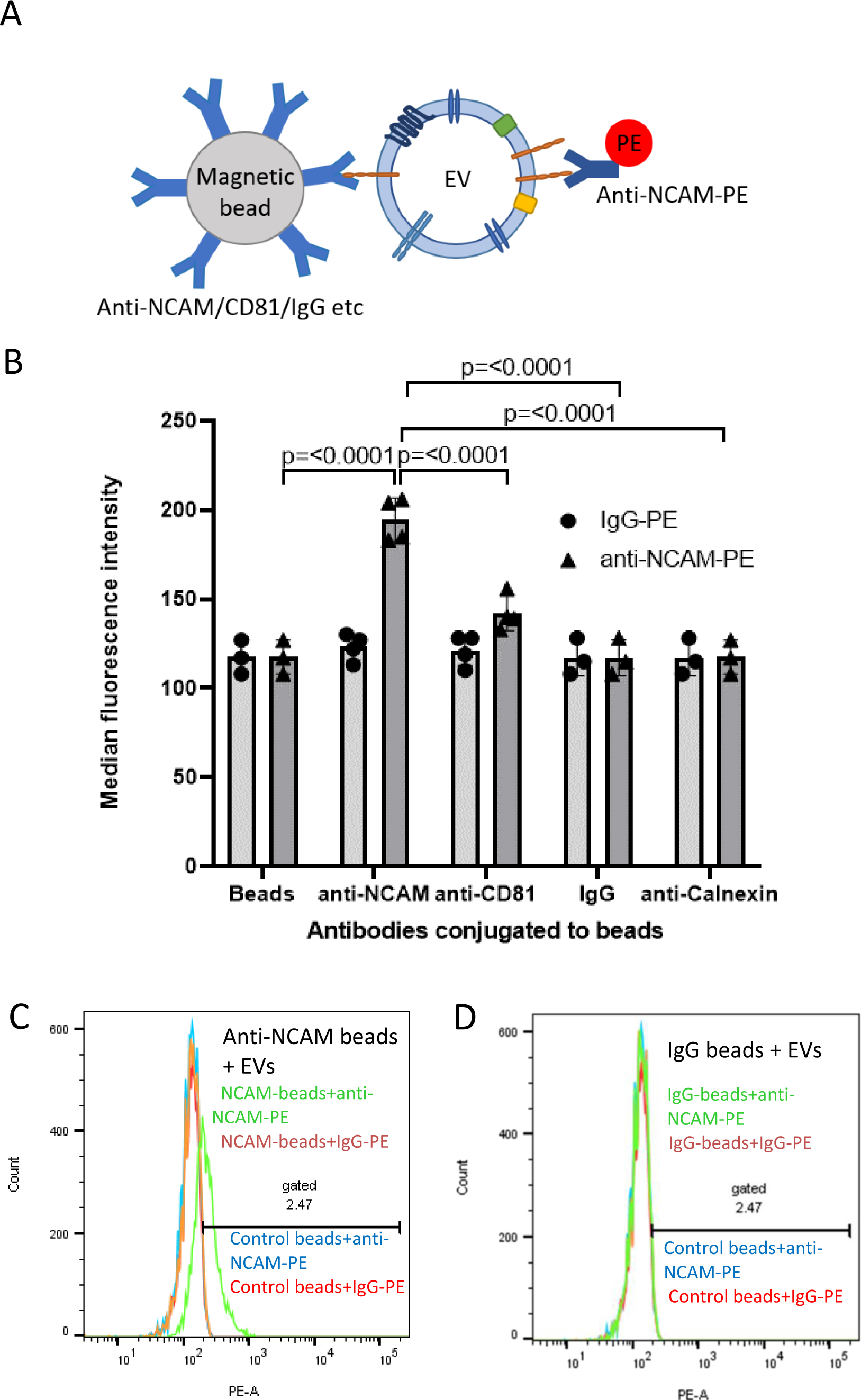
Flow cytometry of NCAM-positive SH-SY5Y EVs immunocaptured using anti-NCAM Dynabeads. A) Schematic representation of an EV captured by an anti-NCAM/CD81/IgG-conjugated immunocapture bead and labelled with anti-NCAM-PE. B) SH-SY5Y cell-derived EVs (100 µl) were incubated overnight with antibody-conjugated Dynabeads and then labelled with anti-NCAM-PE or IgG-PE antibodies and analysed by flow cytometry (n=3/4 ± SD, One Way ANOVA). Histograms of anti-NCAM Dynabeads (C) or IgG Dynabeads (D) labelled with anti-NCAM-PE (green histogram) or IgG-PE (orange histogram) compared to control beads (without conjugated antibody) labelled with anti-NCAM-PE (blue histogram) or IgG-PE (red histogram).

To confirm that intact EVs were captured by the immunocapture beads, some sets of anti-NCAM beads were incubated with SH-EVs and then washed with distilled water containing 0.25 % (v/v) Triton X-100 immediately after EV immunocapture in order to lyse the EVs [30,31] without disrupting protein-antibody interactions [32]. The beads were then washed and labelled with the anti-CD81-PE antibody and analysed by flow cytometry. Solubilisation of the lipids using Triton X-100 resulted in a significant reduction in MFI values of these samples compared to EVs immunocaptured on anti-NCAM beads without Triton X-100 treatment (p=0.0035) (Figures 3C&F). This indicated that the EVs captured by the anti-NCAM beads were intact and that the anti-CD81-PE antibody recognised CD81 on NCAM-positive lipid vesicles bound to the beads.

The presence of NCAM on EVs immunocaptured by anti-NCAM beads was also examined by labelling captured EVs with an anti-NCAM-PE antibody (clone TULY56 which does not block CMSSB binding) followed by flow cytometry (Figure 4A). A significant increase in MFI values was observed for anti-NCAM beads incubated with SH-EVs and labelled with an anti-NCAM-PE antibody compared to EVs incubated with anti-calnexin beads (p<0.0001), IgG beads (p<0.0001), or control beads without antibody (p<0.0001) (Figures 4B-D), further confirming the immunocapture of NCAM-positive EVs by anti-NCAM capture immunocapture beads.

### NCAM-positive SH-SY5Y-derived EVs containing the EV marker Tsg101 were detected on anti-NCAM immunocapture beads using immunoblot analysis

Since flow cytometric analysis only provided information on surface markers on the immunocaptured EVs, the experiment was repeated but the EVs immunocaptured on the beads were lysed and examined by SDS-PAGE and immunoblot analysis. This revealed the presence of the EV marker Tsg101 and full length NCAM in SH-EVs immunocaptured using anti-NCAM beads, confirming the immunocapture of intact NCAM-positive EVs by anti-NCAM beads (Figure 5A). Anti-CD81 beads were used as a positive control for EV immunocapture and showed high levels of Tsg101 in the EVs, low levels of NCAM and no detectable L1CAM. Although the 130 kDa L1CAM fragment was detected in the total EV samples, L1CAM was not detected in any of the immunocaptured EV samples, even those incubated with beads conjugated to the anti-L1CAM antibodies 5G3 or UJ127 (Figure 5A). Tsg101 was not detected in samples incubated with anti-L1CAM antibody-conjugated beads, control beads without antibody conjugation, or with the anti-calnexin beads, further indicating that anti-L1CAM beads did not immunocapture SH-SY5Y-EVs. Importantly, calnexin was not observed in any of the immunocaptured samples, indicating that intact EVs, rather than cell debris were present on the beads (Figure 5A).

**Figure 5.**
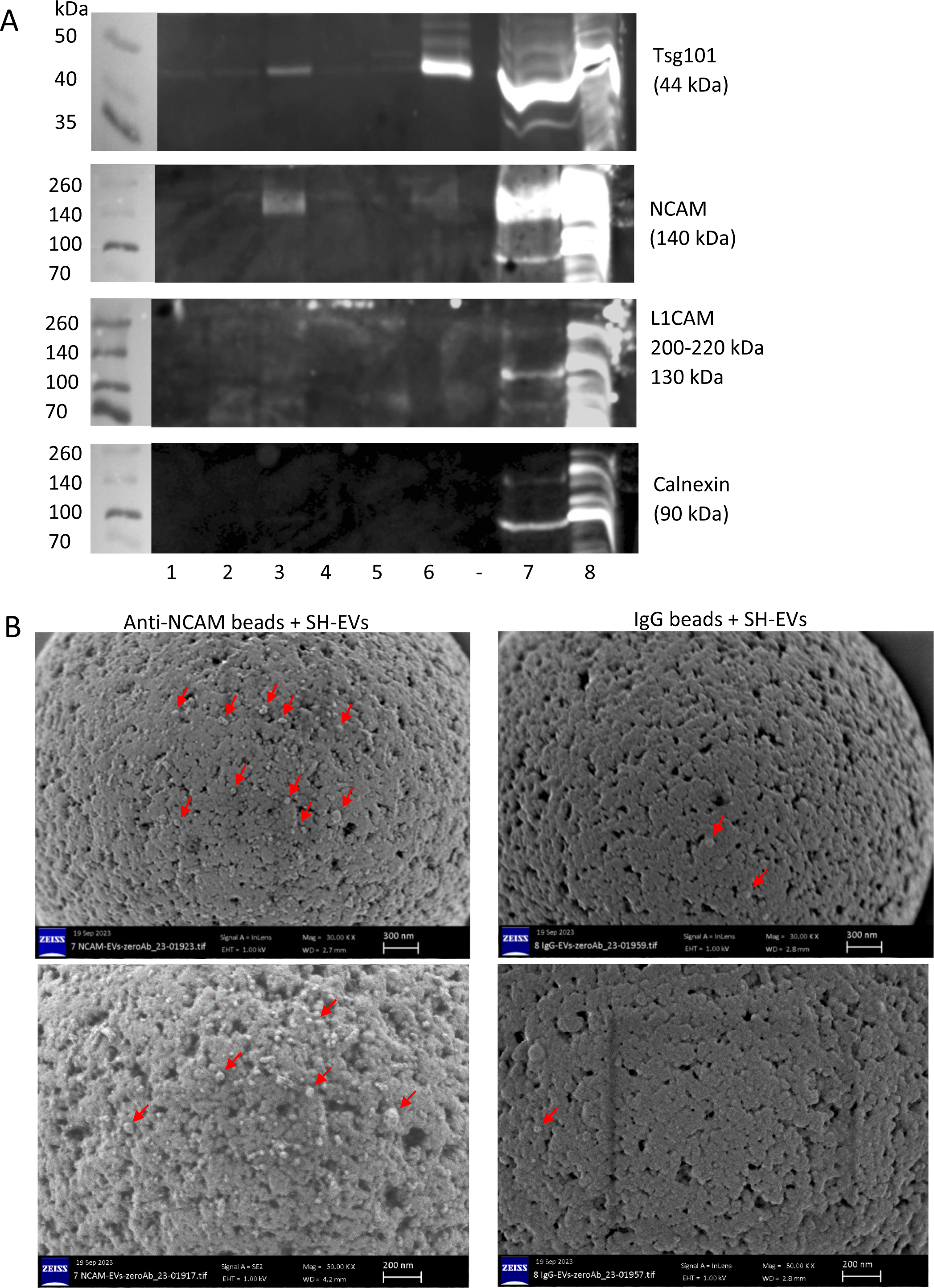
Immunoblot and SEM analysis of SH-SY5Y EVs immunocaptured on anti-NCAM beads. A) SH-SY5Y cell-derived EVs (100 µl) were incubated overnight with antibody-conjugated beads. The beads were then washed and the immunocaptured EVs lysed in RIPA buffer. Proteins were separated by SDS-PAGE and analysed by immunoblot for Tsg101, NCAM, L1CAM and calnexin. The numbers below the blots indicate the antibodies conjugated to the beads as follows; 1= anti-L1CAM clone 5G3, 2 = anti-L1CAM clone UJ127, 3 = anti-NCAM clone CMSSB, 4 = beads without antibody, 5 = anti-calnexin, 6 = anti-CD81, 7 = SH-SY5Y EVs, 8 = SH-SY5Y cell lysate. Gels are representative of three experiments. B) SH-EVs were incubated with anti-NCAM or IgG immunocapture beads overnight. Beads were washed to remove non-bound EVs, and beads were fixed in 2 % PFA and imaged by SEM. Arrows indicate vesicle structures on the surface of the beads. Images shown are at x30,000 (upper images) and x50,000 magnification (lower images).

### SH-SY5Y-derived EVs were visualised on anti-NCAM immunocapture beads using scanning electron microscopy (SEM)

SEM analysis of anti-NCAM immunocapture beads following incubation with SH-EVs was used to examine whether vesicles were present on the surface of the beads and further validate this method for nEV immunocapture. SEM analysis of beads alone without incubation with EVs revealed that the beads had the expected size of 4.5 µm (Supplemental Data File 1 Figures 5A&B). Anti-NCAM beads incubated with SH-EVs appeared to be decorated with vesicles, whereas very few SH-EVs were observed on IgG-beads (Figure 5B and Supplemental Data File 1 Figures 5C-J). In some SEM images, non-vesicular material was visible which may be protein aggregates (Supplemental Data File 1 Figure 5K). This confirmed that intact EVs were specifically captured on the surface of the anti-NCAM immunocapture beads.

### Characterisation of plasma-derived EVs

NTA of EVs isolated from the plasma of three healthy donors using Exo-spin columns and concentrated using Amicon centrifugal filters revealed a mode diameter of 144 nm ± 12.6 nm (Figure 6A). TEM analysis of plasma-derived EVs showed the presence of cup-shaped EVs (Figure 6B). The presence of EV markers in plasma-derived EVs was examined by immunoblot analysis and confirmed the presence of EV markers CD81 (Figure 6C), CD63 (Figure 6D) and Tsg101 (Figure 6E), as well as the cytoplasmic protein alpha tubulin (Figure 6F). Both Tsg101 and alpha tubulin in plasma-derived EVs had higher molecular weights than expected which may be due to post-translational modification of these proteins within plasma-derived EVs [33]. Calnexin was not detected in any of the plasma-EV samples indicating lack of contamination of the EV samples with cellular debris (Figure 6G). However, ApoAI was observed in plasma EV samples, suggesting carry-over of lipoproteins during size exclusion chromatography (Figure 6H). NCAM was not detectable in plasma-EVs by immunoblot analysis (Supplemental Data File 1 Figure 6).

**Figure 6.**
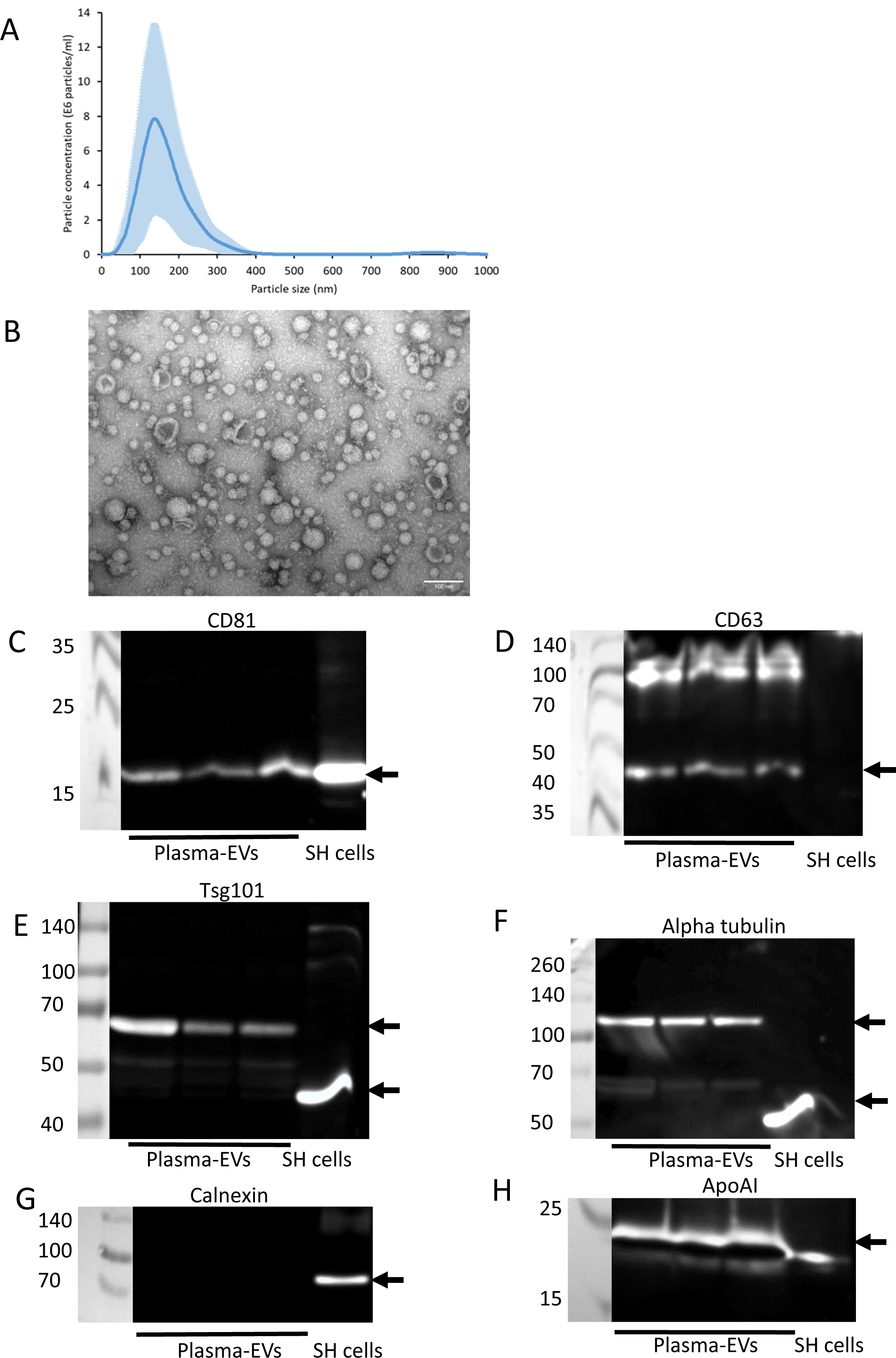
Characterisation of plasma-derived EVs. EVs were isolated from 0.5 ml of plasma using Exo-spin size exclusion chromatography columns and concentrated using 10 kDa MWCO centrifugal filters. A) The size distribution of the EVs was analysed by NTA (average data from three donors ± SD). B) TEM of negatively stained plasma-derived EVs revealed the presence of EVs with a cup-shaped morphology (scale bar = 100 nm). EVs from three donors were lysed in RIPA buffer and analysed by SDS-PAGE and immunoblot analysis for CD81 (C), CD63 (D), Tsg101 (E), alpha-tubulin (F), calnexin (G) and ApoAI (H).

### NCAM-positive plasma-derived EVs were immunocaptured on anti-NCAM immunocapture beads

Next, the immunocapture of NCAM-positive EVs from plasma-derived EVs was examined. NCAM antigen was detected in samples lysed from anti-NCAM beads incubated with plasma-derived EVs but not IgG-beads (Figure 7A). Total EVs isolated using SEC columns and concentrated using Amicon centrifugal filters were used as a positive control for NCAM expression in plasma-derived EVs (Supplemental Data File 1 Figure 7). Six samples (20 µl of anti-NKG2A beads conjugated to 0.2 µg antibody each) from the NK-EV depletion step were pooled together and lysed, but NCAM was not detected in these samples (Figure 7A). We also attempted to measure levels of the EV marker CD9 on EVs captured on anti-NCAM immunocapture beads using a CD9 ELISA, but measurements were below the limit of detection of the assay.

**Figure 7.**
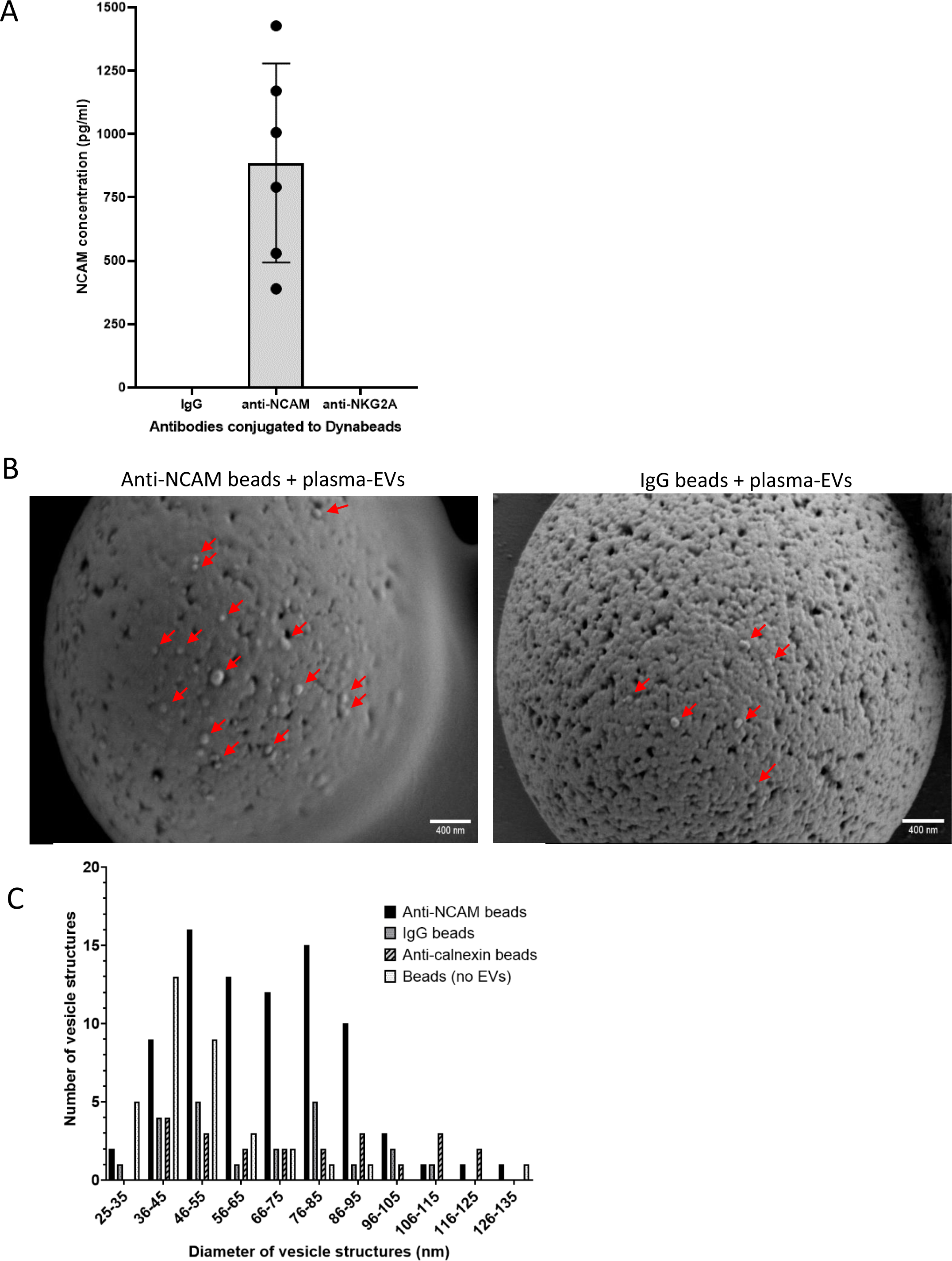
Verification of the immunocapture of NCAM-positive plasma-derived EVs using anti-NCAM immunocapture beads. EVs were isolated from plasma by size exclusion chromatography and depleted of NK-derived EVs using anti-NKG2A-conjugated beads. EVs were then incubated with anti-NCAM or IgG beads. A) EVs were lysed from the anti-NCAM, anti-NKG2A and IgG beads and levels of NCAM were measured using the MSD electrochemiluminescence NCAM assay (data from six healthy donors ± SD). B) SEM images of plasma-derived EVs immunocaptured on anti-NCAM beads or control IgG beads. C) Size distributions of plasma-derived EVs captured on anti-NCAM beads, IgG beads or anti-calnexin beads. Beads not incubated with EVs were used as a negative control. The diameters of vesicle structures on beads from ten images at x25,000 magnification for anti-NCAM beads, IgG beads and beads without EVs, and nine images from anti-calnexin beads were measured using ImageJ.

The presence of EVs on anti-NCAM beads incubated with plasma-derived EVs from two donors was further examined using SEM and revealed vesicle structures on anti-NCAM beads but also some on IgG beads and anti-calnexin beads, indicating some non-specific binding (Figure 7B and Supplemental Data File 2). The smaller average diameters of EVs estimated from SEM images compared to EV diameters measured using NTA may be due to dehydration and shrinkage of the vesicles during processing for SEM. As an additional control, donor 2 plasma-derived EVs were incubated with anti-calnexin conjugated-beads and showed low numbers of vesicle structures on the surface (Supplemental Data File 2). A few images of beads alone without conjugation to antibodies or incubation with EVs also showed the presence of spherical structures (Supplemental Data File 2). The diameters of spherical structures on the anti-NCAM and control immunocapture beads as well as beads not incubated with EVs were measured using ImageJ. This revealed that all beads, even those not incubated with EVs, had spherical structures of 35-55 nm on the surface (Figure 7C). However, anti-NCAM beads additionally had vesicle structures which ranged from of 56-95 nm in diameter which were not present on any of the control bead samples and indicate the presence of intact plasma-derived EVs on the surface of anti-NCAM immunocapture beads (Figure 7C).

### miRNAs were readily detected in total plasma-derived EVs but few miRNAs were detected in plasma EVs immunocaptured on anti-NCAM beads

Finally, the potential to measure the expression of miRNAs in immunocaptured nEVs was assessed using qPCR. Initially the expression of hsa-miR-16-5p, which is highly and stably expressed in plasma [34], was examined in plasma-derived EVs immunocaptured on anti-NCAM or IgG beads as well as in total plasma-derived EVs. Low levels of hsa-miR-16-5p were detected in EVs immunocaptured on anti-NCAM beads with Cp values ranging from 35.5-39.2 cycles (Figure 8A). However, two samples incubated with IgG beads had Cp values of >38 cycles, indicating there was non-specific binding of miRNAs or EVs to the beads, resulting in low levels of amplification. Total plasma-derived EVs from the three healthy donors had higher levels of hsa-miR-16-5p with Cp values ranging from 26.4-27.7 cycles (Figure 8A). The UniSp6 spike-in control showed good amplification in all samples, confirming efficient reverse transcription and PCR (Supplemental Data File 1 Figure 8).

**Figure 8.**
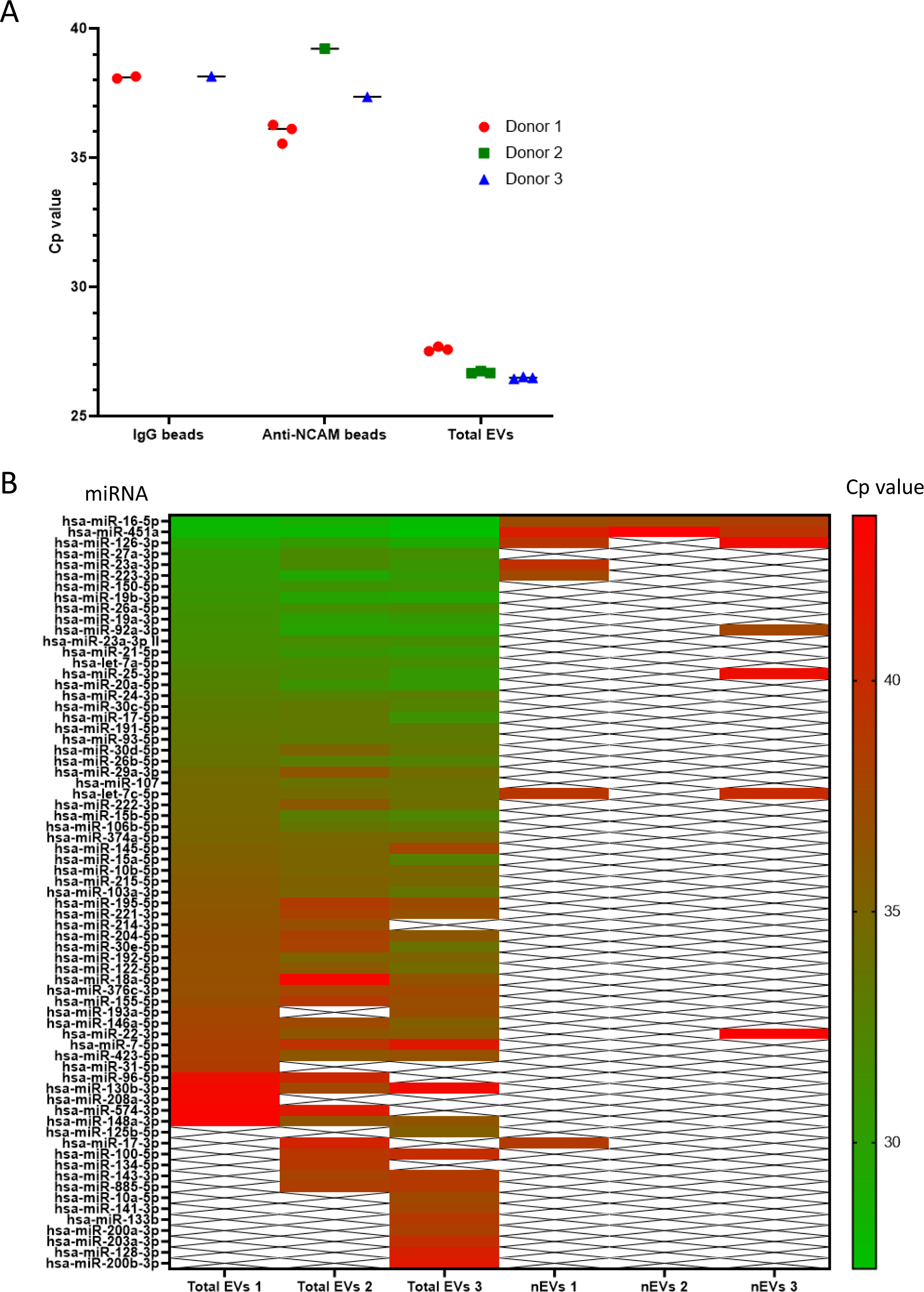
Analysis of the miRNA content of plasma-derived EVs immunocaptured on anti-NCAM immunocapture beads. Total plasma-derived EVs and plasma-derived EVs immunocaptured on anti-NCAM beads or IgG beads from three healthy donors were lysed, the miRNA extracted and cDNA amplified. A) RT-qPCR analysis of hsa-miR-16-5p levels in total plasma-derived EVs and plasma-derived EVs immunocaptured on anti-NCAM or IgG beads. B) RT-qPCR analysis using the serum/plasma miRNA panel for the analysis of the expression of 86 miRNAs in total plasma-derived EVs and plasma-derived EVs immunocaptured on anti-NCAM beads from three healthy donors.

The miRNA content of both plasma-derived EVs immunocaptured on anti-NCAM beads and total plasma-derived EVs from three healthy donors was then examined using the human serum/plasma miRCURY LNA focus qPCR panel, which allows quantification of 86 miRNAs commonly found in serum/plasma. The majority of the 86 miRNAs were detected in total plasma EVs from all three donors with Cp values ranging 27-43 cycles (Figure 8B). The most highly expressed miRNAs in total plasma EVs were hsa-miR-16-5p, hsa-miR-451a and hsa-miR-126-3p. However, only between two and seven miRNAs were detected in EVs immunocaptured on anti-NCAM beads from the plasma of three healthy donors (Figure 8B). Amplification of spike-in controls were similar in all samples indicating similar efficiencies of RNA isolation and RT-qPCR reactions for all samples. Finally, the miRNA Tissue Atlas 2 database [35] was used to generate a heat map of the tissue distributions of the 10 miRNAs detected in EVs immunocaptured on the anti-NCAM beads. This revealed that the majority of these miRNAs are enriched in tissues or organs such as blood vessels, lung, bone, thyroid and heart whereas none of the miRNAs immunocaptured on anti-NCAM beads were shown to be enriched in the brain (Supplemental Data File 1 Figure 9).

## Discussion

The enrichment of nEVs from biofluids such as plasma offers a non-invasive technique to indirectly observe processes in the brain of patients with neurodegenerative and psychiatric conditions. Finding a neuron-specific marker that is expressed on the surface of nEVs is therefore crucial to allow the specific immunocapture and enrichment of nEVs from biofluids for their use as biomarkers for brain disorders to aid in the diagnosis and therapy of patients. Two neuronal adhesion molecules have been used previously for the enrichment of nEVs from biofluids: L1CAM and NCAM. Therefore, this study examined in detail the enrichment of EVs from both cell culture media and plasma using anti-NCAM immunocapture beads.

EVs were isolated from either cell culture media or human plasma and were characterised according to current MISEV guidelines [36]. EVs from SH-SY5Y cells showed the expected size distributions, a cup-shaped morphology by TEM, and the presence of the EV markers CD81 and Tsg101. Flow cytometry and immunoblot analysis showed that NCAM-positive SH-EVs immunocaptured on anti-NCAM beads also expressed the EV markers CD81 (Figure 3) and Tsg101 (Figure 5A) which were absent in samples incubated with IgG beads, anti-calnexin beads, or beads alone. The presence of the cytoplasmic protein Tsg101 and the lack of calnexin in SH-EVs immunocaptured on anti-NCAM beads (Figure 5A), together with the loss of the anti-CD81-PE fluorescence signal after treatment of anti-NCAM immunocaptured SH-EVs with detergent [31] (Figure 3), indicated that intact vesicles rather than cell debris or protein aggregates were immunocaptured by anti-NCAM beads. This was further confirmed by SEM analysis which permitted direct visualisation of the presence of vesicle structures on the surface of anti-NCAM beads (Figure 5B). These data therefore confirmed that anti-NCAM immunocapture beads can be used to specifically isolate NCAM-positive EVs from cell conditioned media.

According to the Protein Atlas (https://www.proteinatlas.org/), NCAM is expressed across all brain regions, but the expression of this receptor is not strictly neuron-specific as NCAM is also expressed by natural killer (NK) cells, some T cell populations, oligodendrocytes and astrocytes. Nonetheless, a recent study by Huang *et al*. (2023) [24] detected abundant expression of NCAM on EVs isolated from eight different brain regions, with the highest NCAM levels on EVs derived from the medulla. Ali Moussa *et al*. (2022) [25] found that EVs released from induced pluripotent stem cell (iPSC)-derived cortical neurons and brain organoids express NCAM. Furthermore, single EV flow cytometry detected NCAM on a population of EVs isolated from plasma, with the loss of fluorescent signal following treatment with detergent indicating that NCAM was on intact EVs [25]. A proteomics-based study also identified NCAM alongside ATP1A3 as specific markers of EVs derived from excitatory neurons established from iPSCs [22]. Finally, a recent study further characterised NCAM expression on nEVs from multiple sources including iPSC-derived neurons, brain tissue, CSF and plasma using highly sensitive methods such as a Single Particle Interferometric Reflectance Imaging Sensor (SP-IRIS) assay, TEM and super resolution microscopy [23]. This study detected NCAM, and to a lesser extent L1CAM, on nEVs from various sources associated with the EV markers CD9, CD81 and CD63 [23]. NCAM has been used as a neuronal marker for the immunocapture of nEVs from plasma in several previous studies which compared levels of Alzheimer disease-related proteins such as tau and amyloid beta 42 (Aβ42) in nEVs from patients and healthy controls to determine their potential use as biomarkers of disease [14,15,26,27]. Importantly, some of these studies also detected the EV markers CD81 or Alix [15,26] in the anti-NCAM immunocaptured plasma-derived EVs, indicating the immunocapture of nEVs. You *et al.* (2023) [23] also immunocaptured brain-derived EVs using anti-NCAM beads which were shown to have several EV-specific and neuronal-specific markers. Therefore, our findings are in agreement with these studies that full length NCAM is released from neuronal cells in EVs and can be used to immunocapture these EVs using an anti-NCAM bead approach. However, this is the first study to use a range of techniques including SEM to show that NCAM-positive EVs derived from cell conditioned media are immunocaptured on the surface of anti-NCAM immunocapture beads.

In this study, the L1CAM antibodies 5G3 and UJ127 which target the N-terminal region of the extracellular domain of L1CAM, and the fourth fibronectin domain respectively, did not immunocapture SH-EVs as shown by flow cytometry and immunoblot analysis, although this may be due to lower expression levels of full-length L1CAM compared to NCAM in SH-EVs (Figure 5A and Supplemental Data File 1 Figure 2). Many studies have used anti-L1CAM targeted immunocapture methods to enrich for nEVs from plasma or serum from patients with various neurodegenerative and psychiatric conditions [11,37,38,39,40,41], and there is evidence that EVs immunocaptured using anti-L1CAM antibodies have enrichment of brain-specific miRNAs [42,43] and neuronal-specific proteins such as glutamate receptor 1, neuron-specific enolase, tau and Aβ42 [14,15,44]. However, L1CAM is not neuron-specific because it is expressed in other cell types and tissues [16,17,18,19,20]. Furthermore, the extracellular domain of L1CAM is susceptible to proteolysis resulting in the release of the extracellular domain from the cell surface and EVs [45,46]. This is highlighted by the fact that in this study the main form of L1CAM detected in SH-EVs was a 130 kDa fragment (Supplemental Data File 1 Figure 2) which can be generated by proteolysis of the extracellular domain by plasmin [47,48]. Finally, a study by Norman *et al.* (2021) [21] found that most L1CAM in plasma is in the form of the soluble extracellular domain and is not associated with EVs.

Following the successful immunocapture of SH-EVs using anti-NCAM immunocapture beads, we focused on whether this neuronal marker could be used to enrich nEVs from human plasma. Plasma-derived EVs showed the expected size distributions, cup-shaped morphology, and the presence of EV markers CD81, CD63 and Tsg101 (Figure 6). Lipoprotein ApoAI was also present in the eluted EVs indicating carry-over of lipoproteins during size exclusion chromatography, possibly due to their similar size to EVs [49,50]. Surprisingly, immunoblot analysis of plasma-derived EVs showed that both Tsg101 and alpha-tubulin had higher molecular weights than expected (Figure 6), which may be due to post-translational modifications of these proteins within the plasma-derived EVs such as sumoylation [51,52,53]. Since immunoblot analysis was not sensitive enough to detect NCAM even in the total plasma-derived EVs (Supplemental Data File 1 Figure 6), a highly sensitive electrochemiluminscent ELISA-based assay was used to detect NCAM antigen in plasma-EVs. This revealed the presence of NCAM antigen in plasma-derived EVs immunocaptured on anti-NCAM beads but not IgG control beads (Figure 7A) indicating the enrichment of NCAM-positive EVs from plasma. It is possible soluble NCAM was present in plasma [54]. However, size exclusion chromatography isolation of plasma-derived EVs should remove soluble proteins. We did not detect CD9 on NCAM-positive EVs using an anti-CD9 ELISA. This is in contrast to the study by You *et al.* (2023) [23] which observed the colocalisation of CD9 and NCAM on neuronal EVs using TEM and single-EV techniques. This highlights the importance of using these more sensitive techniques for the validation of the presence of nEVs on immunocapture beads. SEM analysis of anti-NCAM beads incubated with plasma-derived EVs also confirmed the presence of vesicle-like structures on the surface of the beads (Figure 7B). However, these vesicles were also present on IgG and anti-calnexin beads following incubation with plasma-EVs, indicating some non-specific binding of plasma-derived EVs to control beads. A few images of beads not incubated with plasma-derived EVs also had spherical structures which may be the result of SEM processing forming “salt bubbles” [55], or the polystyrene surface of the beads. ImageJ analysis of the diameters of the EVs revealed that all beads, even those not incubated with EVs, had spherical structures ranging from 35-55 nm, possibly due to the polystyrene material of the bead surface, whereas anti-NCAM beads also had vesicle-like structures of 56-95 nm which are likely to be EVs (Figure 7C). Immuno-gold labelling of the EVs on immunocapture beads using antibodies against EV markers or neuronal-specific markers would be required to definitely confirm that the vesicle structures on the anti-NCAM immunocapture beads are EVs. However, these data support previous findings that NCAM can be used a marker to enrich NCAM-positive EVs from plasma [14,15,26].

Since NCAM can also be expressed by NK cells, an additional step was included before the enrichment of nEVs using anti-NCAM immunocapture beads to deplete NK cell-derived EVs. NCAM was not detected in EV lysed from anti-NKG2A beads (Figure 7A), suggesting that NK cell-derived NCAM-positive EVs were at levels below the detection limit of the MSD NCAM assay. However, since inflammatory cytokines increase the release of EVs from NK cells [56], it is possible that levels of NK cell-derived EVs in the peripheral circulation may be higher in patients with neurodegenerative or psychiatric conditions. It is also recognised that the addition of the NK-EV depletion step increases complexity of the EV isolation procedure and time to isolate nEVs when processing a large number of samples.

Finally, the miRNA content of plasma-derived EVs immunocaptured on anti-NCAM beads was examined using RT-qPCR. Over 50 of the 86 miRNAs in the serum/plasma small focus qPCR panel were detected in total plasma EVs from the three healthy donors, whereas only a small number of miRNAs (two to seven) were detected from anti-NCAM beads incubated with plasma-EVs (Figure 8B). The majority of miRNAs detected on anti-NCAM bead samples were the most highly expressed miRNAs in the total EV fraction. For example, hsa-miR-16-5p which is expressed at high levels in plasma [34], was detected in total plasma EV samples and at lower levels in plasma-EVs samples immunocaptured using anti-NCAM beads or IgG beads from all three donors (Figure 8A), possibly due to non-specific binding of miRNAs or EVs to the beads. Comparison of the 10 miRNAs detected from the anti-NCAM beads to the miRNA Tissue Atlas 2 database [35] indicated there was no enrichment of brain-derived miRNAs in the EVs immunocaptured on anti-NCAM beads (Supplemental Data File 1 Figure 9). This is in contrast to the study by Cha *et al.* (2019) [42] which detected increased levels of the brain-specific hsa-miR-9 in L1CAM-immunocaptured plasma-derived EVs compared to total EVs, suggesting enrichment of nEVs. This indicates that the immunocapture of NCAM-positive EVs from small volumes of plasma may not allow capture of sufficient amounts of material for reliable qPCR analysis of miRNA expression and future studies should be aware of this limitation.

A potential limitation of this study was the analysis of miRNAs using a qPCR panel with a defined number of miRNAs available as opposed to a more unbiased approach such as RNASeq, although many brain-specific miRNAs such as hsa-miR-9-5p [57] and hsa-miR-124-3p [58] were present in the panel but were not detected in anti-NCAM immunocaptured plasma-derived EVs. Specifically examining the expression of miRNAs known to be enriched in the brain or CNS would be a more efficient way of confirming the neuronal origin of immunocaptured nEVs. The small volume of plasma (0.5 ml) used may have limited nEV enrichment, although small volumes are often used for clinical studies which seemed appropriate for the relatively high-throughput method development we envisioned. It must also be considered that miRNAs can be associated with lipoproteins [59]. Since SEC of plasma samples also resulted in the elution of lipoproteins as observed by the presence of ApoAI (Figure 6), plasma-EV miRNAs may also include miRNAs bound to lipoproteins. Another limitation of this study is that the percentage of nEVs in blood is likely to be low compared to the large number of circulating EVs derived from leukocytes, platelets and erythrocytes [60]. In fact, a study by Li *et al.* 2020 [61] estimated that only 0.2 % of plasma EVs are derived from solid tissues, and of this population only 0.64 % are from the brain. A more recent study showed that only 0.9% of pan-tetraspanin-positive EVs isolated from human plasma were NCAM-positive [23].

In conclusion, this study has characterised the enrichment of nEVs using an anti-NCAM immunocapture method from both cell culture media and human plasma. Our data indicate that NCAM is a suitable marker for the enrichment of NCAM-positive EVs from cell culture media but has limitations when immunocapturing nEVs from small volumes of plasma for miRNA analysis, possibly due to low levels of nEVs in the peripheral circulation, low expression of NCAM on nEVs, and other potential sources of NCAM-positive EVs. This indicates that using a combination of immunocapture beads targeting multiple neuronal markers could be used to more efficiently isolate neuronal EVs. Future studies using proteomics to examine the protein content of EVs isolated from biological fluids using this immunocapture technique would also be useful to validate whether this approach is suitable for small volumes of plasma typically used in clinical studies. In addition, the use of sensitive single-EV techniques such as super resolution microscopy and SP-IRIS allows the validation and characterisation of immuno-captured EVs in detail [23,62]. Therefore, the challenge remains to identify specific and sensitive neuronal markers that are expressed at high levels on nEVs, potentially such as ATP1A3 [22,23], for the definitive isolation of nEVs from biofluids, ideally from small volumes of starting material to permit efficient isolation of nEVs in clinical studies for subsequent cargo analysis.

## Acknowledgements

We would like to thank Hash Patel for organising blood collection and to all blood donors and phlebotomists in the Department of Cardiovascular Sciences, University of Leicester. We would also like to thank Josh Whittingham at the Electron Microscopy Facility at the University of Leicester for preparing bead-EV samples for scanning electron microscopy and image acquisition. Finally, thanks to the Flow Cytometry Facility at the University of Leicester for support with flow cytometry and the use of the BD FACSCanto II flow cytometer.

## Declarations

### Ethics approval and consent to participate

Blood collection was approved by the University of Leicester Ethical Practices Committee (application 23825-fg36-Is) and all subjects provided written informed consent.

### Consent for publication

We confirm that all authors have read the manuscript and approved the submission of the manuscript.

### Availability of data and materials

The data that support the findings of this study are available upon reasonable request to the corresponding author.

## Authors’ contributions

Mary Collier: conceptualisation, methodology, investigation, writing - original draft. Natalie Allcock: methodology, investigation, resources, writing – review and editing. Nicolas Sylvius: methodology, resources, writing – review and editing. Jordan Cassidy: investigation. Flaviano Giorgini: conceptualisation, writing – review and editing, supervision, funding acquisition.

## Data Availability Statement

**Clinical Trial Number**: not applicable

