## Supplemental Data File 1 for "Examination of the enrichment of neuronal extracellular vesicles from cell conditioned media and human plasma using an anti-NCAM immunocapture bead approach"

A

Non-differentiated	Differentiated

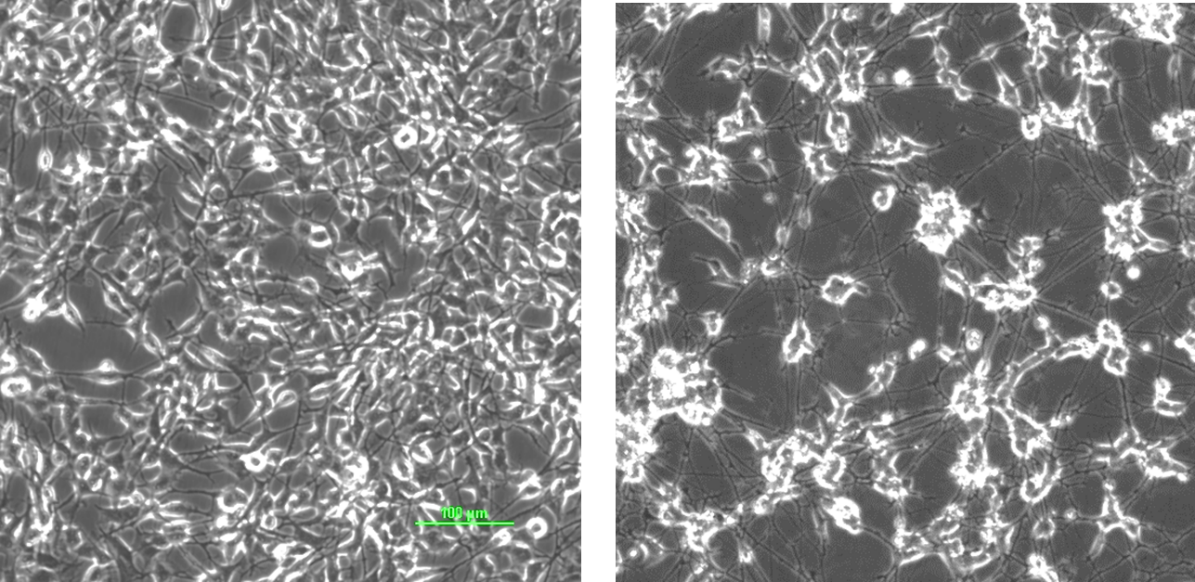

C

B

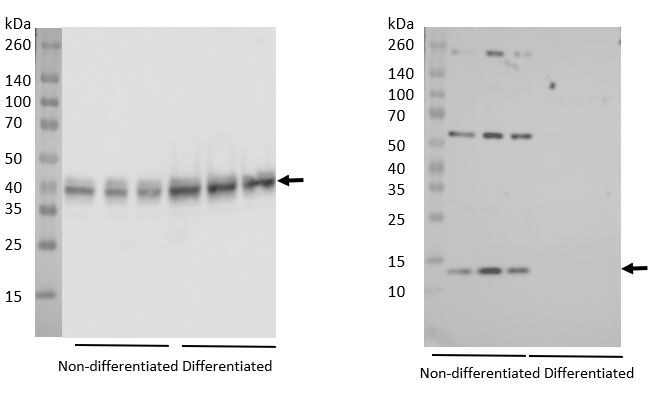

D

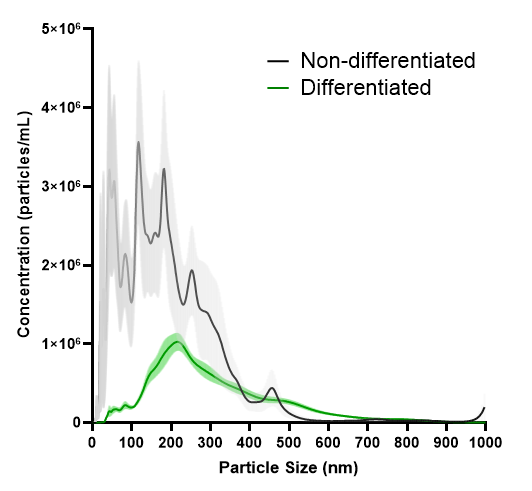

**Supplemental Figure 1.** Characterisation of differentiated and non-differentiated SH-SY5Y cells and EV release from these cells. A) Images of non-differentiated SH-SY5Y cells and cells differentiated for 11 days with 10 µM RA together with BDNF (12.5 ng/ml) for the final six days. Immunoblot analysis of synaptophysin (B) and ID2 (C) in differentiated and non-differentiated SH-SY5Y cells. D) NTA of EVs released from non-differentiated and differentiated SH-SY5Y cells.

kDa

260

140

100

70

50

40

35

25

15

SH cells	SH-EVs

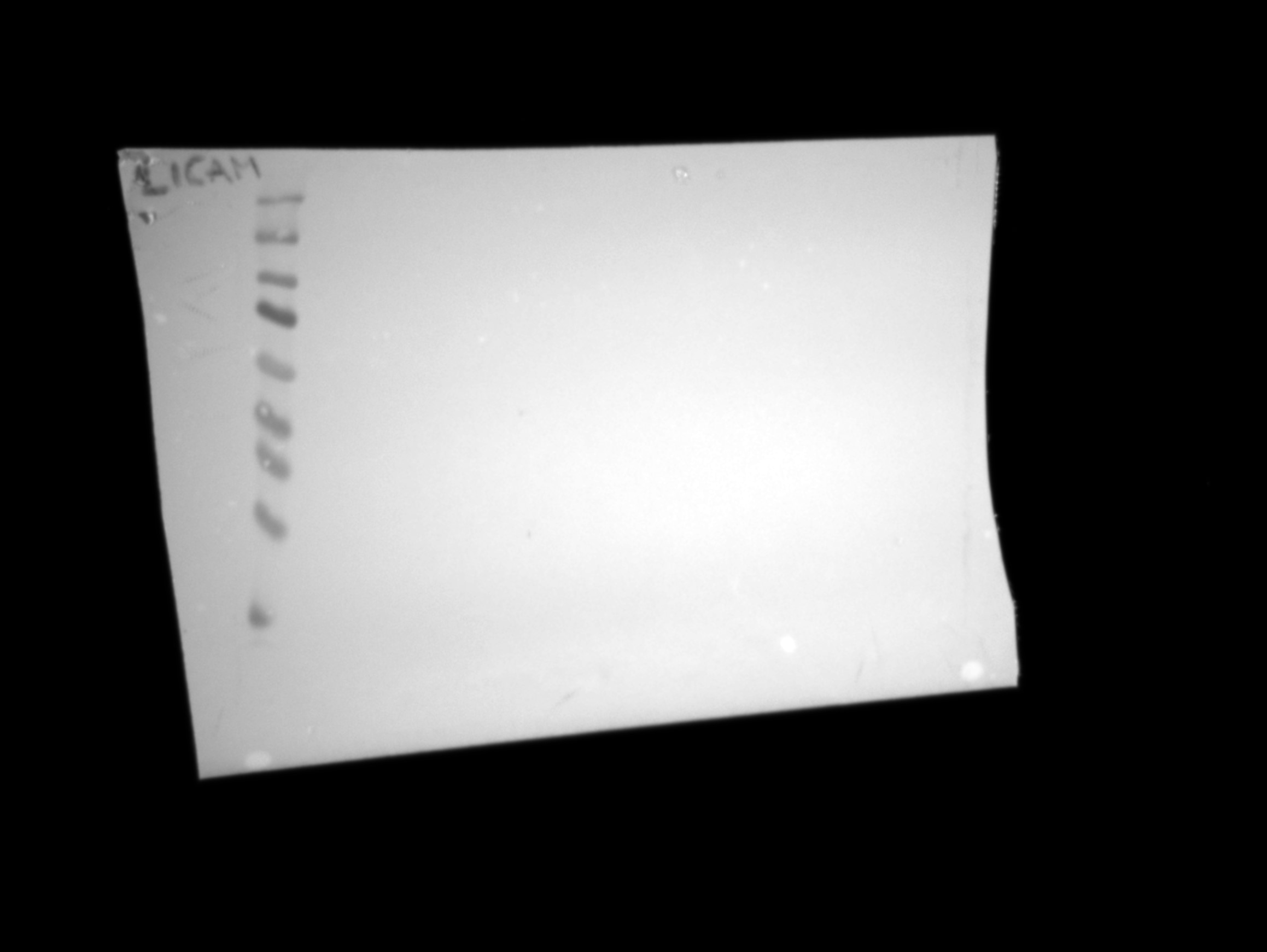

Non-specific band

130kDa fragment

Full length 200-220kDa

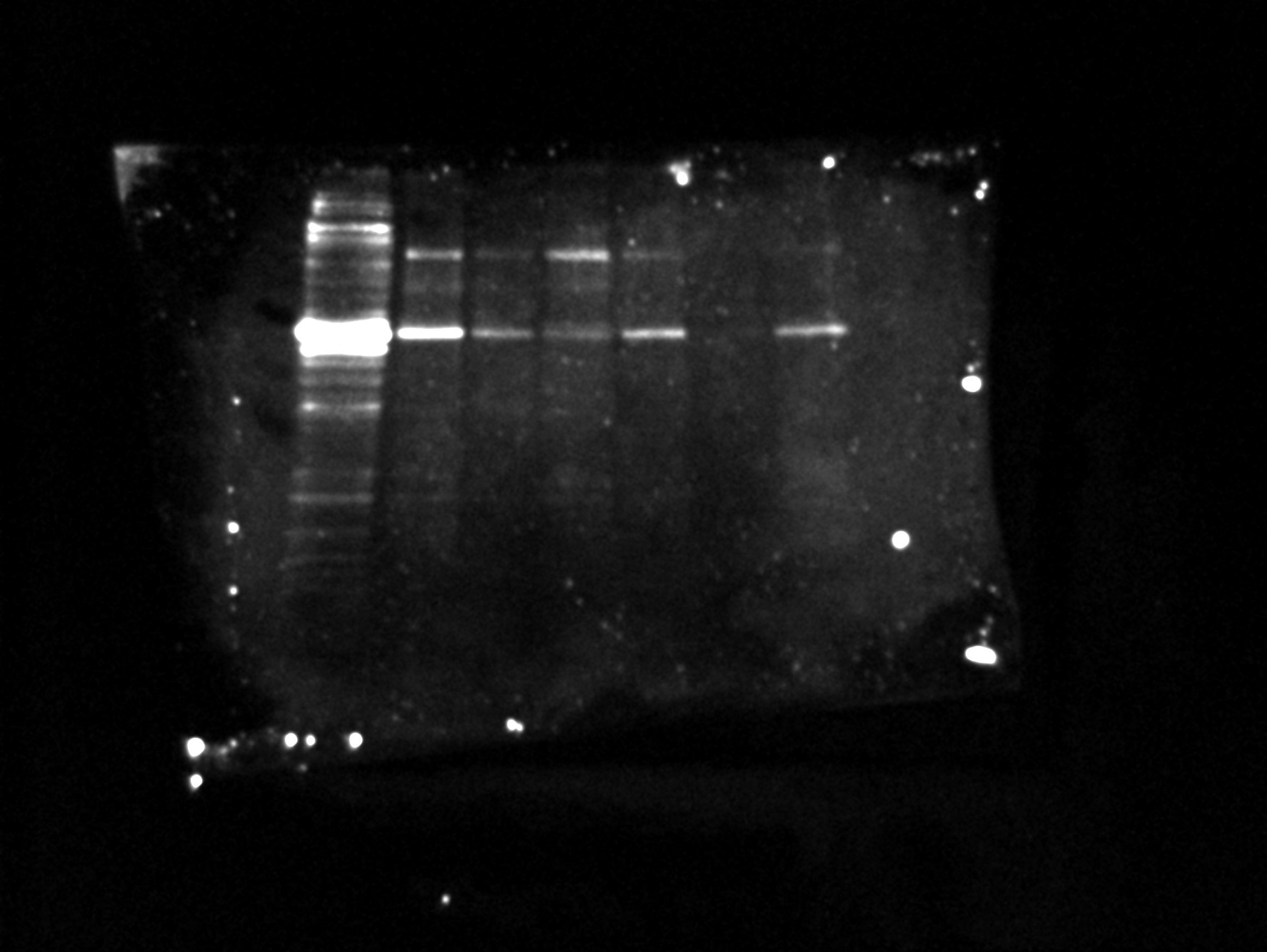

**Supplemental Figure 2.** Immunoblot analysis of L1CAM expression in SH-SY5Y EVs isolated from the conditioned media of three different T75 flasks of SH-SY5Y cells using ExoQuick.

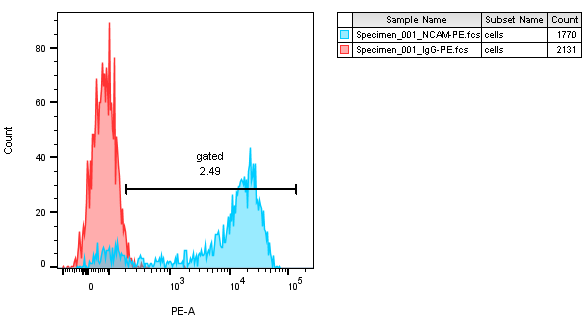

IgG-PE

Anti-NCAM-PE

**Supplemental Figure 3.** The expression of NCAM on the surface of SH-SY5Y cells was confirmed by flow cytometry. SH-SY5Y cells were detached using 2 mM EDTA in PBS and resuspended at 1.5 × 10^6^ cells in 100 µl of PBS containing 0.3 % (w/v) BSA and 0.1 % (v/v) sodium azide. Cells were labelled with anti-NCAM-PE (TULY-56) (0.06 µg in 100 µl) or the equivalent concentration of IgG-PE isotype control and incubated on ice for 1 h. Cells were washed twice with 1 ml of PBS/BSA/sodium azide, resuspended in 400 µl PBS and analysed on a Canto II flow cytometer. Red histogram = IgG-PE, blue histogram = anti-NCAM-PE.

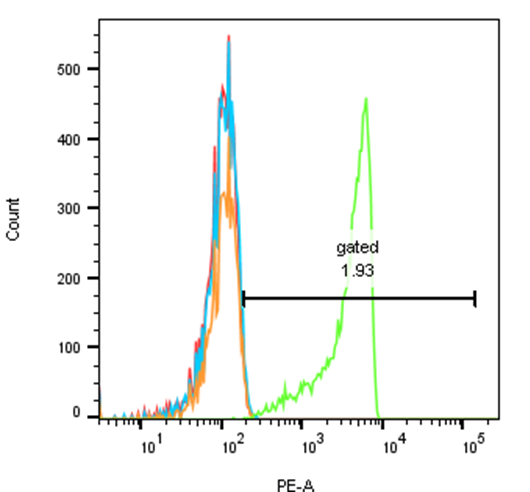

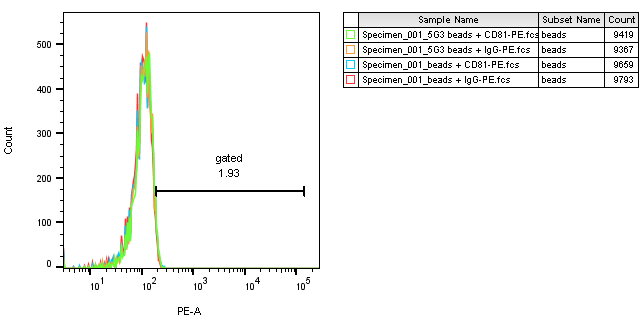

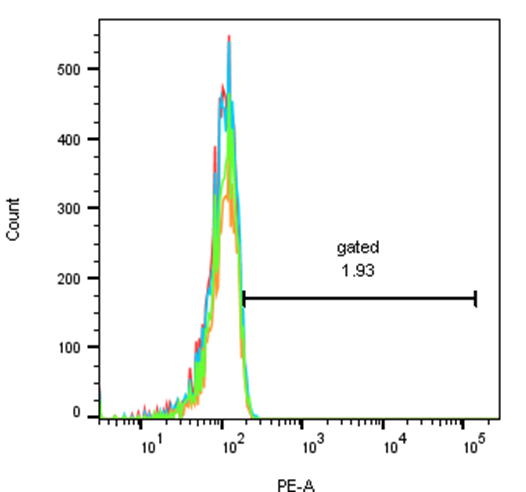

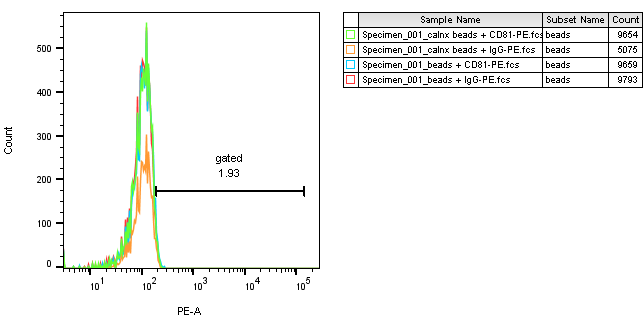

Anti-L1CAM UJ127 beads + EVs

UJ127-beads+anti-CD81-PE

UJ127-beads+IgG-PE

Control beads +anti-CD81-PE

Control beads+IgG-PE

Anti-L1CAM 5G3 beads + EVs

5G3-beads+anti-CD81-PE

5G3-beads+IgG-PE

Control beads +anti-CD81-	PE

Control beads+IgG-PE

Anti-CD81 beads

+ EVs

CD81-beads+

anti-CD81-PE

CD81-beads+

IgG-PE

Control beads

+anti-CD81-PE

Control beads

+IgG-PE

B

A

Anti-calnexin beads + EVs

Calnexin-beads+anti-CD81-PE

Calnexin-beads+IgG-PE

Control beads +anti-CD81-PE

Control beads+IgG-PE

C

D

**Supplemental Figure 4.** **Flow cytometry of CD81-positive SH-SY5Y EVs immunocaptured using anti-CD81 or anti-L1CAM Dynabeads .**SH-SY5Y cell-derived EVs (100 µl) were incubated overnight with antibody-conjugated beads and then labelled with anti-CD81-PE or IgG-PE antibodies and analysed by flow cytometry. Histograms are shown for anti-CD81 beads (A), anti-L1CAM clone 5G3 beads (B), anti-L1CAM beads clone UJ127 beads (C) and anti-calnexin beads (D) labelled with anti-CD81-PE (green histograms) or IgG-PE (orange histograms) compared to control beads (without conjugated antibody) labelled with anti-CD81-PE (blue histograms) or IgG-PE (red histograms).

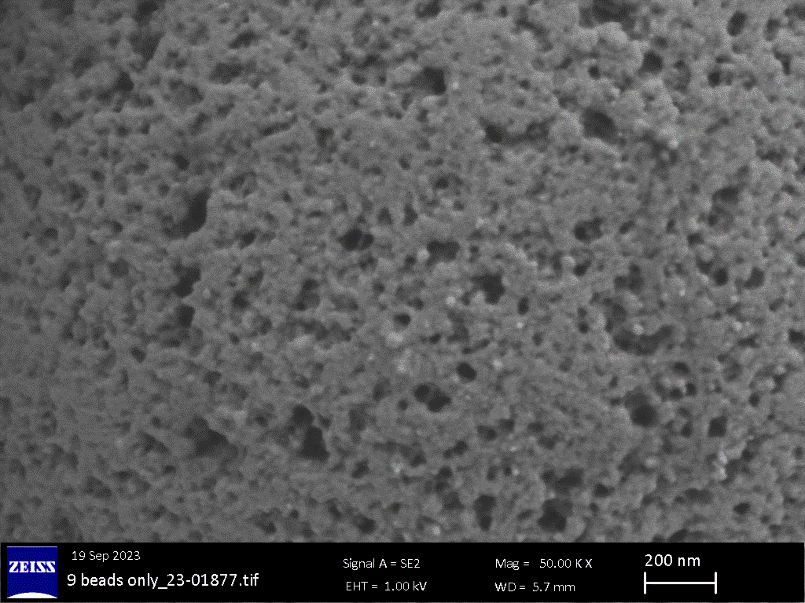

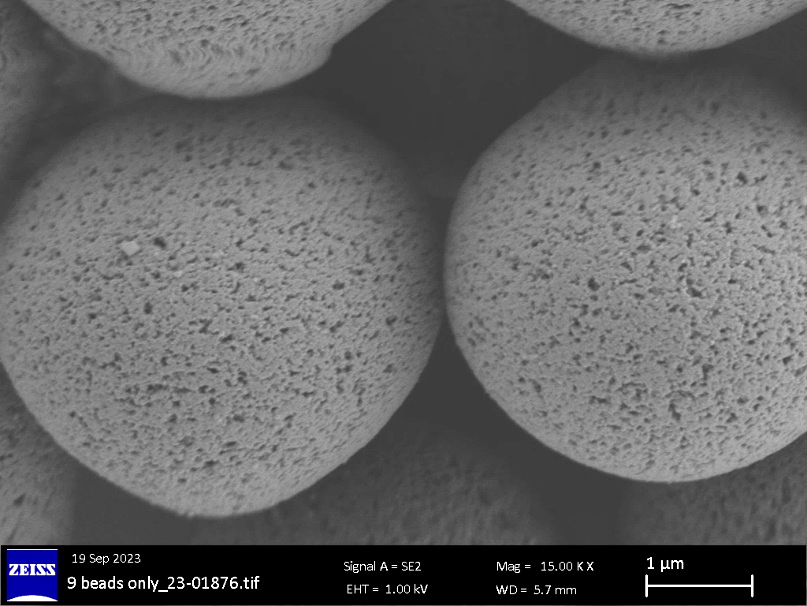

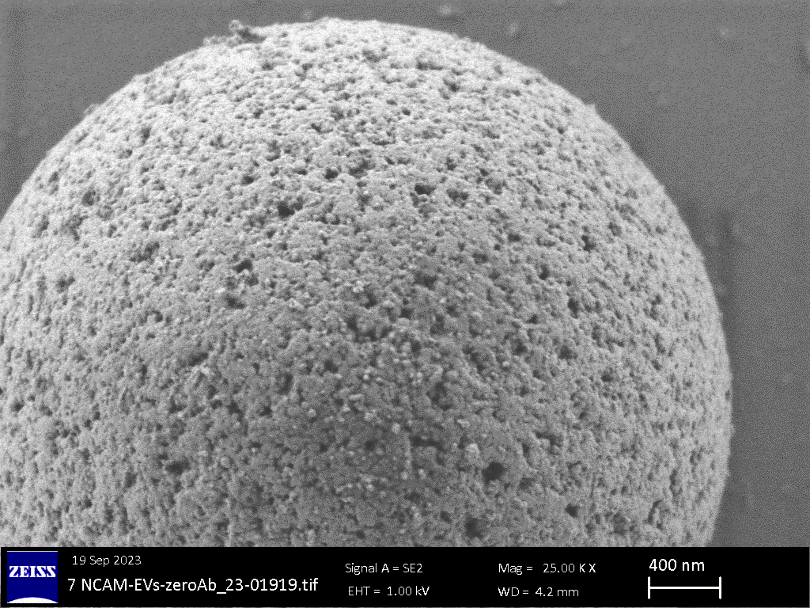

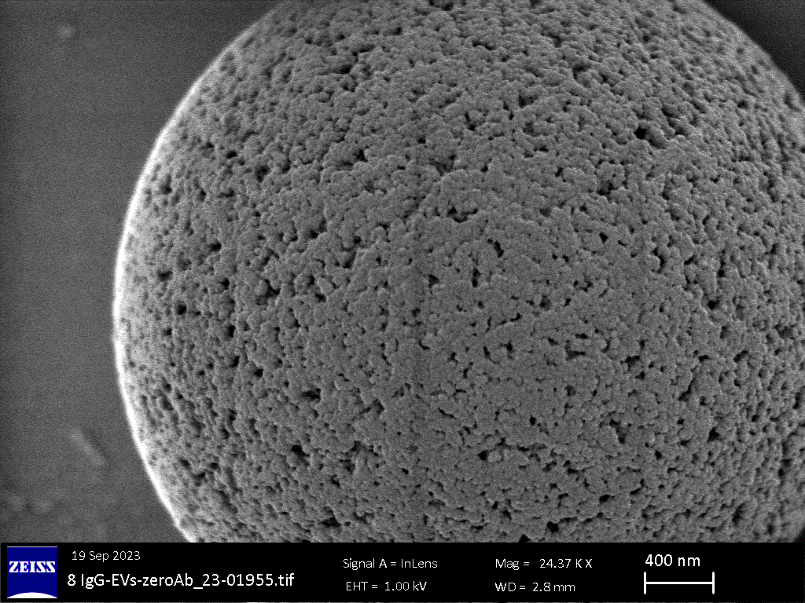

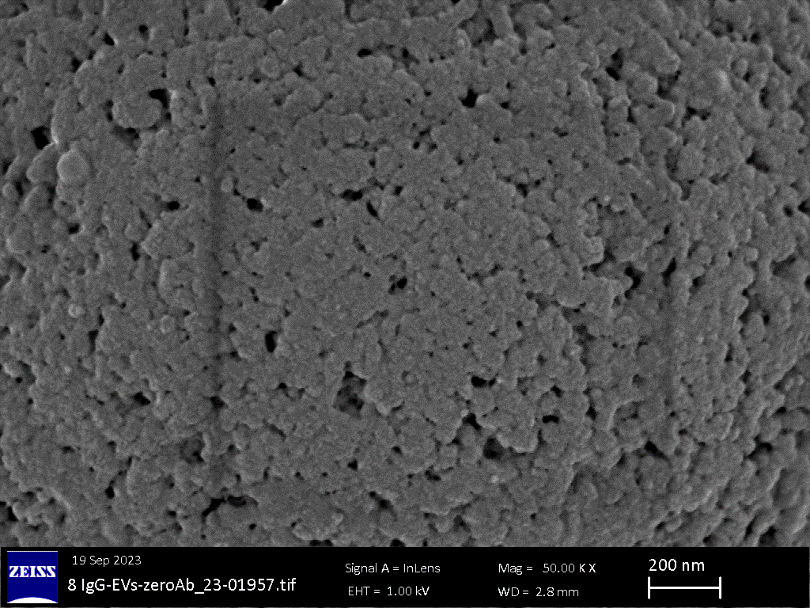

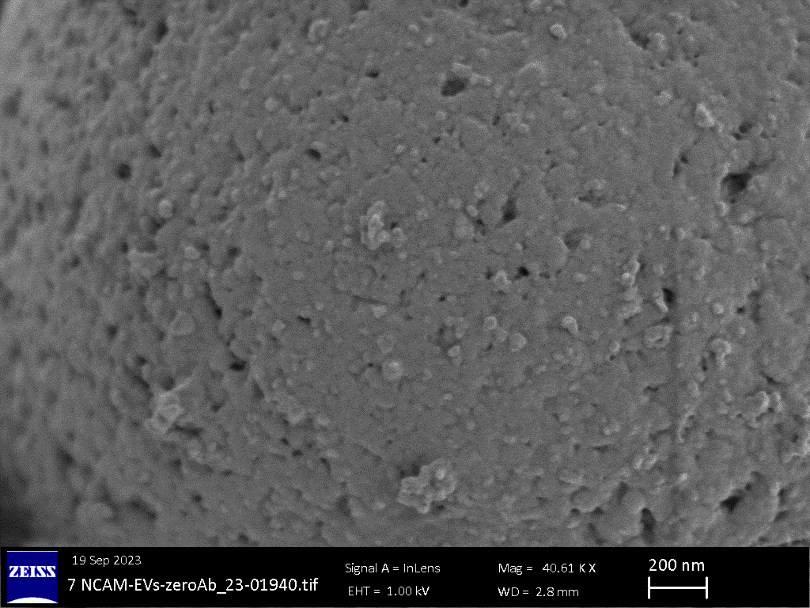

D

F

E

C

B

A

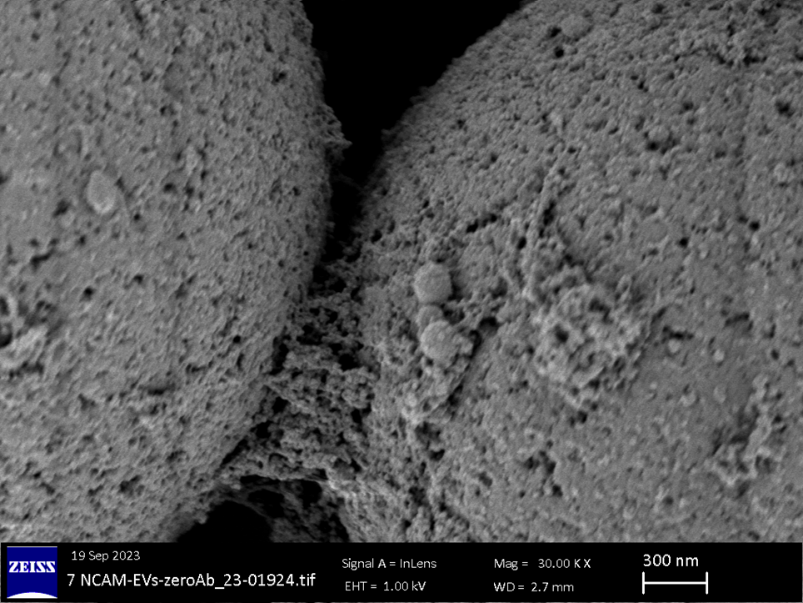

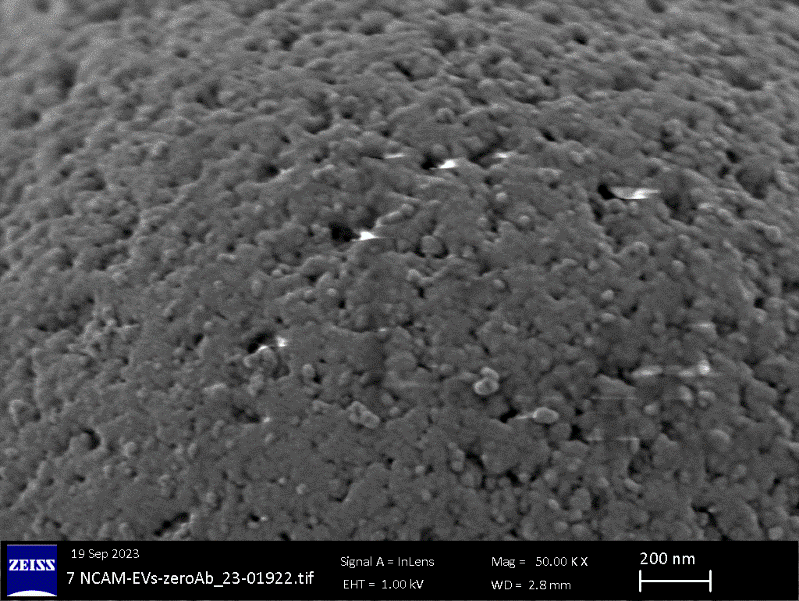

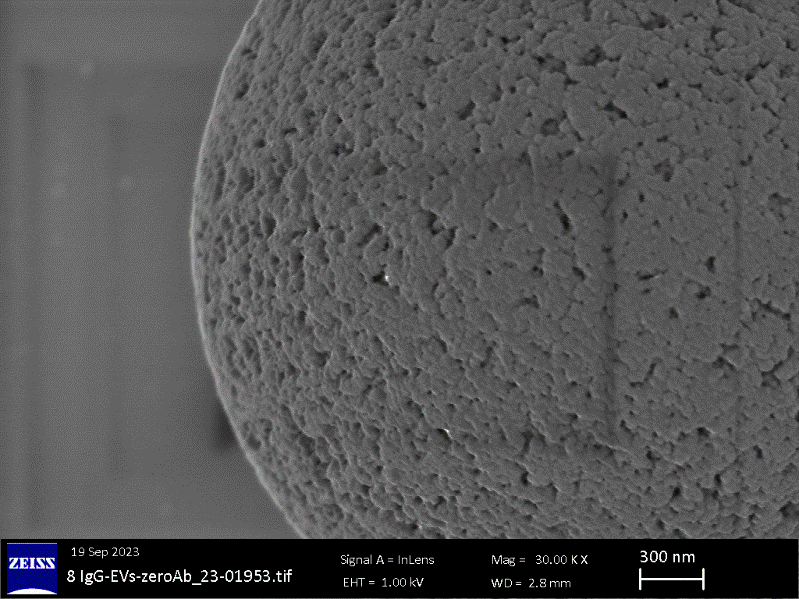

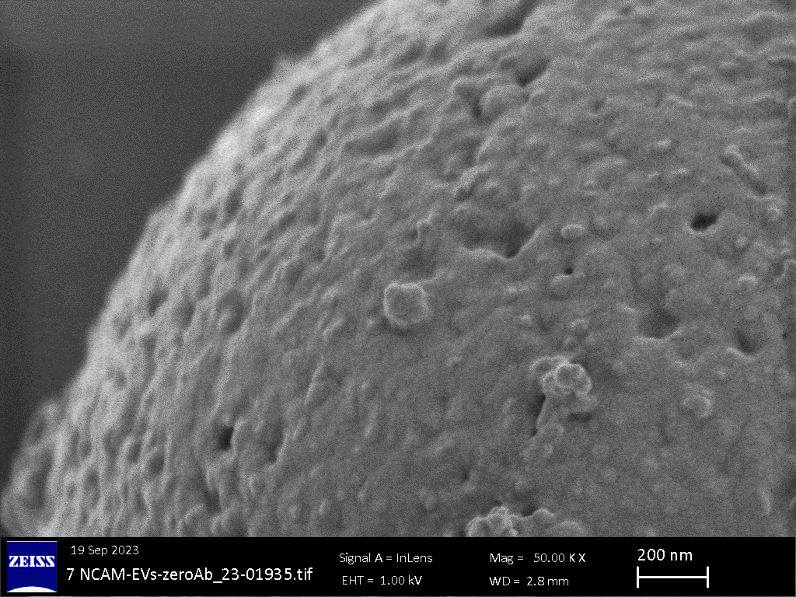

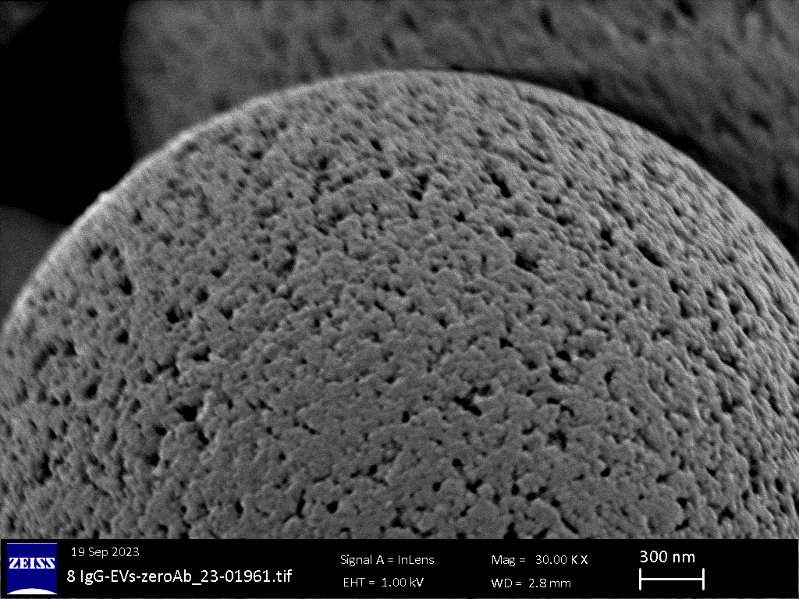

K

J

I

H

G

**Supplemental Figure 5. SEM analysis of SH-EVs on anti-NCAM beads.** A&B) SEM images of beads alone without incubation with SH-EVs. C,E,G,I) Anti-NCAM beads incubated with SH-EVs. D,F,H,J) IgG beads incubated with SH-EVs. K) Protein strands on anti-NCAM beads. Red arrows indicate EVs.

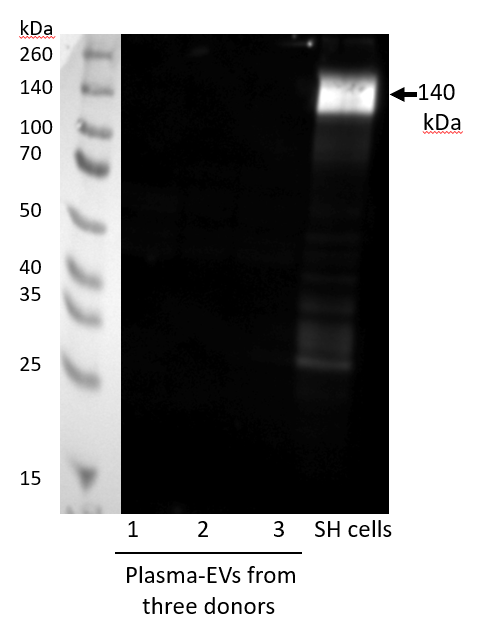

**Supplemental Figure 6.** **Immunoblot analysis of NCAM in plasma-derived EVs.** NCAM antigen was not observed in plasma-derived EVs isolated from 0.5 ml of plasma using SEC columns using immunoblot analysis. SH-SY5Y cell lysate was used as a positive control.

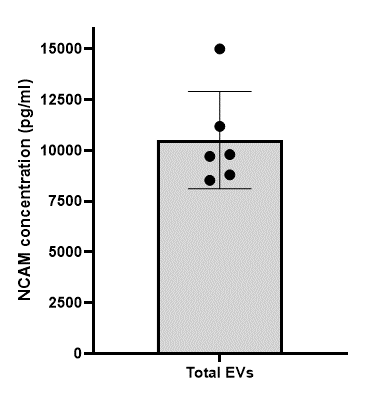

**Supplemental Figure 7.** NCAM antigen levels in total plasma EVs from six healthy donors following isolation using SEC columns and concentration using centrifugal 10kDa MWCO filters. Total EVs were lysed and NCAM concentrations were measured using the NCAM MSD assay.

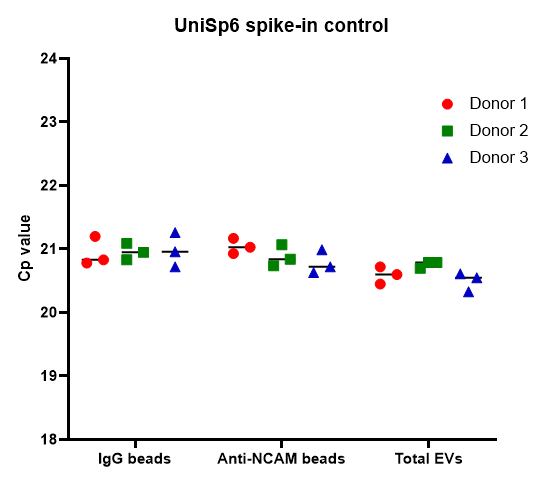

**Supplemental Figure 8.** qPCR of UniSp6 spike-in control in plasma-derived EV samples from three healthy donors. UniSp6 was spiked into the RT reaction for each sample and detected using UniSp6 primers in the miRCURY LNA miRNA qPCR assay.

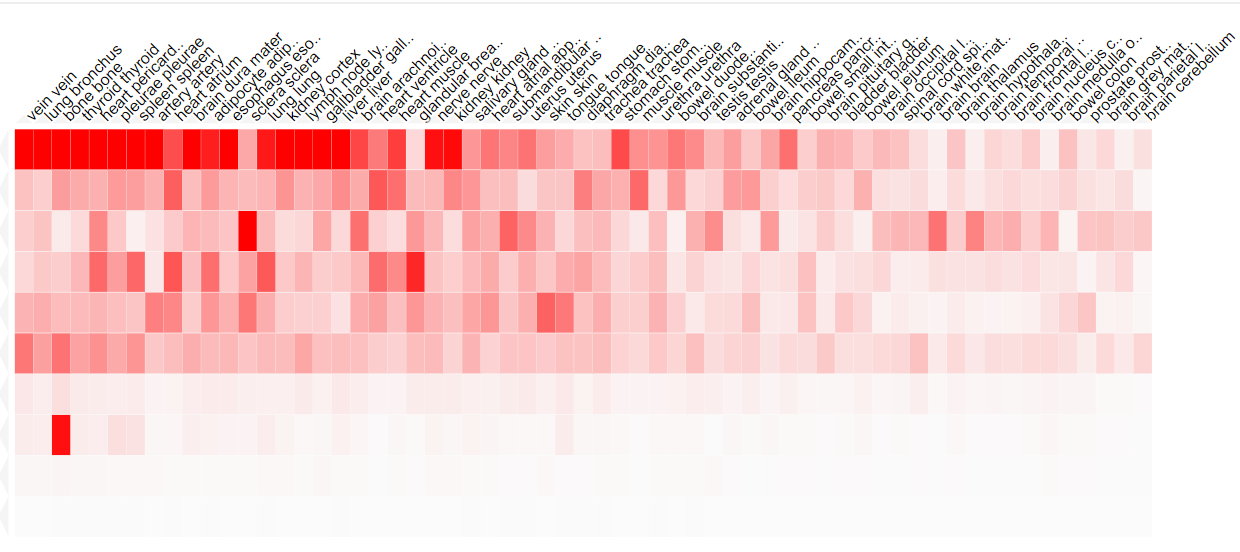

hsa-miR-451a

hsa-miR-22-3p

hsa-let-7c-5p

hsa-miR-126-3p

hsa-miR-23a-3p

hsa-miR-16-5p

hsa-miR-92a-3p

hsa-miR-223-3p

hsa-miR-25-3p

hsa-miR-17-3p

**Supplemental Figure 9. Heat map of the tissue distribution of miRNAs in EVs immunocaptured on anti-NCAM immunocapture beads.** The 10 miRNAs detected in EVs immunocaptured on anti-NCAM beads were compared to the miRNA Tissue Atlas 2 database to identify the possible tissue and organ sources of these miRNAs.
