## Supplemental Data File 2 for "Examination of the enrichment of neuronal extracellular vesicles from cell conditioned media and human plasma using an anti-NCAM immunocapture bead approach"

### Slide 1
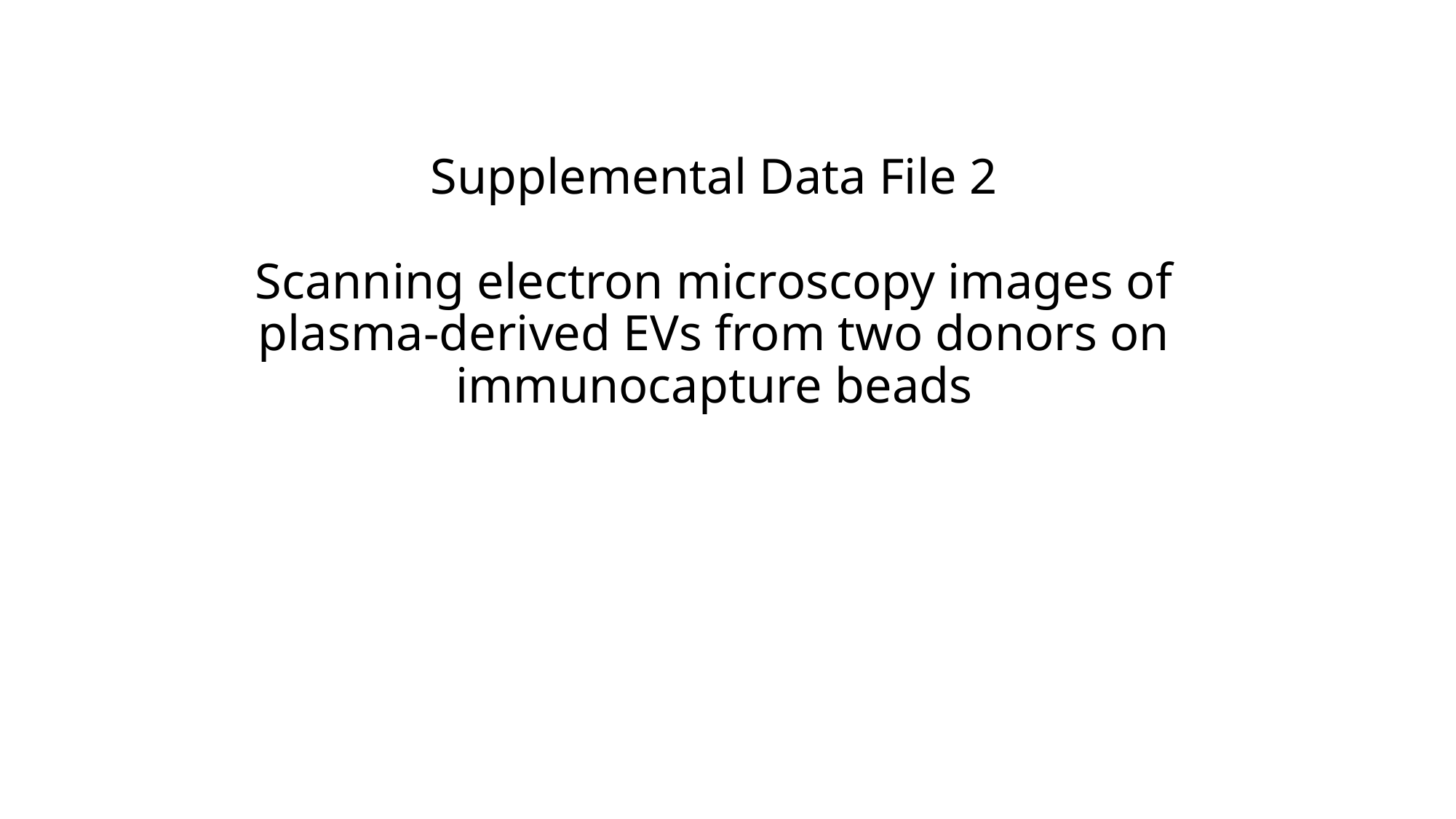

#### Supplemental Data File 2 Scanning electron microscopy images of plasma-derived EVs from two donors on immunocapture beads

### Slide 2
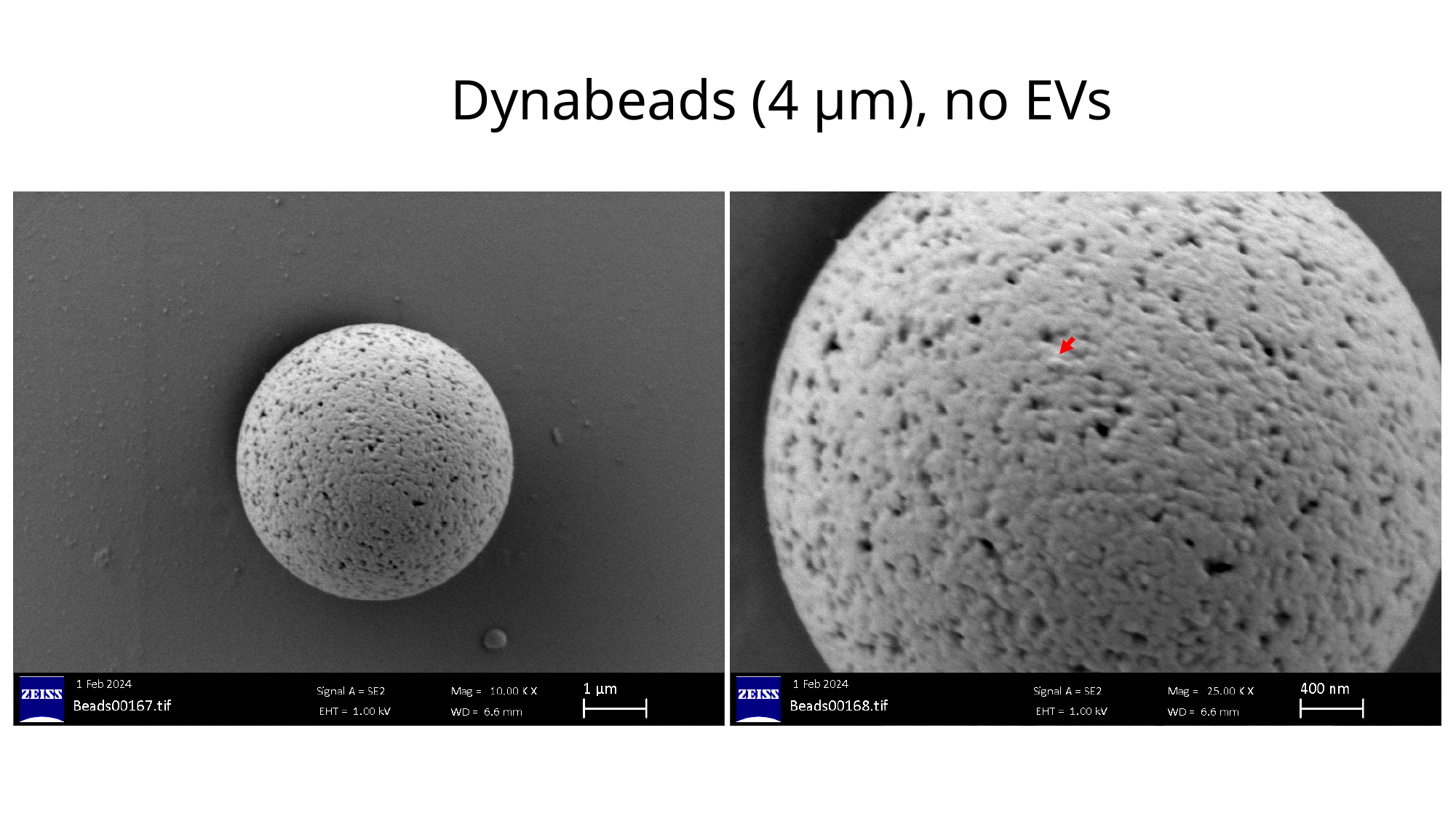

#### Dynabeads (4 µm), no EVs

### Slide 3
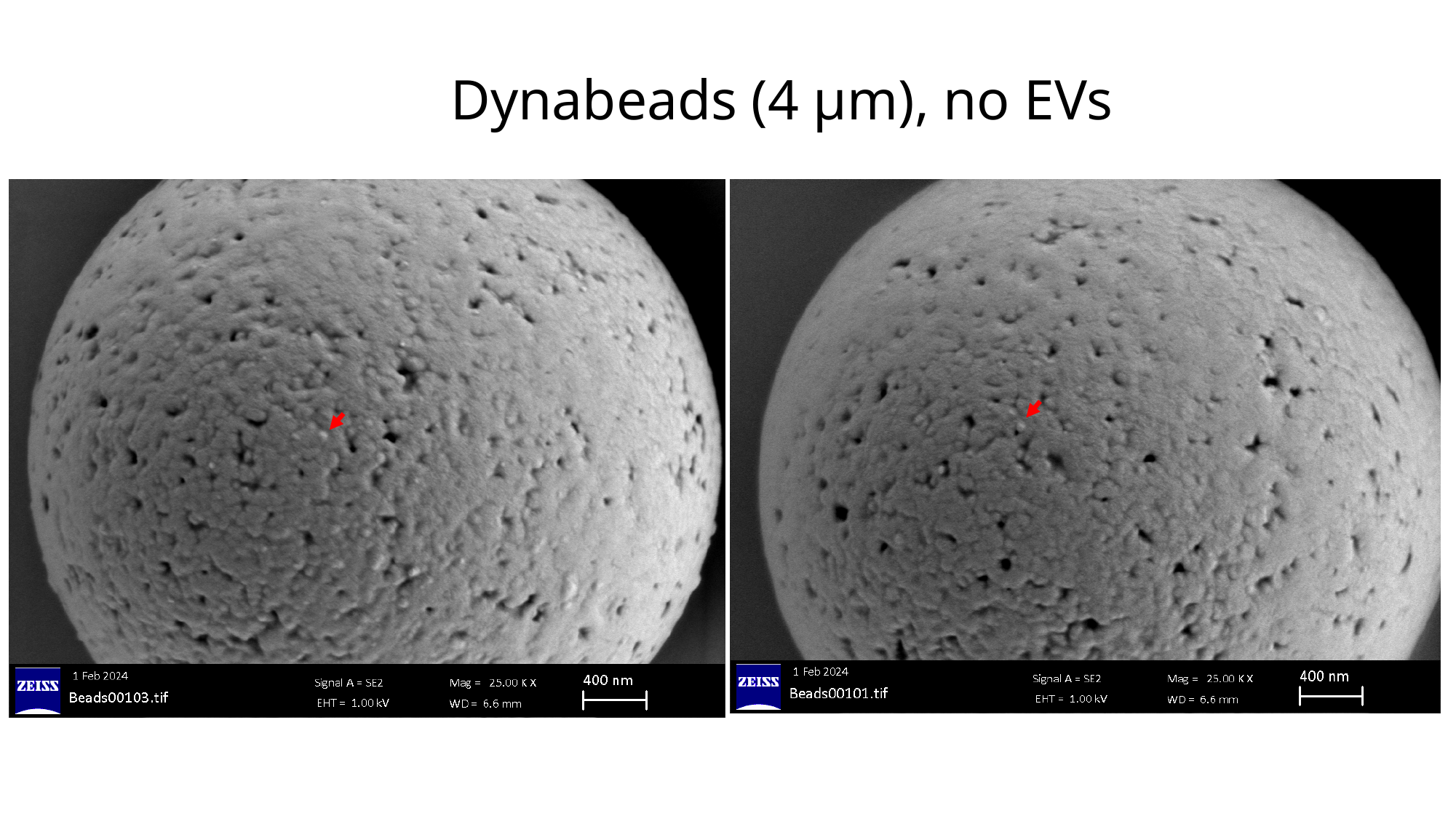

#### Dynabeads (4 µm), no EVs

### Slide 4
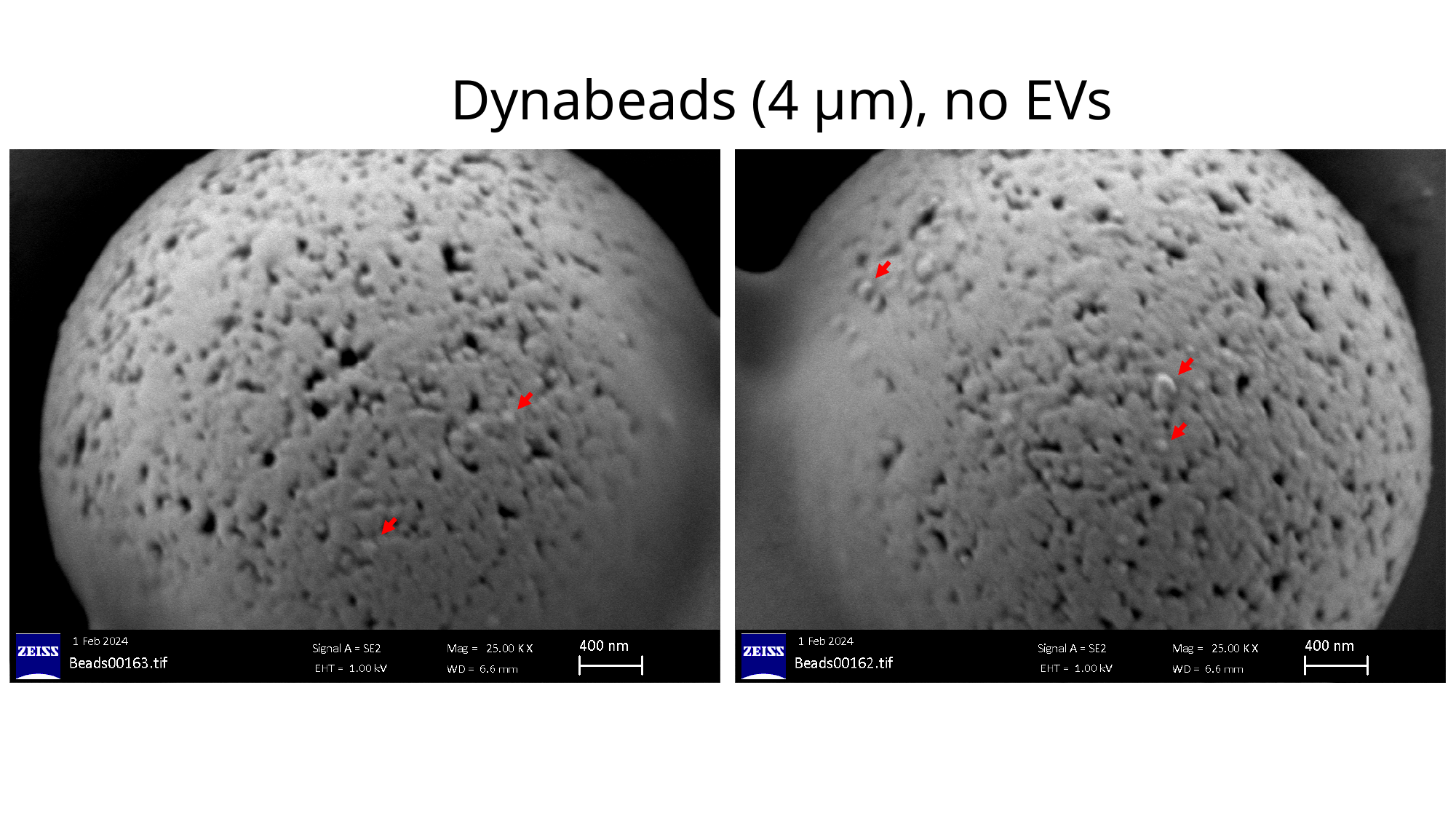

#### Dynabeads (4 µm), no EVs

### Slide 5
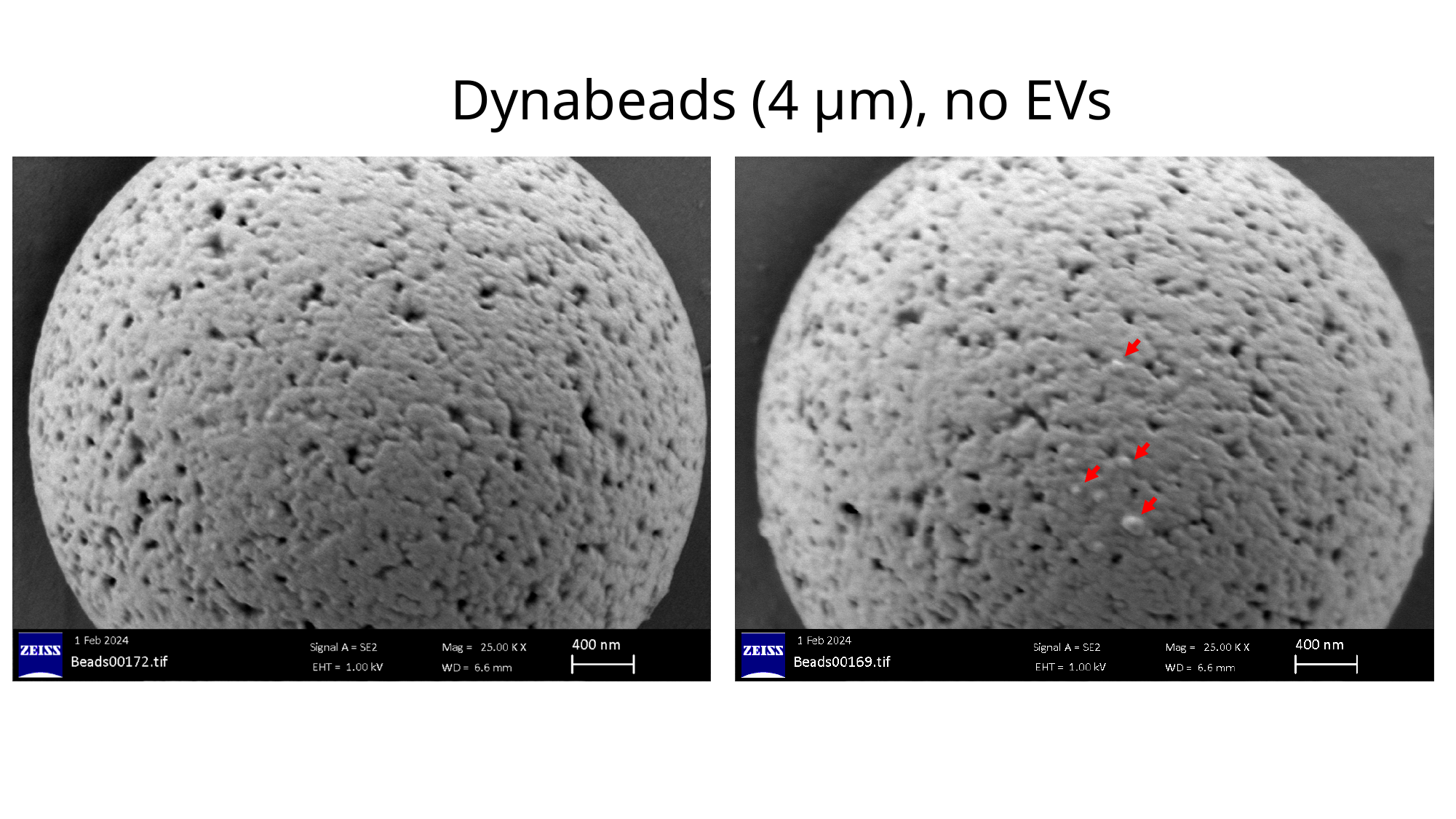

#### Dynabeads (4 µm), no EVs

### Slide 6
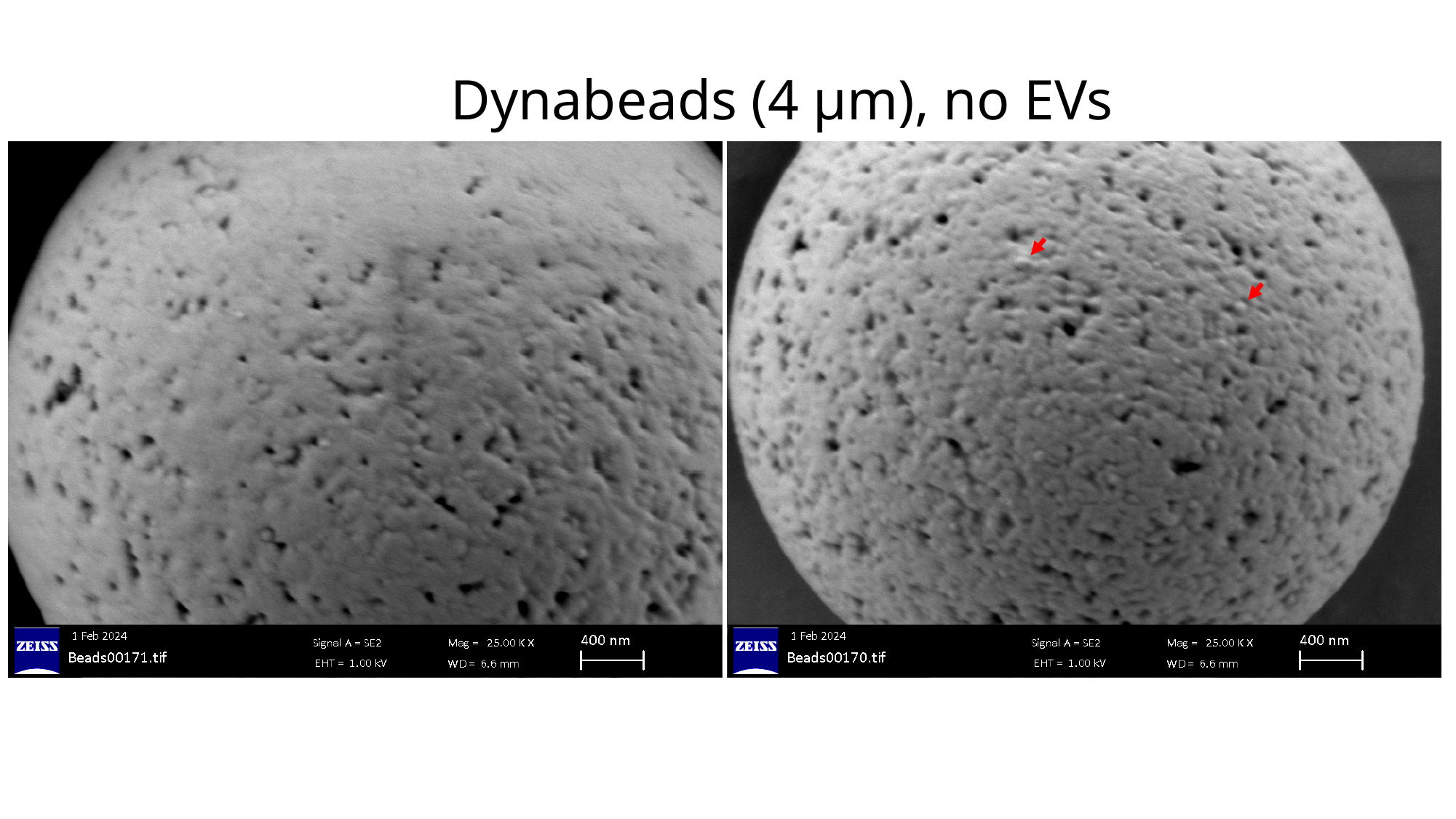

#### Dynabeads (4 µm), no EVs

### Slide 7
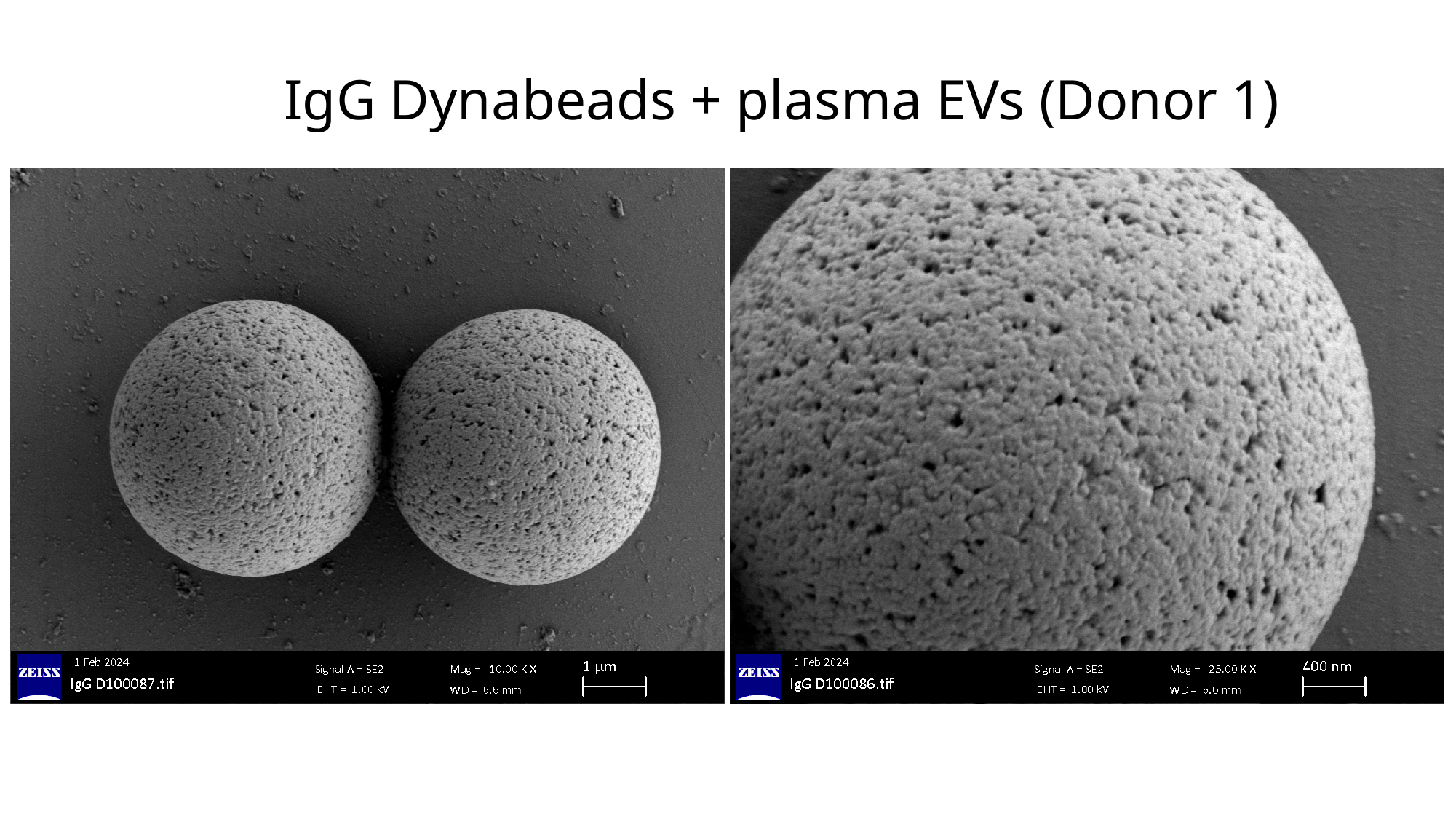

#### IgG Dynabeads + plasma EVs (Donor 1)

### Slide 8
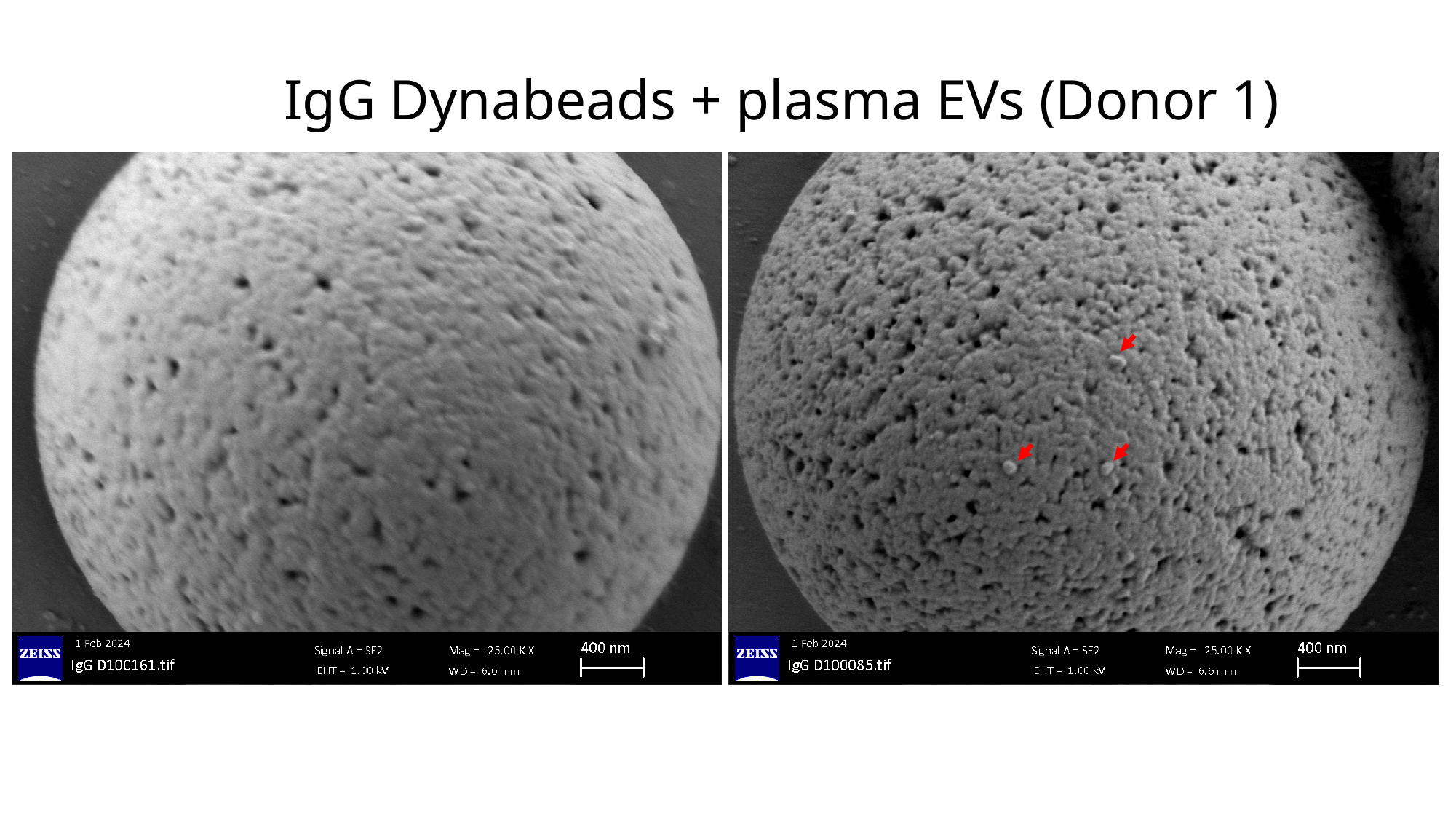

#### IgG Dynabeads + plasma EVs (Donor 1)

### Slide 9
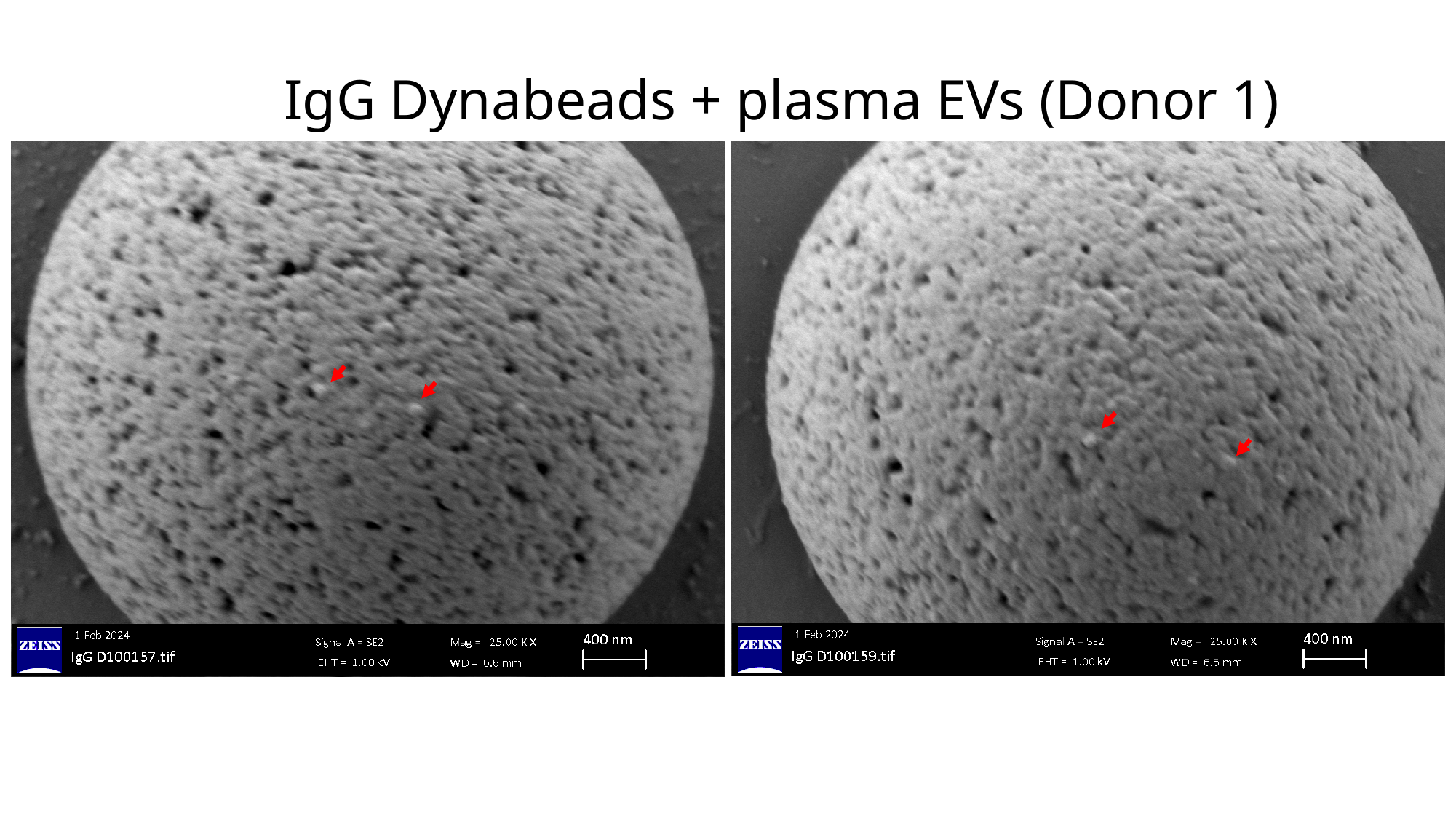

#### IgG Dynabeads + plasma EVs (Donor 1)

### Slide 10
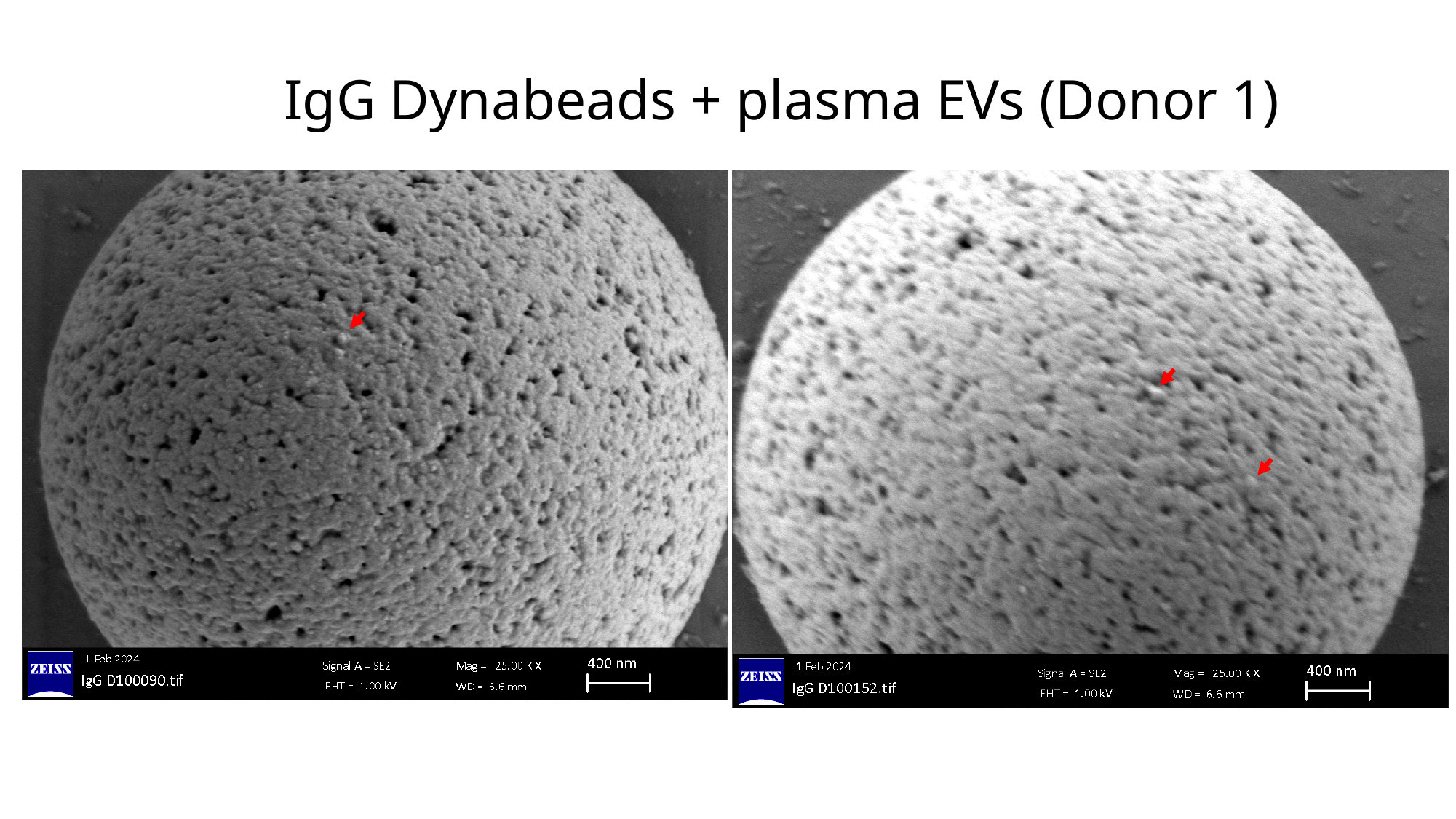

#### IgG Dynabeads + plasma EVs (Donor 1)

### Slide 11
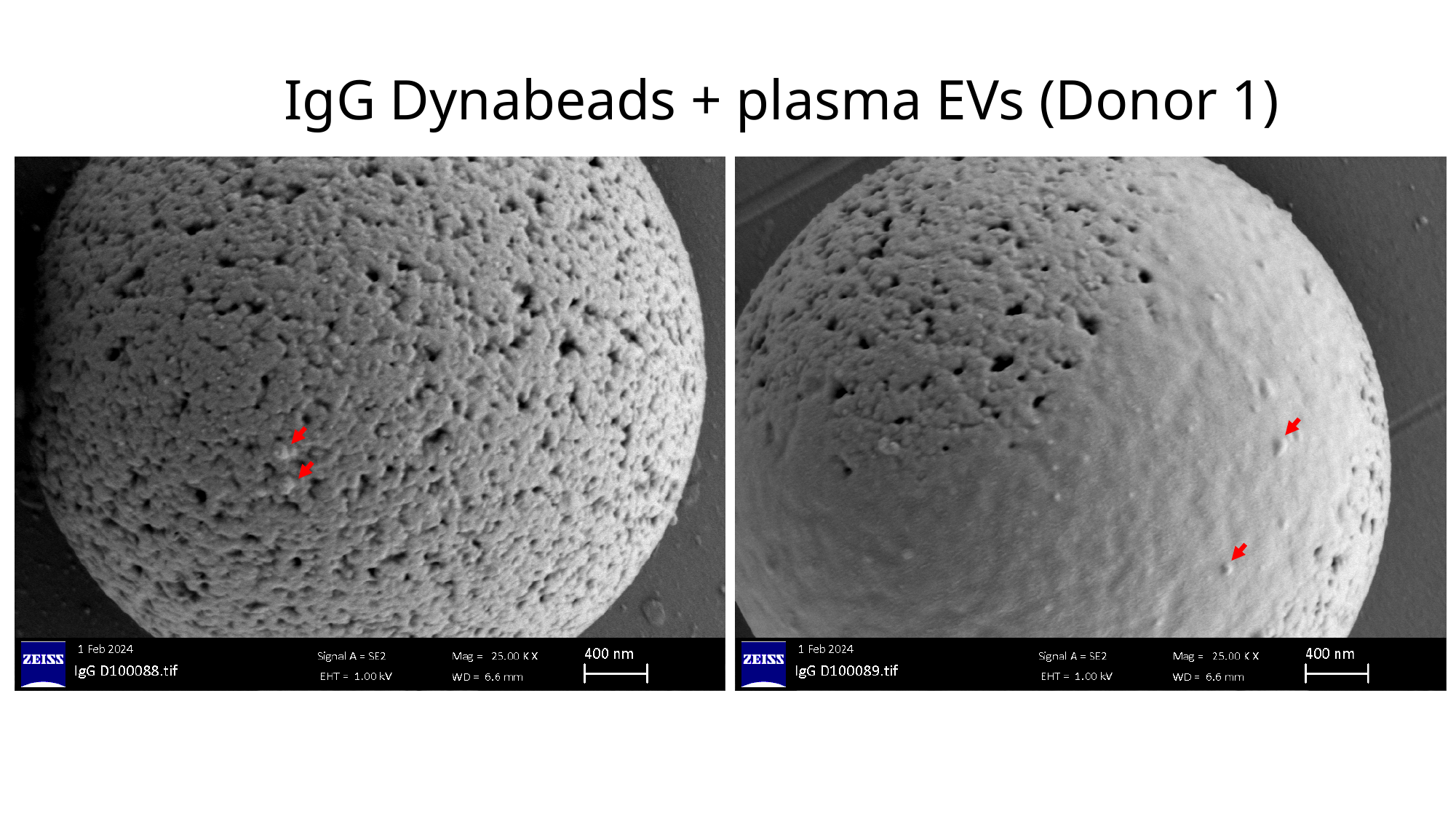

#### IgG Dynabeads + plasma EVs (Donor 1)

### Slide 12
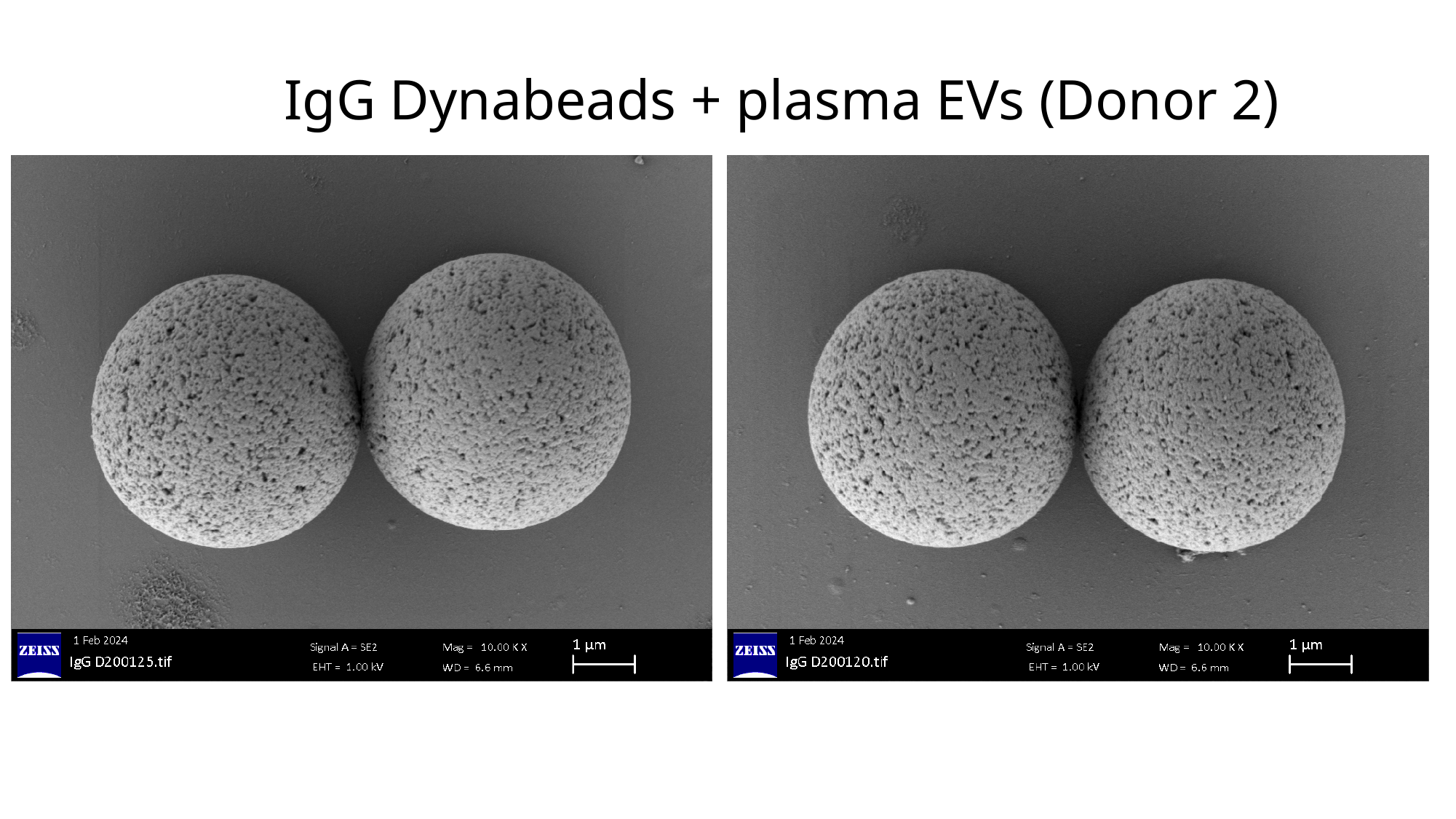

#### IgG Dynabeads + plasma EVs (Donor 2)

### Slide 13
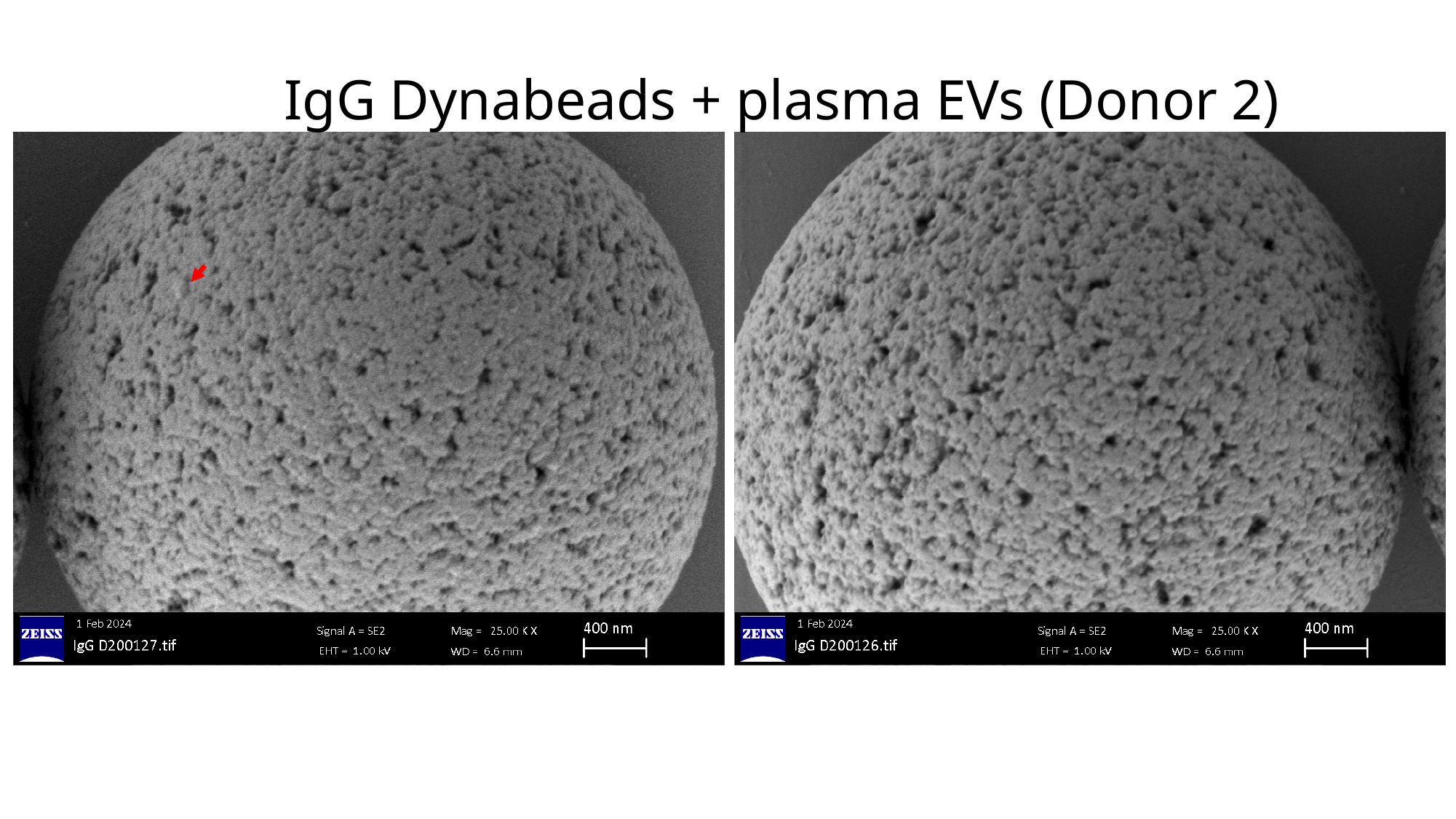

#### IgG Dynabeads + plasma EVs (Donor 2)

### Slide 14
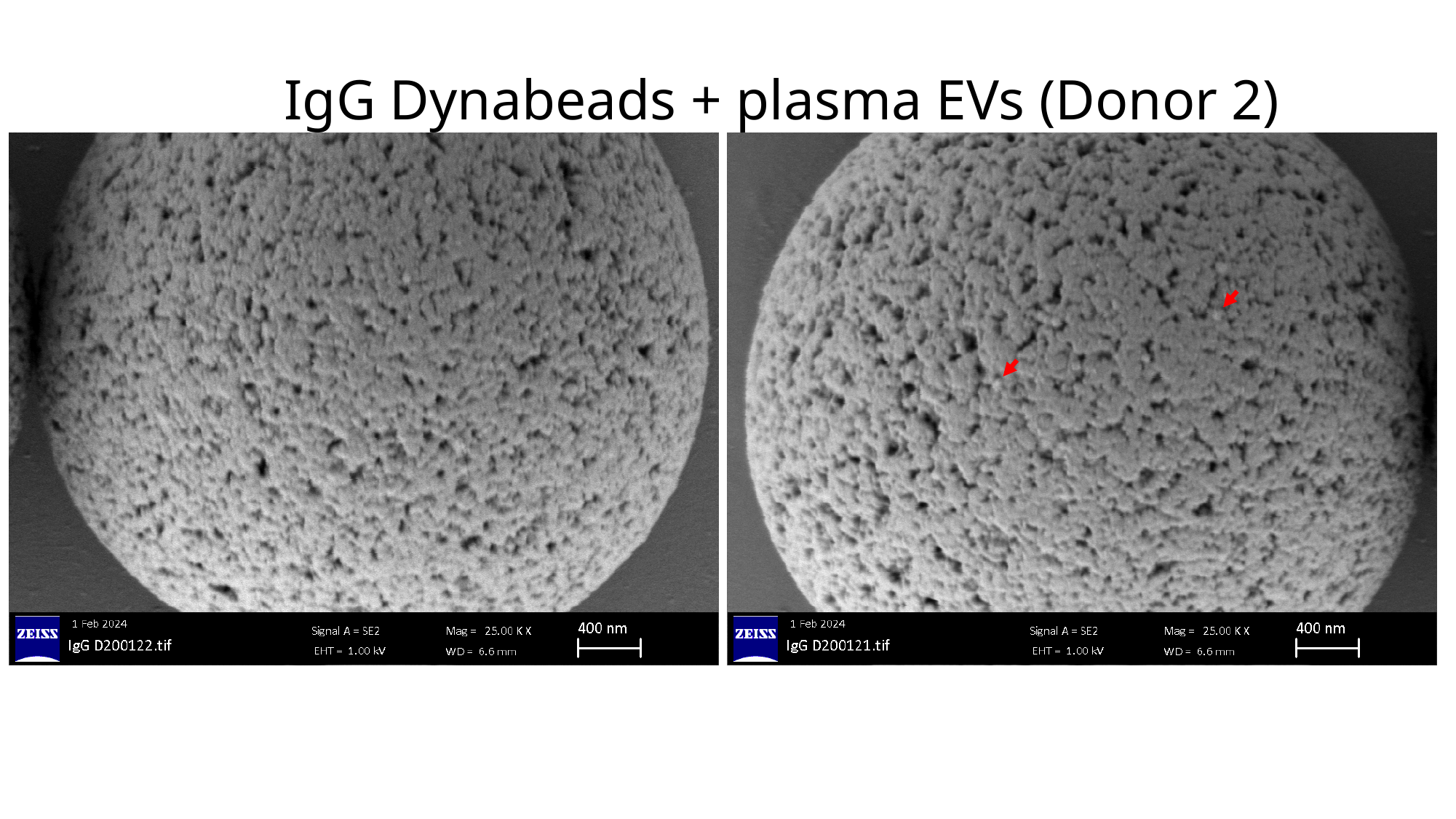

#### IgG Dynabeads + plasma EVs (Donor 2)

### Slide 15
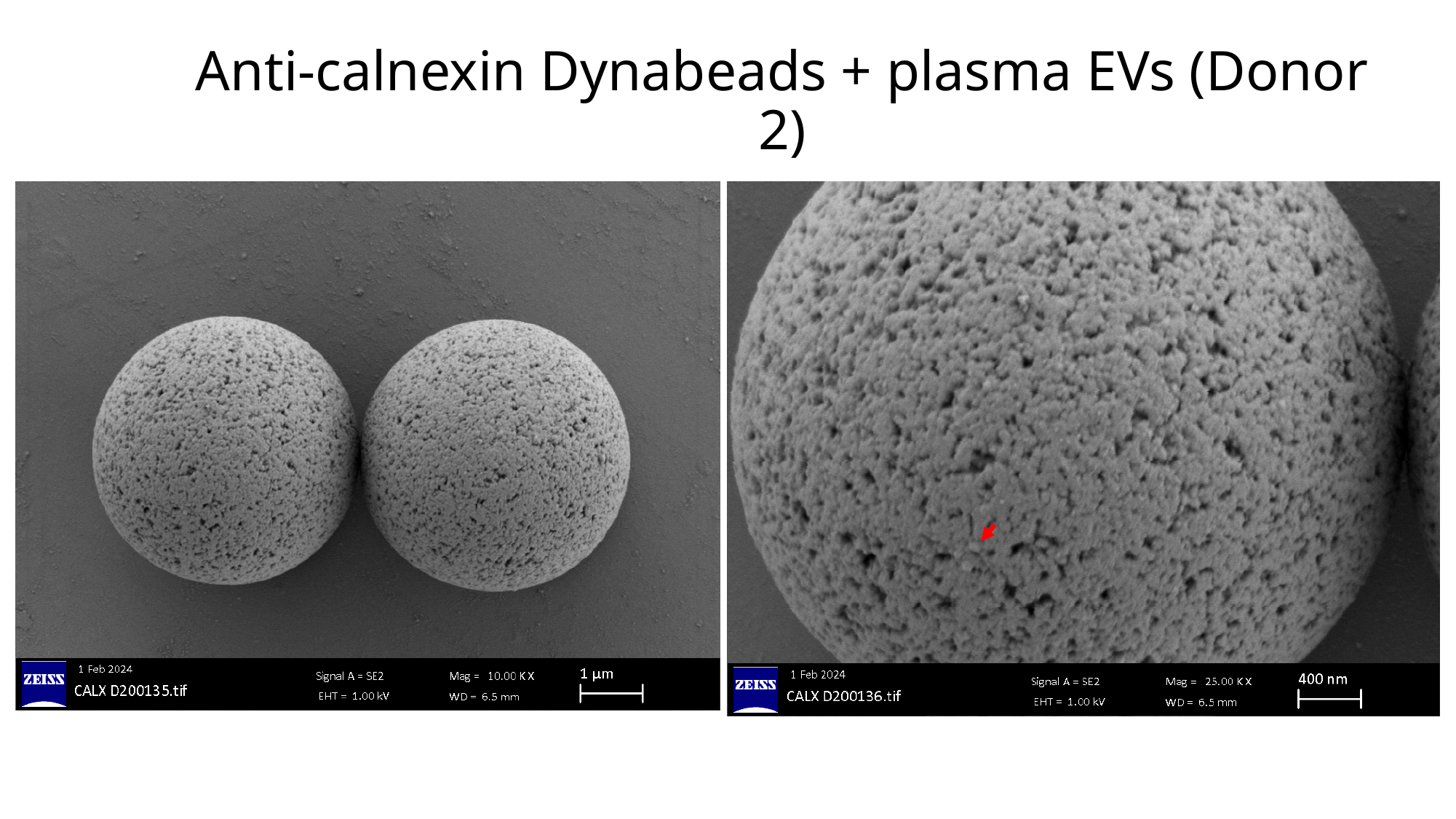

#### Anti-calnexin Dynabeads + plasma EVs (Donor 2)

### Slide 16
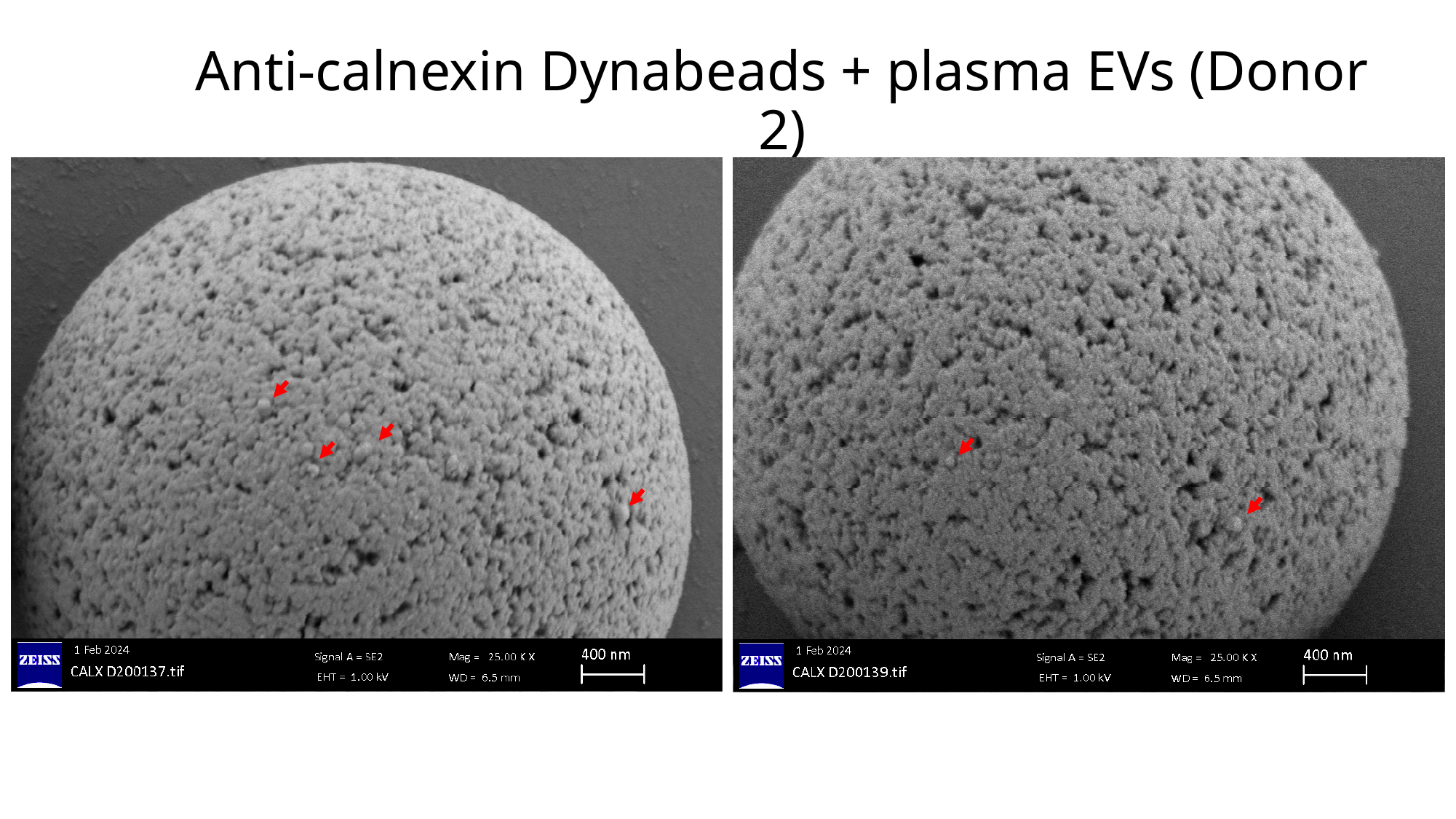

#### Anti-calnexin Dynabeads + plasma EVs (Donor 2)

### Slide 17
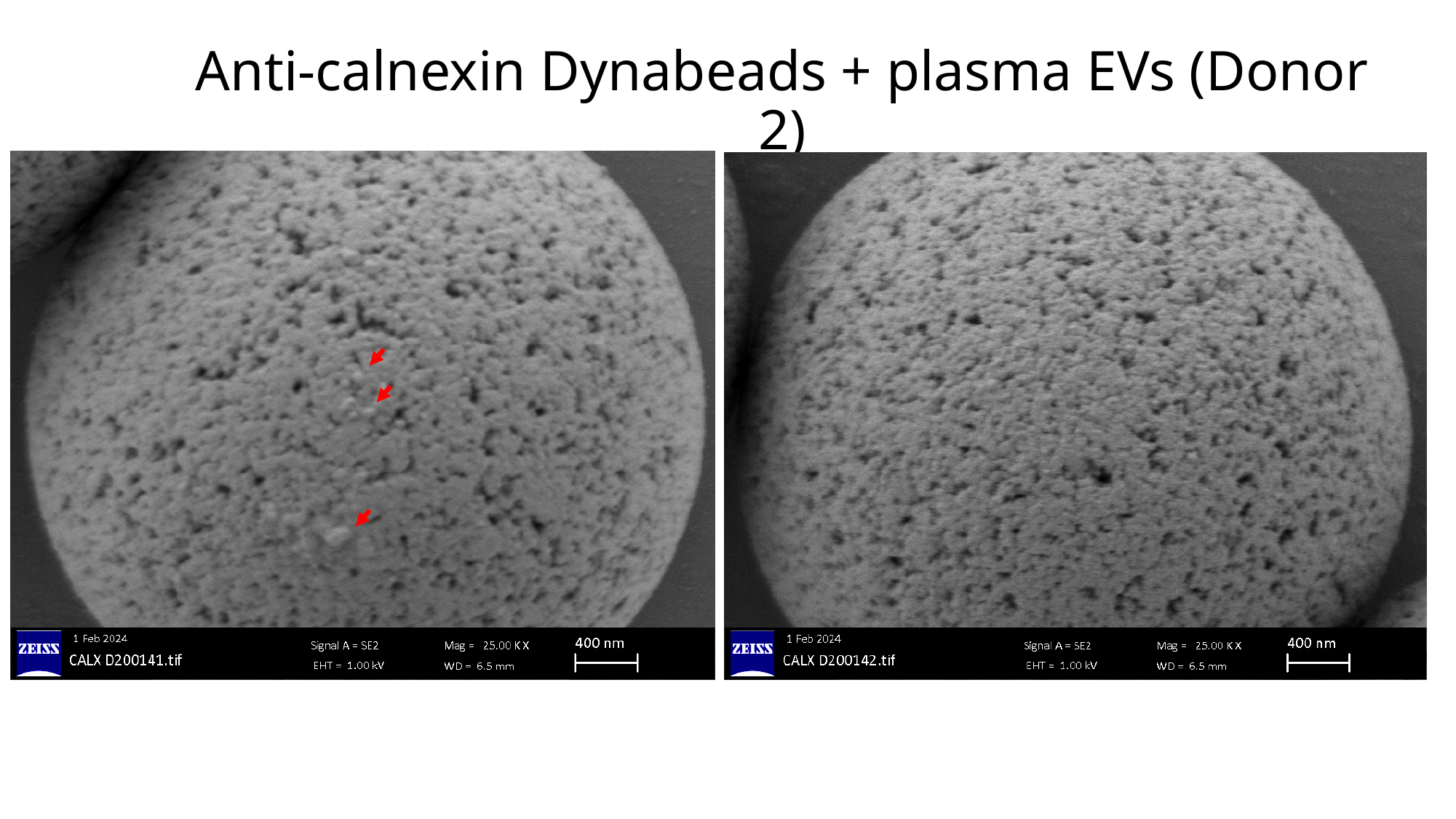

#### Anti-calnexin Dynabeads + plasma EVs (Donor 2)

### Slide 18
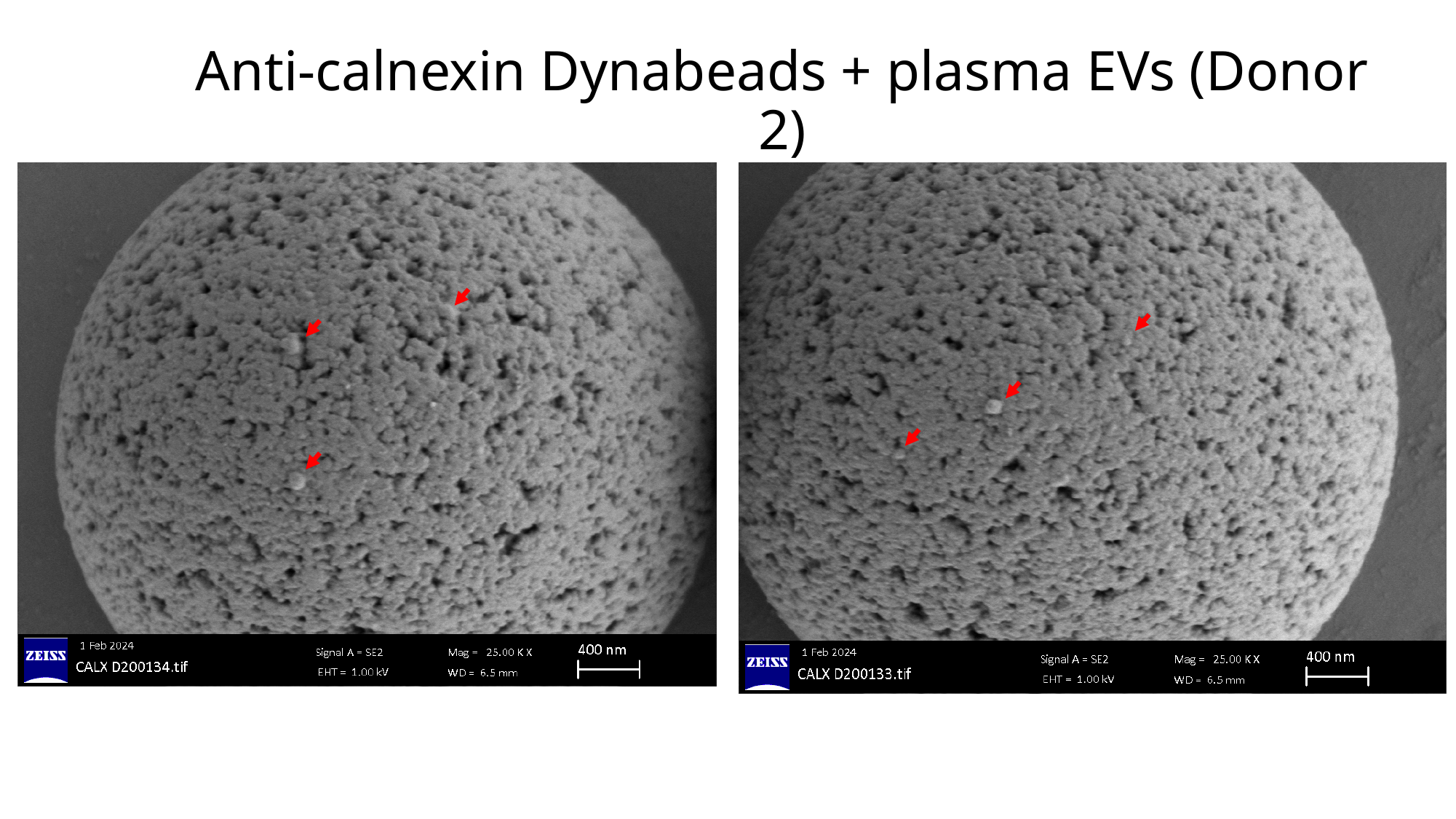

#### Anti-calnexin Dynabeads + plasma EVs (Donor 2)

### Slide 19
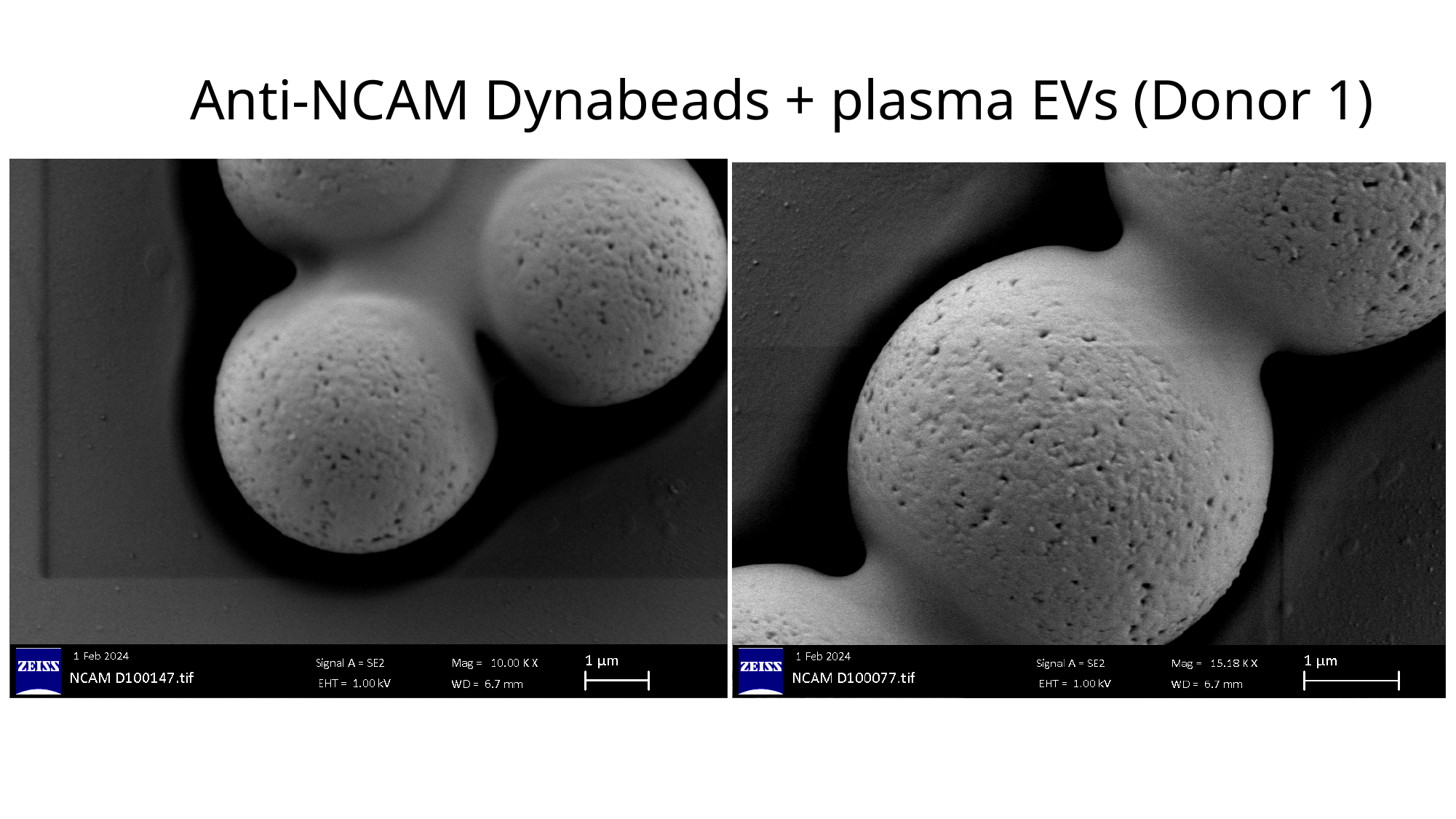

#### Anti-NCAM Dynabeads + plasma EVs (Donor 1)

### Slide 20
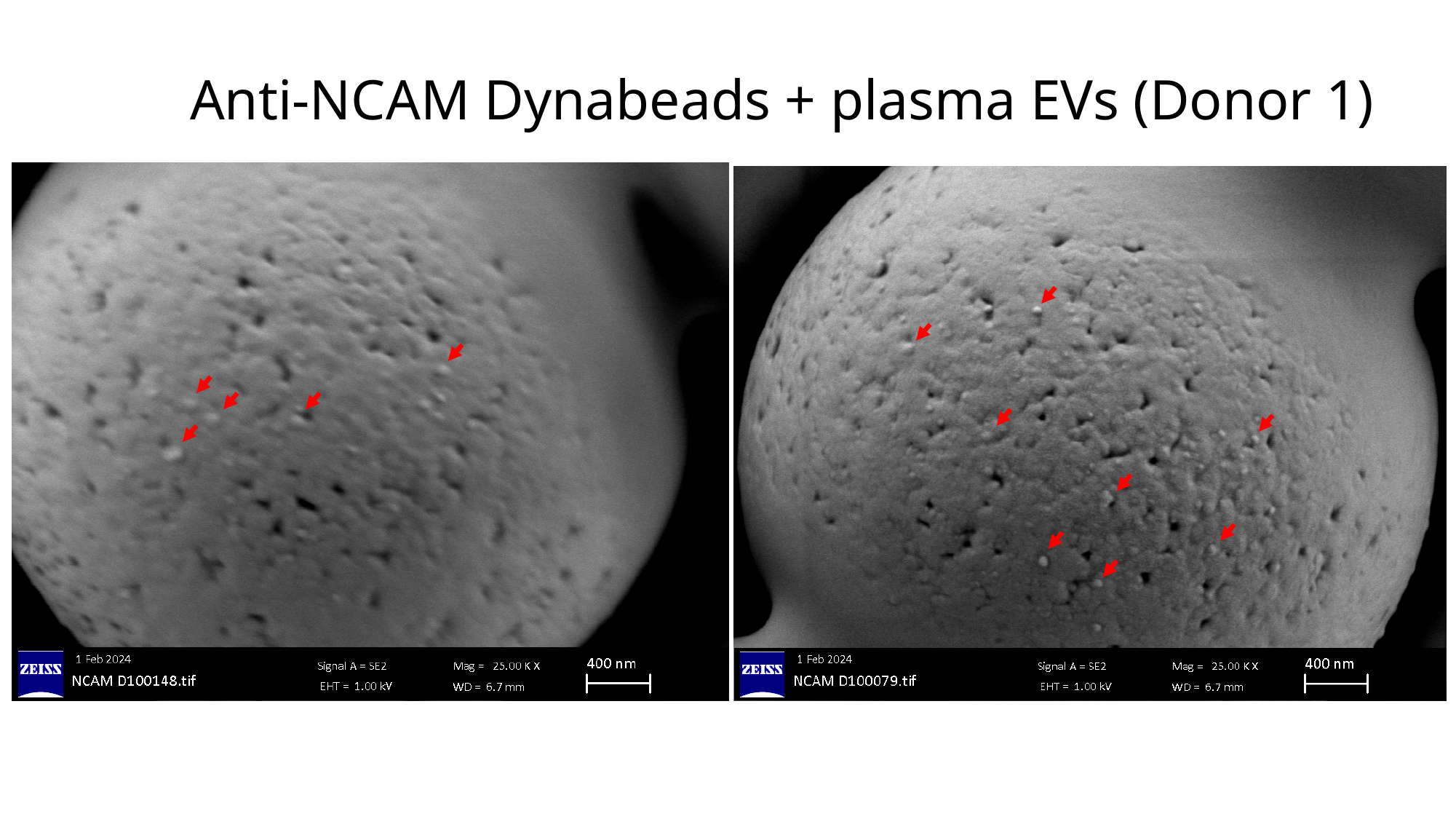

#### Anti-NCAM Dynabeads + plasma EVs (Donor 1)

### Slide 21
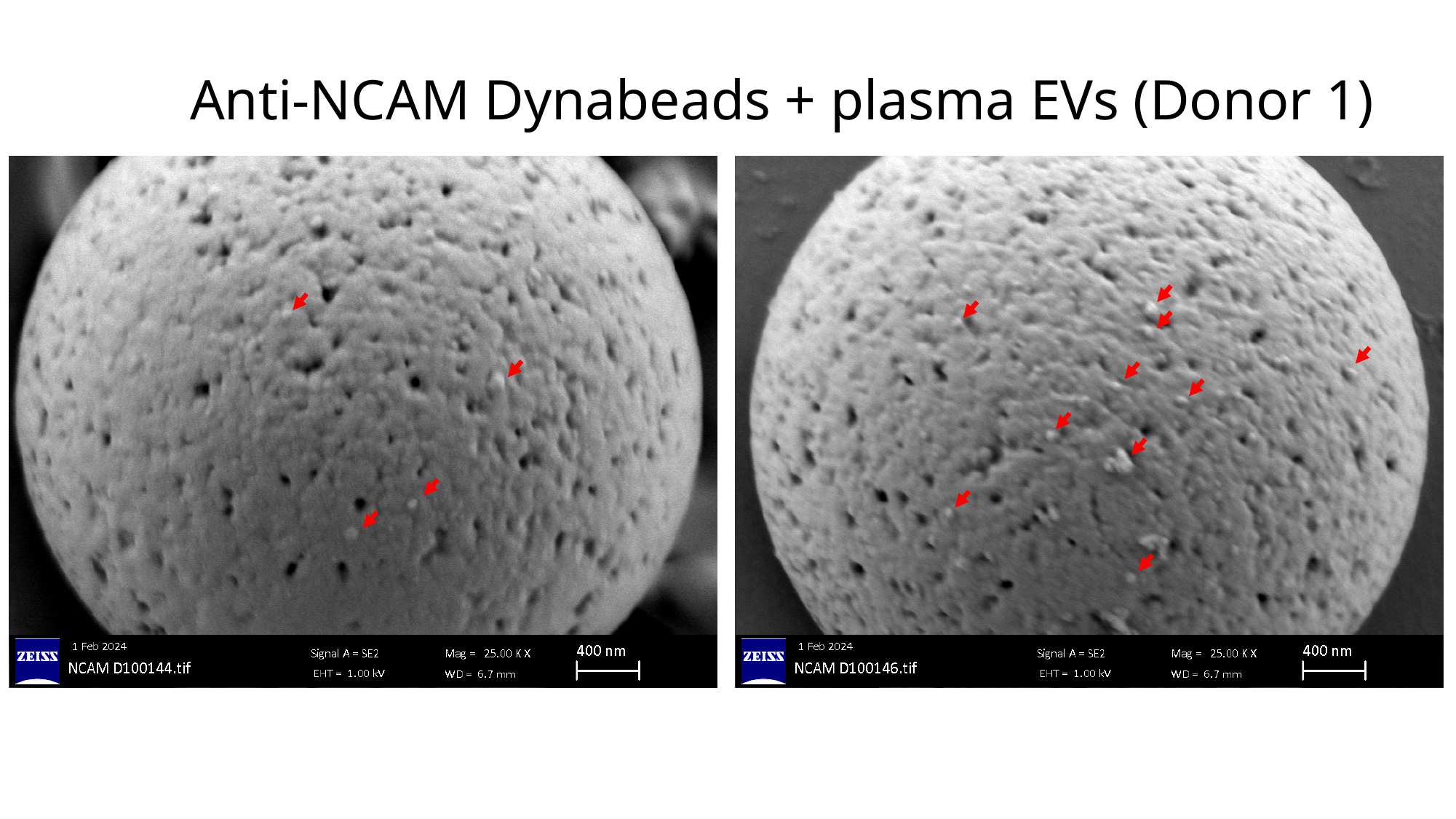

#### Anti-NCAM Dynabeads + plasma EVs (Donor 1)

### Slide 22
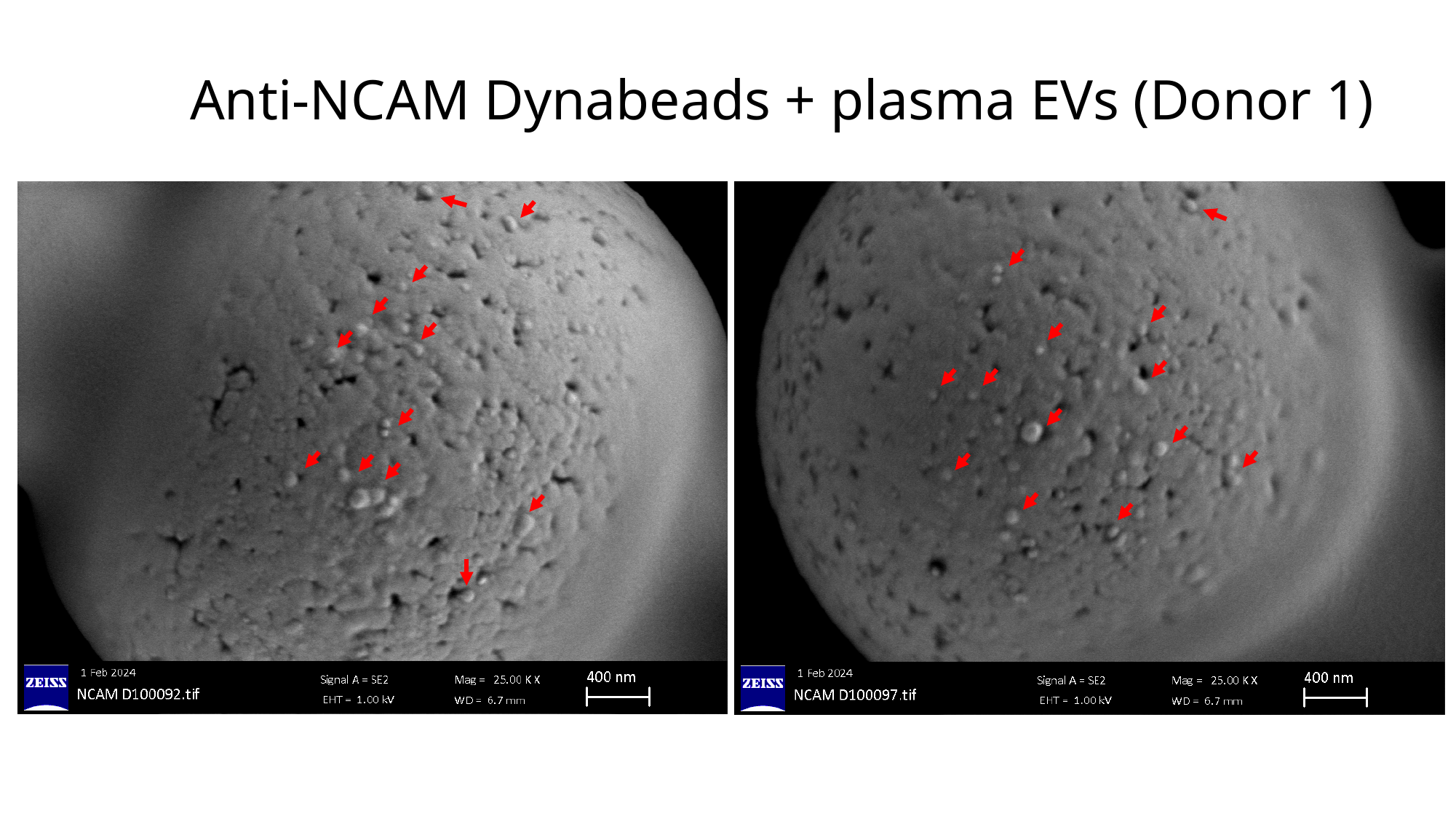

#### Anti-NCAM Dynabeads + plasma EVs (Donor 1)

### Slide 23
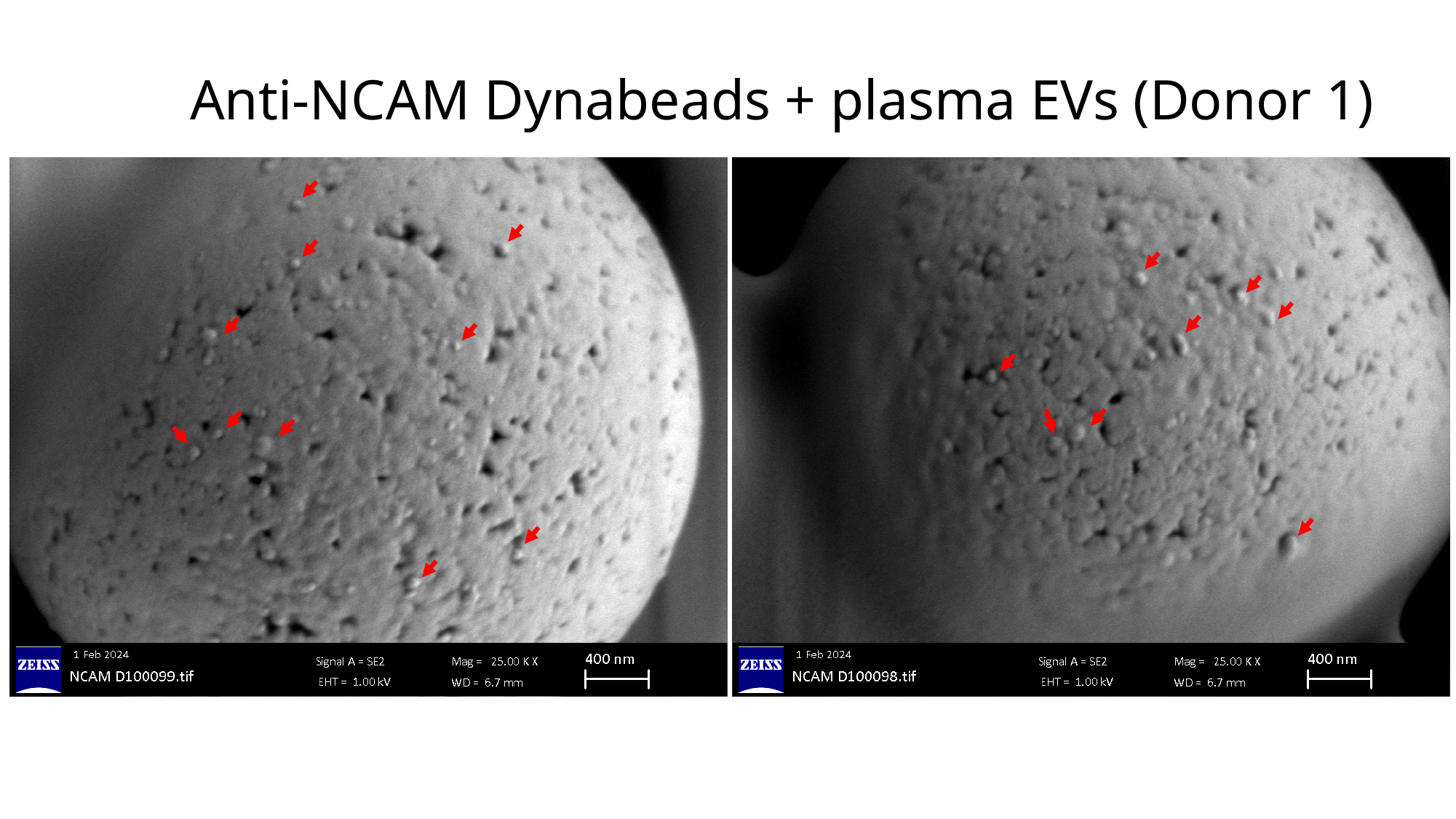

#### Anti-NCAM Dynabeads + plasma EVs (Donor 1)

### Slide 24
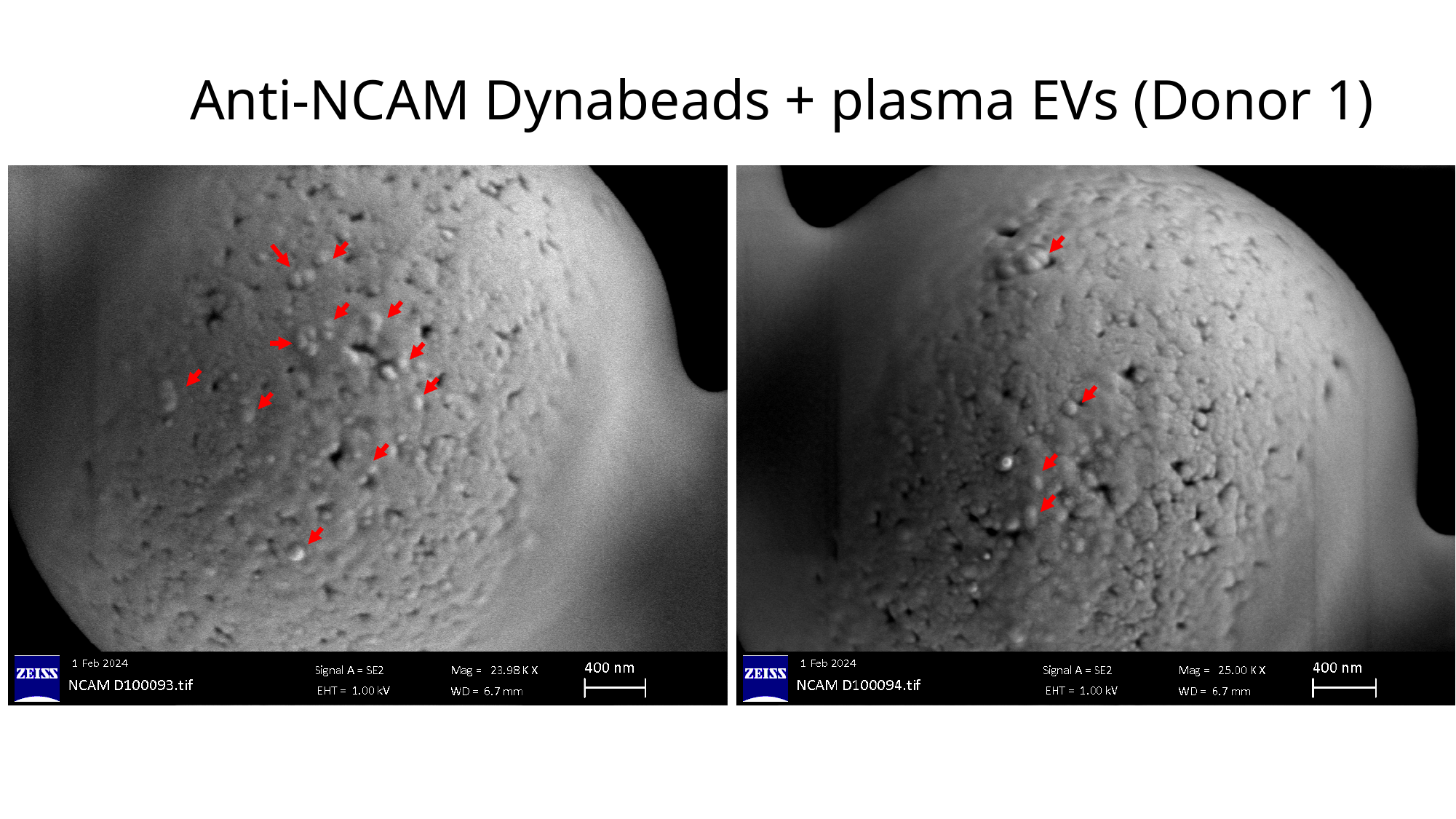

#### Anti-NCAM Dynabeads + plasma EVs (Donor 1)

### Slide 25
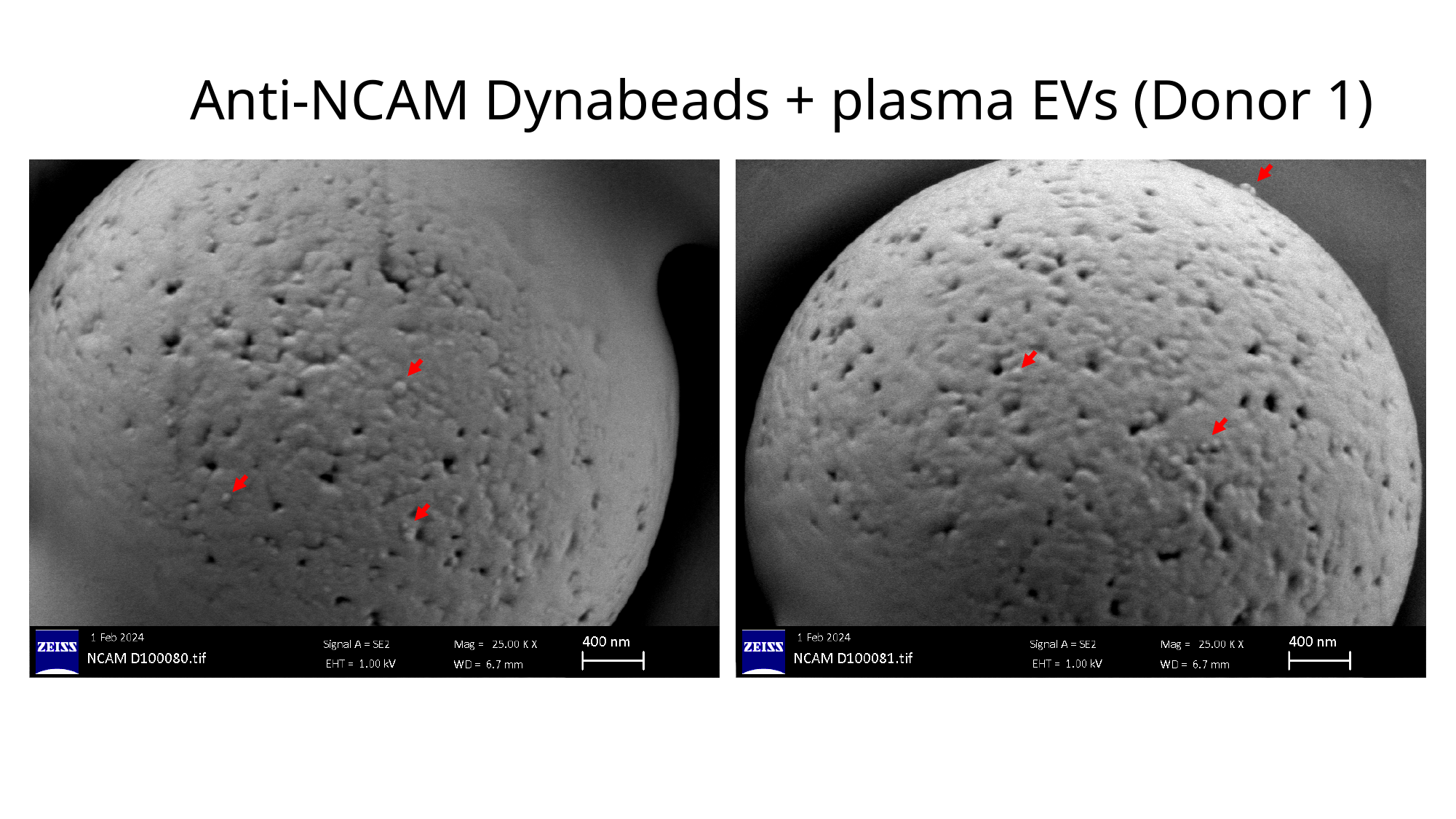

#### Anti-NCAM Dynabeads + plasma EVs (Donor 1)

### Slide 26
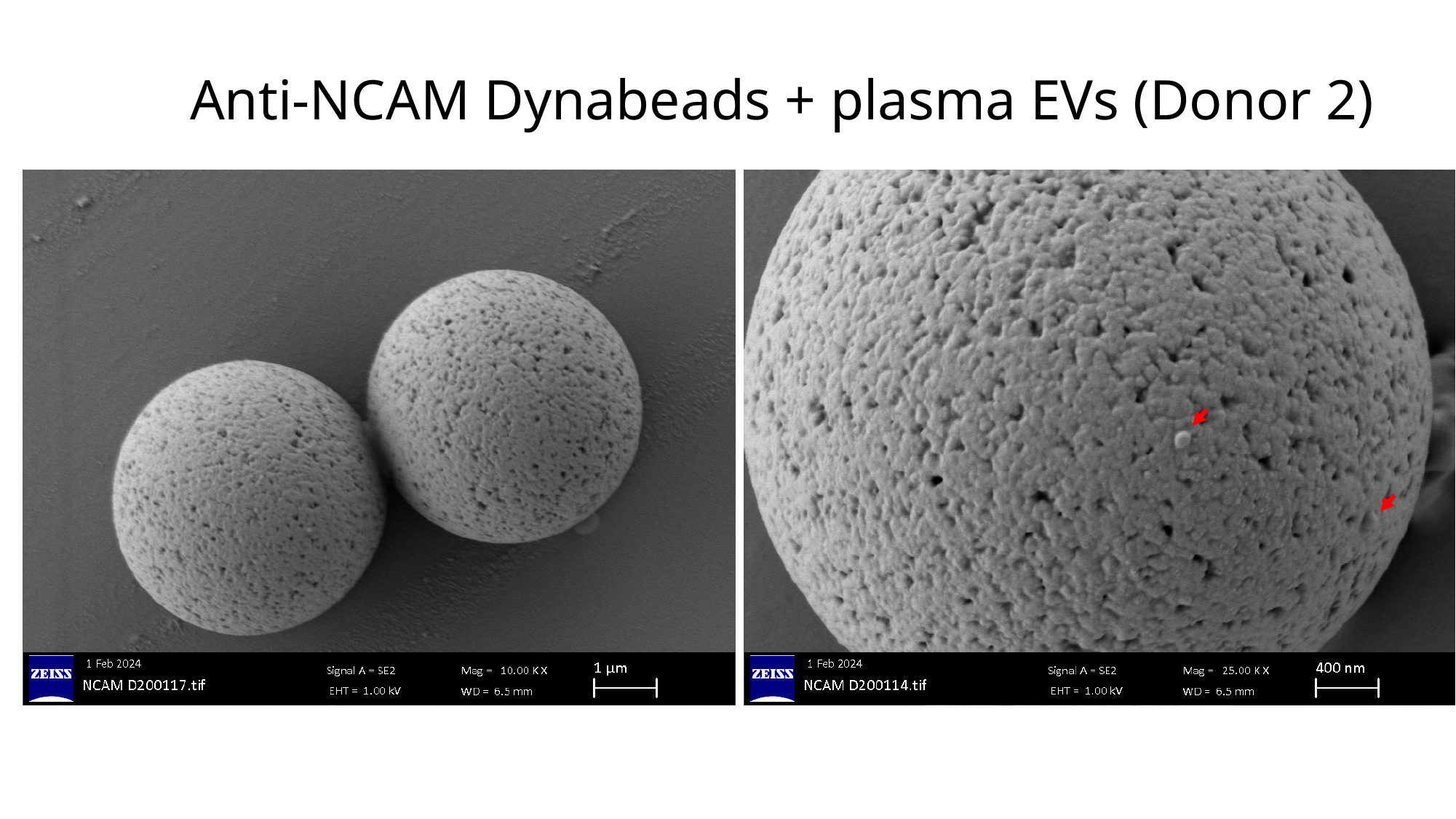

#### Anti-NCAM Dynabeads + plasma EVs (Donor 2)

### Slide 27
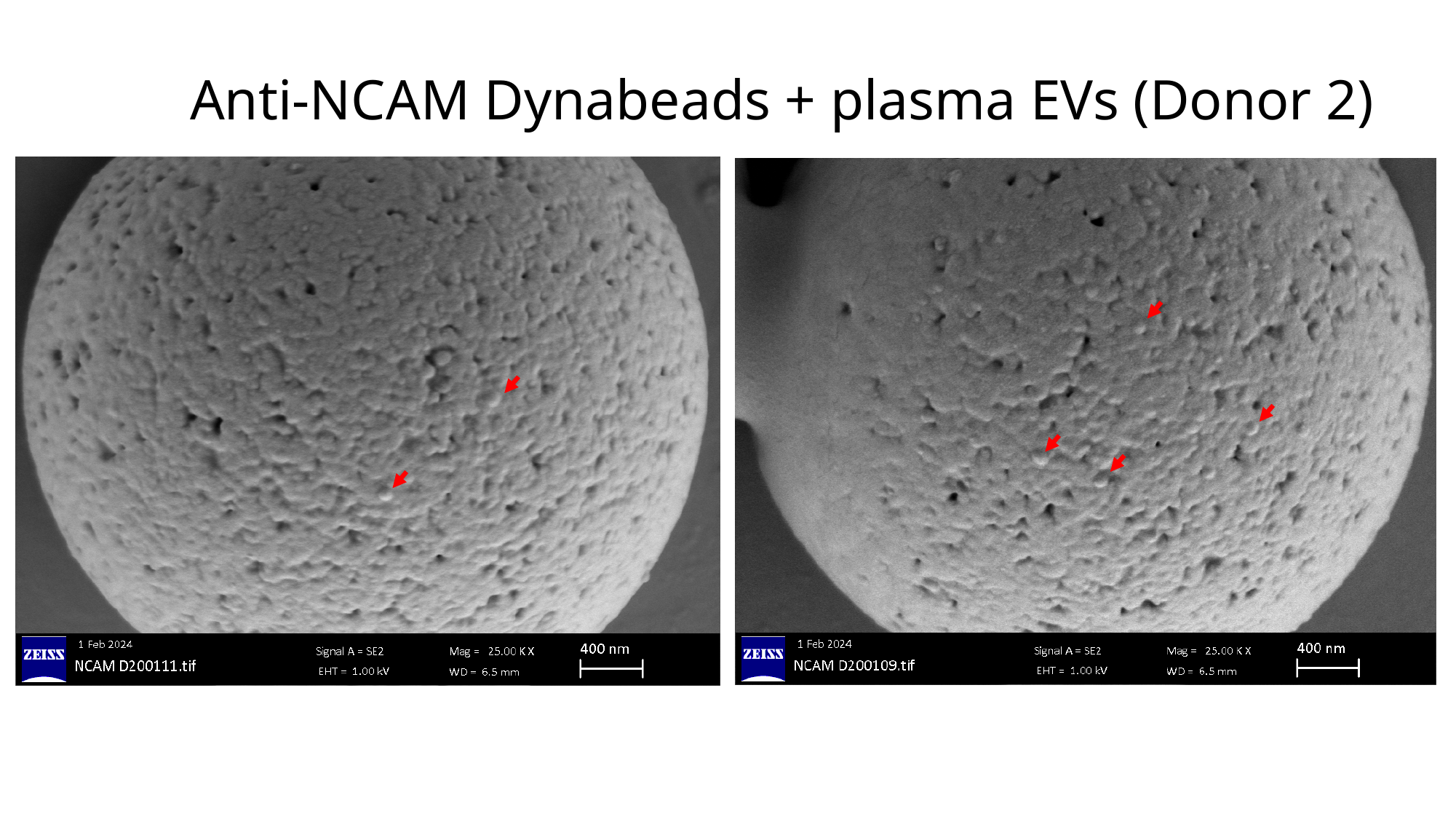

#### Anti-NCAM Dynabeads + plasma EVs (Donor 2)

### Slide 28
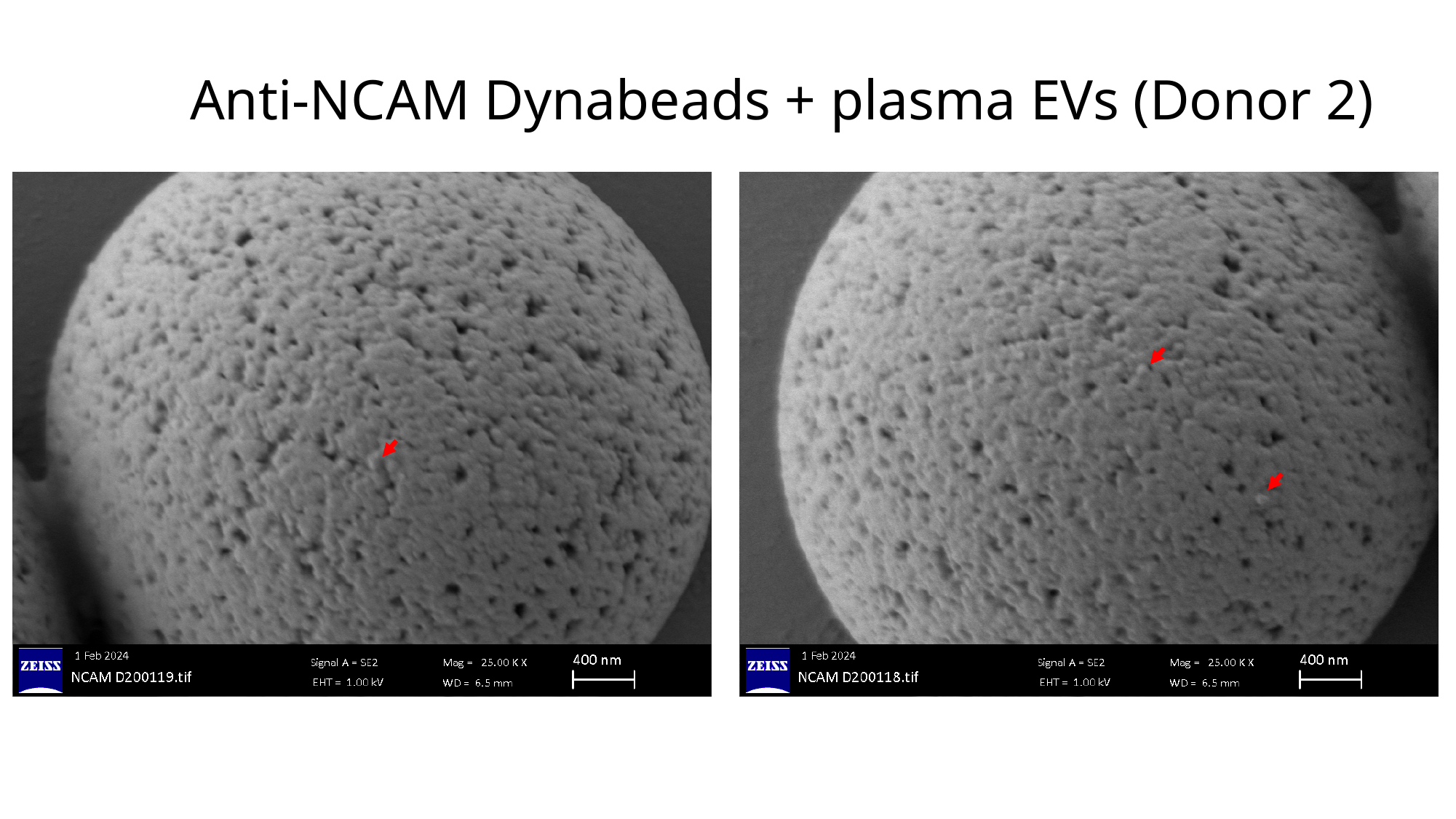

#### Anti-NCAM Dynabeads + plasma EVs (Donor 2)

### Slide 29
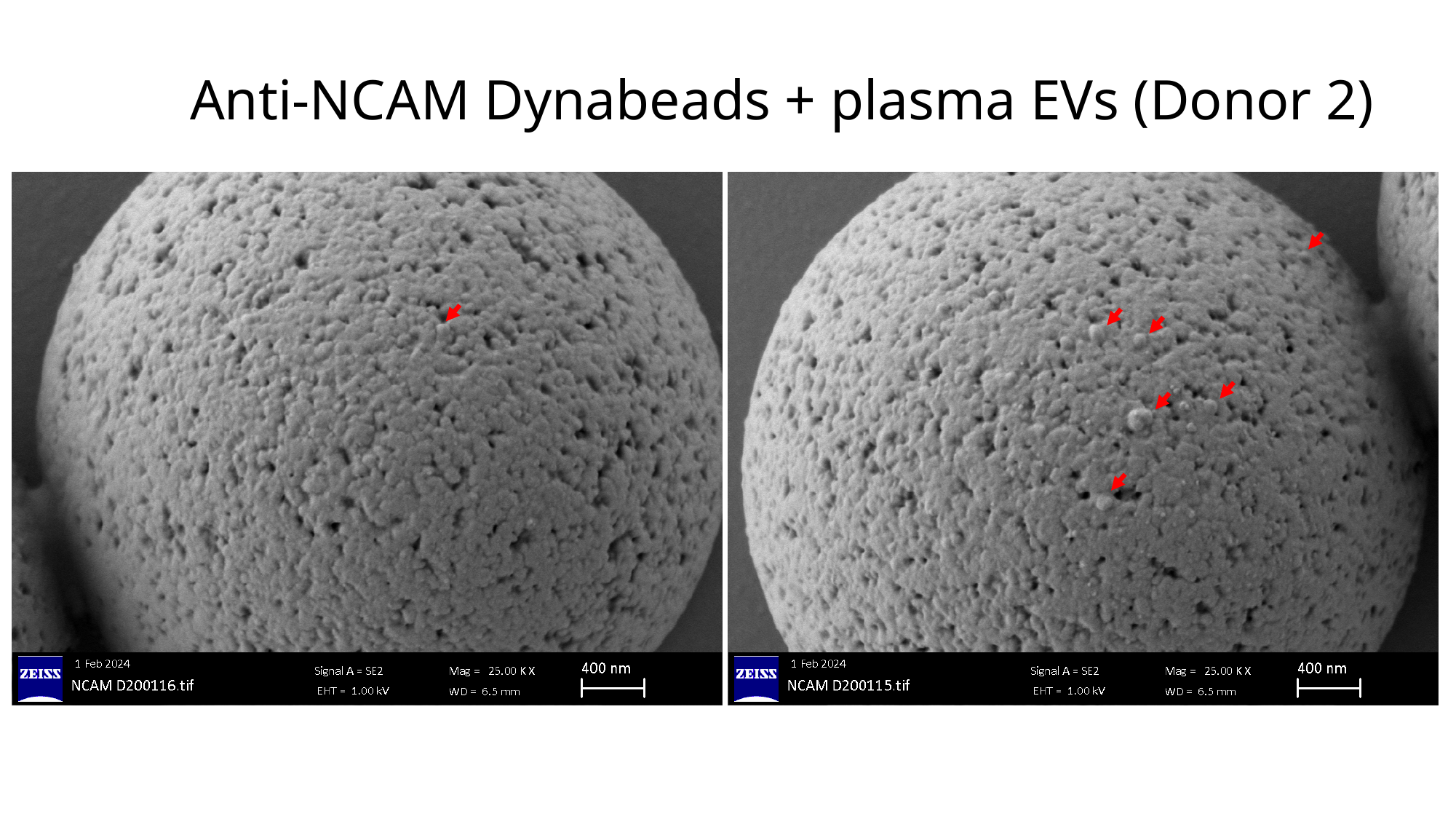

#### Anti-NCAM Dynabeads + plasma EVs (Donor 2)

### Slide 30
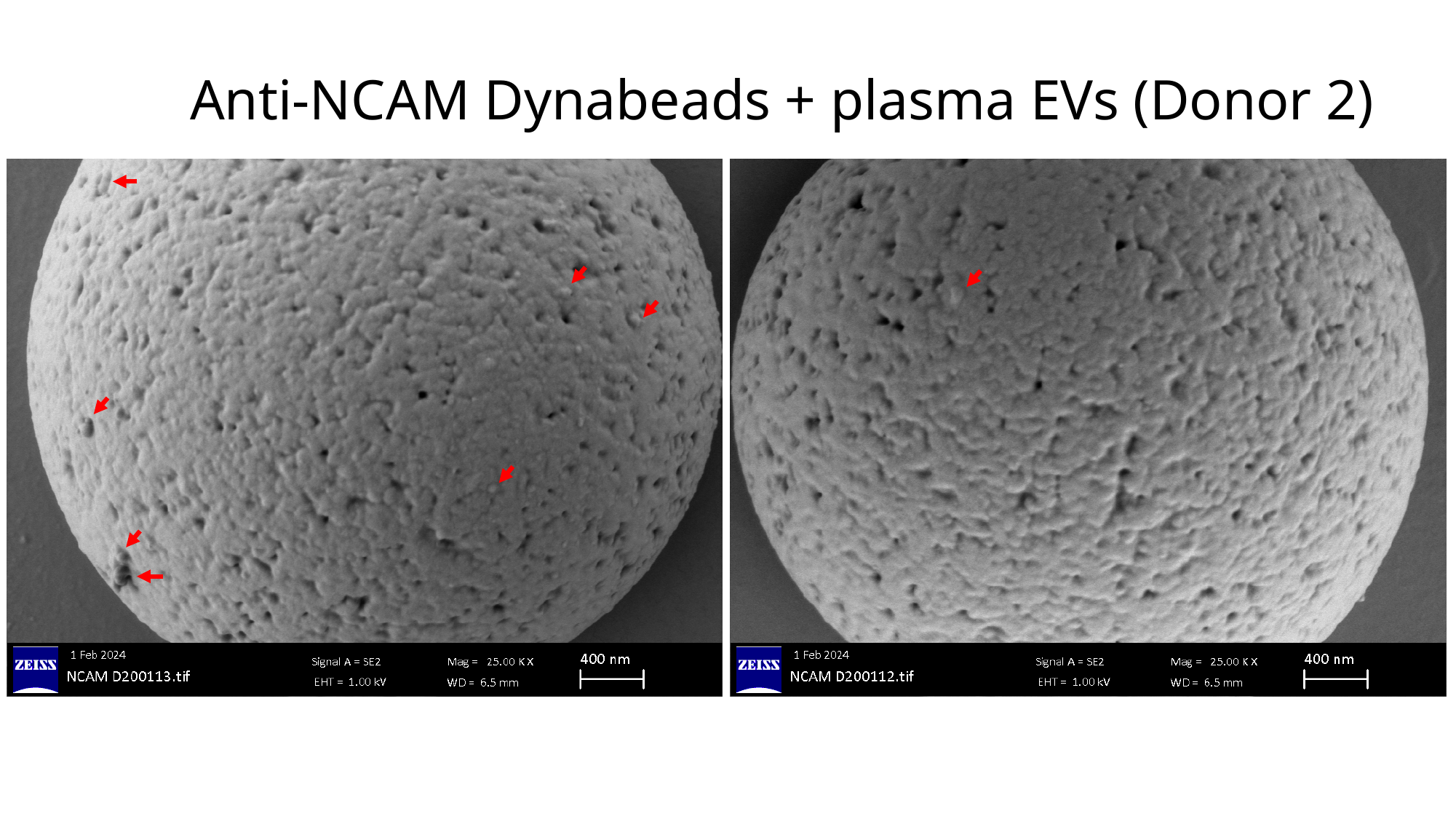

#### Anti-NCAM Dynabeads + plasma EVs (Donor 2)

### Slide 31
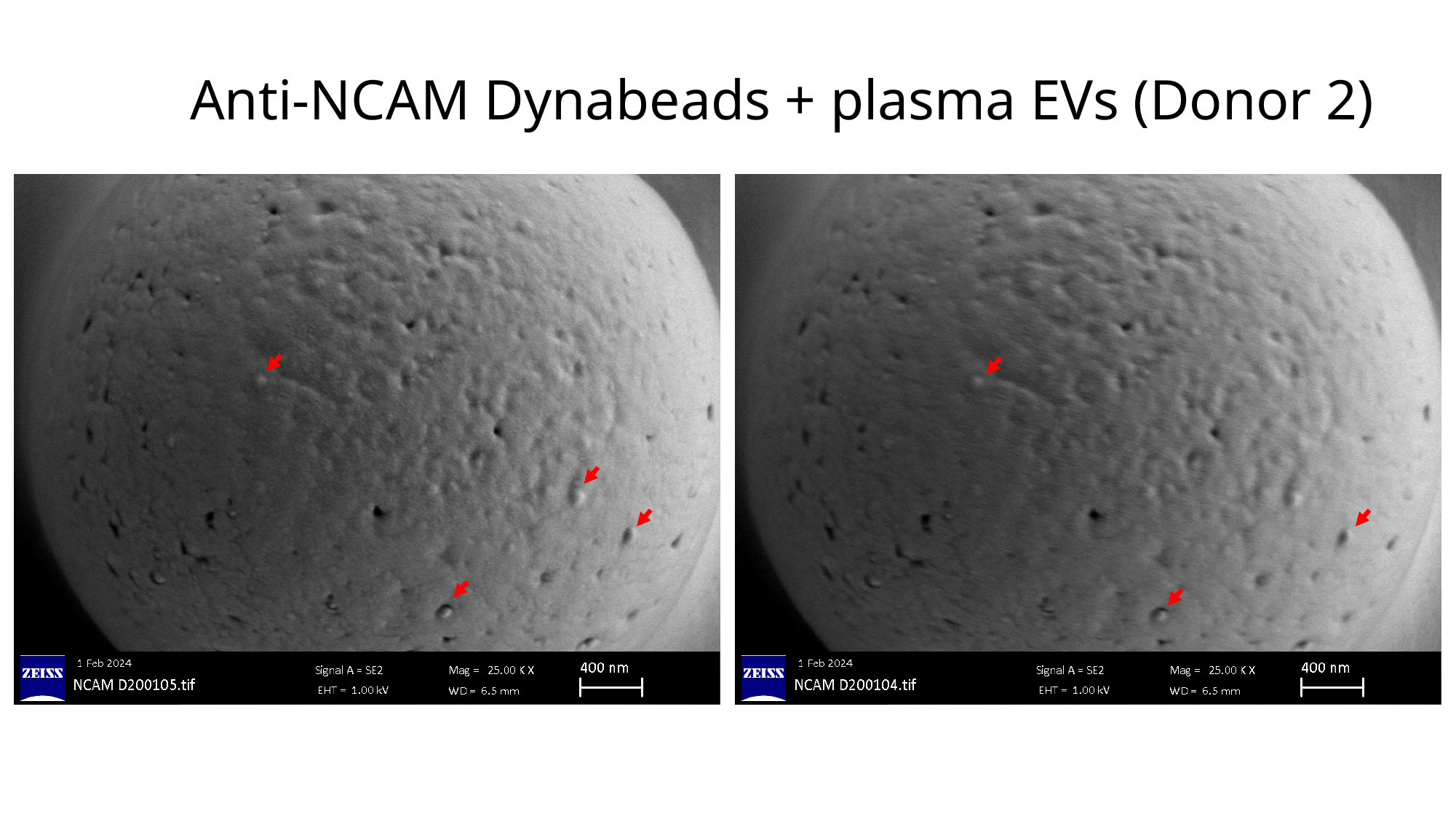

#### Anti-NCAM Dynabeads + plasma EVs (Donor 2)
